# Sensitivity-enhanced magnetic resonance reveals hydrogen intermediates during active [Fe]-hydrogenase catalysis

**DOI:** 10.1101/2023.05.10.540199

**Authors:** Lukas Kaltschnee, Andrey N. Pravdivtsev, Manuel Gehl, Gangfeng Huang, Georgi L. Stoychev, Christoph Riplinger, Maximilian Keitel, Frank Neese, Jan-Bernd Hövener, Alexander A. Auer, Christian Griesinger, Seigo Shima, Stefan Glöggler

## Abstract

Molecular hydrogen (H_2_) is considered an eco-friendly future energy-carrier and an alternative to fossil fuel^1^ and thus, major efforts are directed towards identifying efficient and economical hydrogen catalysts.^2,3^ Efficient hydrogen catalysis is used by many microorganisms, some of them producing H_2_ from organic materials and others consuming it.^4-6^ To metabolize H_2_, these microorganisms use enzymes called hydrogenases.^7,8^ For the future development of efficient catalysts a detailed analysis of the catalytic mechanisms of such hydrogenases is required and existing analytical techniques could not provide a full understanding.^9^ Consequently, new analytical technologies are of utmost importance to unravel natures’ blueprints for highly efficient hydrogen catalysts. Here, we introduce signal-enhanced or hyperpolarized, nuclear magnetic resonance (NMR) to study hydrogenases under turnover conditions. So far undiscovered hydrogen species of the catalytic cycle of [Fe]-hydrogenases, are revealed and thus, extend the knowledge regarding this class of enzymes. These findings pave new pathways for the exploration of novel hydrogen metabolisms *in vivo*. We furthermore envision that the results contribute to the rational design of future catalysts to solve energy challenges of our society.

## Main article

Hydrogenases are grouped into three different evolutionary types, i.e. [NiFe]-, [FeFe]- and [Fe]- hydrogenases.^7^ While intermediates in the catalytic cycle of [NiFe]- and [FeFe]- hydrogenases have been studied by using super-high-resolution X-ray diffraction,^10^ nuclear resonance vibrational spectroscopy (NRVS)^11,12^, EPR and nuclear magnetic resonance spectroscopy (NMR)^13-15^, intermediates of the catalytic cycle of [Fe]-hydrogenases were thus far undetectable. [Fe]-hydrogenases contain a single iron atom at the active site that as proposed, maintains the diamagnetic Fe^+II^-state throughout the catalytic cycle.^16^ The iron is complexed in an iron-guanylyl pyridinol (FeGP) cofactor (Fig. 1A, purple).^16-18^ Upon binding of the substrate methenyl-tetrahydromethanopterin (methenyl-H_4_MPT^+^, also CH≡H_4_MPT^+^) (Fig. 1A, green), the protein changes from an open to a closed conformation, thereby bringing together FeGP and CH≡H_4_MPT^+^ to form the active site (Fig. 1A). In the active site, H_2_ is heterolytically cleaved (H_2_ *⇄* H^+^ + H^-^) and the hydride (H^-^) is stereo-specifically transferred to the *pro-R* position of the methylene carbon (C_14a_) of methylene-H_4_MPT (CH_2_=H_4_MPT) (Fig. 1B).^19^ Due to this heterolytic reaction, no paramagnetic species occur and thus, [Fe]-hydrogenase, which is also referred to as H_2_-forming methylene-H_4_MPT dehydrogenase (Hmd). Hmd catalyzes isotope exchange between water and dissolved hydrogen, where both single and double-isotope exchange can take place in one binding event (Fig. 1C).^20^ Computational models suggest multiple iron hydrogen species along the Hmd catalytic cycle.^16,21,22^ However, these species have not yet been characterized experimentally.^16-19,23-26^

**Fig. 1.**
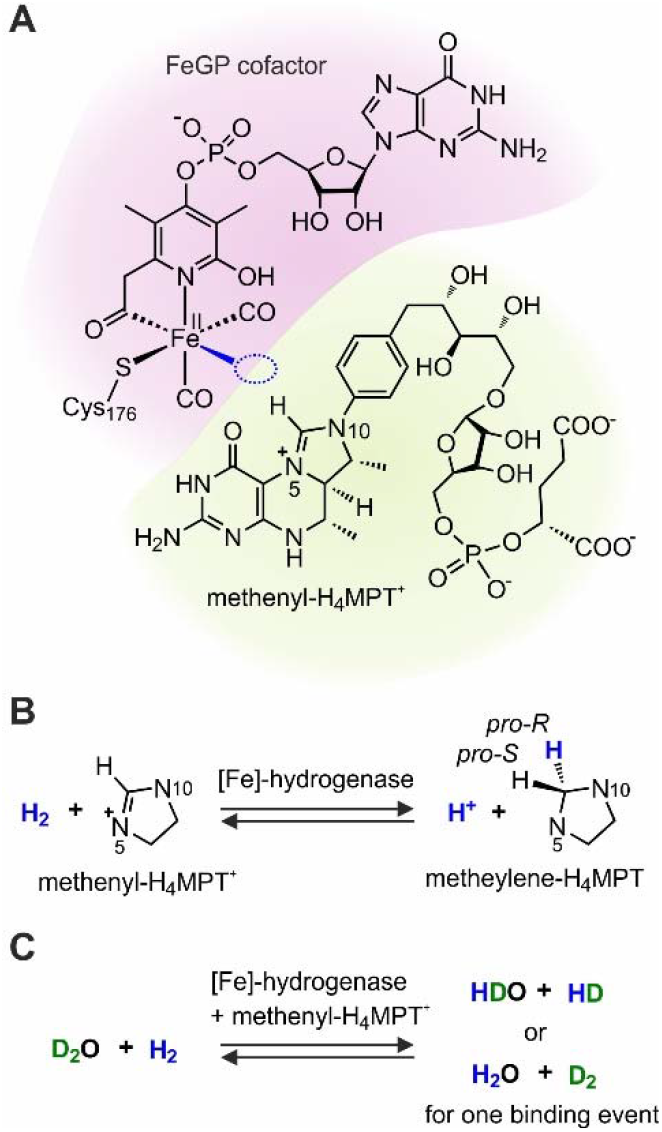
Active site of Hmd and reactions catalyzed by Hmd. (**A**) FeGP cofactor and methenyl-H_4_MPT^+^ substrate in the closed active site. The proposed H_2_ binding site is highlighted in blue. (**B & C**) Reactions catalyzed by Hmd.^19,20,36^ For methenyl-H_4_MPT^+^ and methylene-H_4_MPT, only the imidazolinium and imidazolidine rings are depicted.

To address and overcome this challenge, we pursued metalloprotein studies using sensitivity-enhanced NMR based on parahydrogen (*p*-H_2_) – H_2_ that is enriched in its asymmetric nuclear spin state, i.e. the *para*-state – which creates **p**ara**h**ydrogen-**i**nduced **p**olarization (PHIP).^27,28^ This makes it possible to characterize transiently bound hydrogen species with strongly enhanced sensitivity and as such represents a novel approach for the study of enzymes and proteins.^28, 29,30^.

We observed enhancements emerging from short periods of binding of hydrogen species (H_2_, H^-^, H^+^) at the Hmd catalytic site. During these binding events, the not observable nuclear spin order of *p*-H_2_ is partially converted into observable nuclear spin polarization, e.g. hyperpolarization, and we found that PHIP makes it possible to study H_2_ activation and conversion by Hmd. To characterize these transiently formed intermediates of the Hmd catalytic cycle, we combined PHIP experiments, kinetic modelling of NMR experiments, and quantum chemical computations of NMR parameters.

Specifically, we collected ^1^H-NMR signals that transiently appear after bubbling hydrogen to solutions of reconstituted Hmd holoenzyme from *Methanocaldococcus jannaschii* (jHmd) and its substrate (methenyl-H_4_MPT^+^) (SI 1.1-1.12, Fig. S1). For a comparison, these experiments were performed either with *p*-H_2_ or with “normal” hydrogen in room-temperature equilibrium (*n*-H_2_). As indicated by changes in signal shape and intensity, enhancement effects were only observed when *p*-H_2_ (Fig. 2A) compared to *n*-H_2_ was used (Fig. 2B). Enhancement was not observed in the absence of jHmd or methenyl-H_4_MPT^+^ (Fig. S5), or with the H14A-jHmd mutant (Fig. S9), which is about 100-fold less active than the wild type. Thus, creating enhancement effects requires the active enzyme substrate complex (SI 2.1, Figs. S5-S11, Tables S1-S3) and reports on bound hydrogen intermediates.

**Fig. 2.**
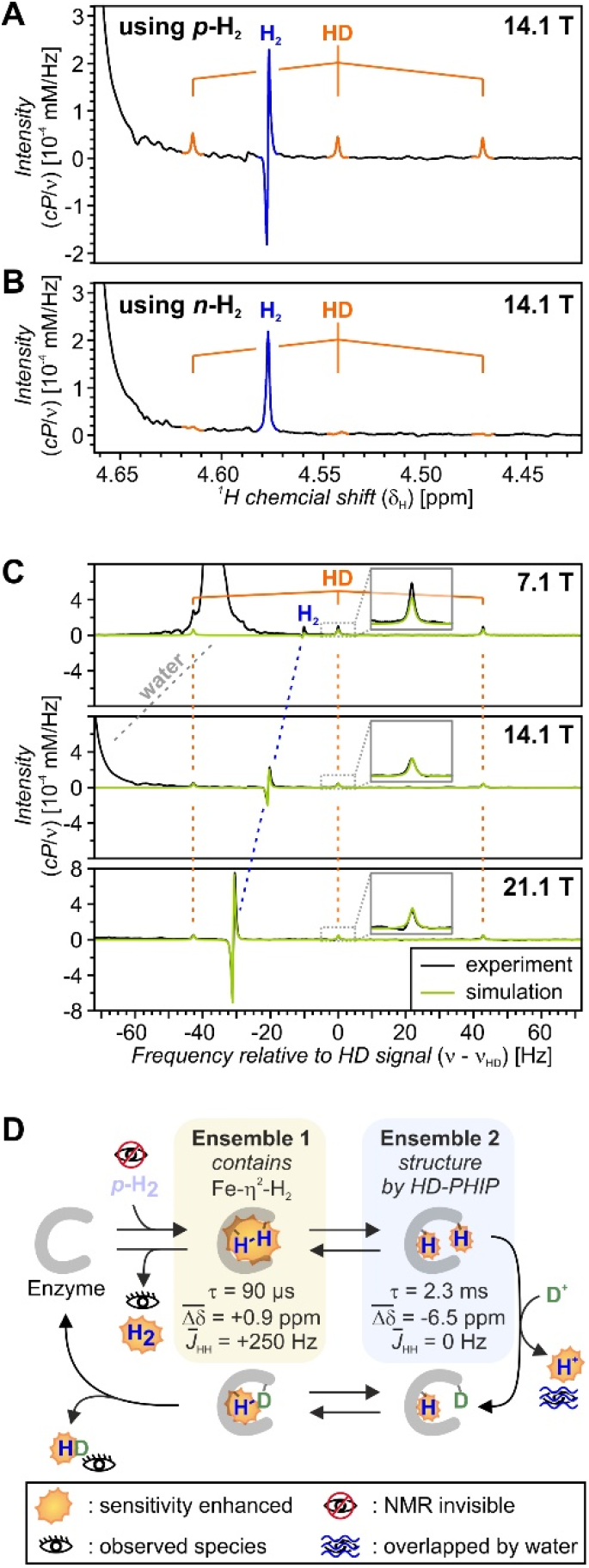
^1^H-PHIP effects observed in the presence of [Fe]-hydrogenase. (**A & B**) Dihydrogen regions of the ^1^H-NMR spectrum observed after bubbling either (**A**) *p*-H_2_ (87% enrichment) or (**B)** *n*-H_2_ through a sample containing [Fe]-hydrogenase, cofactors and substrate for 15 s (pulse sequence Fig. S1). Compare to Fig. S5. The sample was prepared from 1 μM jHmd and 3 μM [^13^C]-CH_2_=H_4_MPT in D_2_O-buffer (*p*D 6.0, 1 mM EDTA, 120 mM potassium phosphate). (**C**) Overlay of measured (black) and simulated (green) ^1^H-PHIP spectra for three *B*_0_ fields. Measured and simulated isotope-exchange kinetics are shown in Fig. S23. H_2;_ water peaks remain at the same chemical shifts (ppm) at all three *B*_0_ fields, however, a Hz-scale, relative to the HD-frequency, was chosen here for clearer visualization. Data are presented on a field-invariant absolute intensity scale, as detailed in SI 1.12. Sample conditions are in comparison to those used in panel A. Fig. S8 and Table S2 compare spectra for multiple samples. (**D**) Pictographic scheme of the model with two distinct bound state ensembles. The complete model is shown in detail in Fig. S15C, SI 2.8 and Tables S5 & S6. Selected parameters for the simulations in panel C are indicated in the figure.

Performing these studies in deuterated aqueous buffer made it possible to observe hyperpolarized H_2_ and HD, and both PHIP effects were sufficiently strong to be observed in a single scan at 1 μM jHmd concentration, which demonstrates the high sensitivity of PHIP.

The PHIP effects on H_2_ and HD originate from two different mechanisms that are both driven by coherent spin evolution. The H_2_-PHIP effect is assigned to a PNL effect (PNL for “partially negative line shape”), which explains the observed dispersion-like line shape specifically at high field (Fig. 2C).^31^ The mechanistic basis for this effect is the reversible binding of H_2_ to an asymmetric site as PNLs strongly depend upon the kinetics of this binding process. Numerical spectrum modeling (SI 2.6) for varying magnetic fields shows clearly that a time-averaged *J*-coupling of 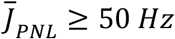 is required for the bound state (Fig. 2C, SI 2.8, Figs. S19-S22). This is compatible with a bound intermediate with an H-H distance below 1.5 Å that participates in the ensemble of bound states causing the PNL.^32^ Structural modeling, based upon the extracted experimental data, proves an iron dihydrogen species with side-on Fe-*η*2-H_2_ binding. The lifetime of this bound intermediate must be in the range of *τ*_*PNL*_ *1 μs - 100 μs* (SI 2.8, Fig. S21B, Table S6), and rapid exchange with dissolved, free H_2_ is required to explain the spectra. This exchange is assigned to interconversion between structures **1 & 2** (Fig. 3).

**Fig. 3.**
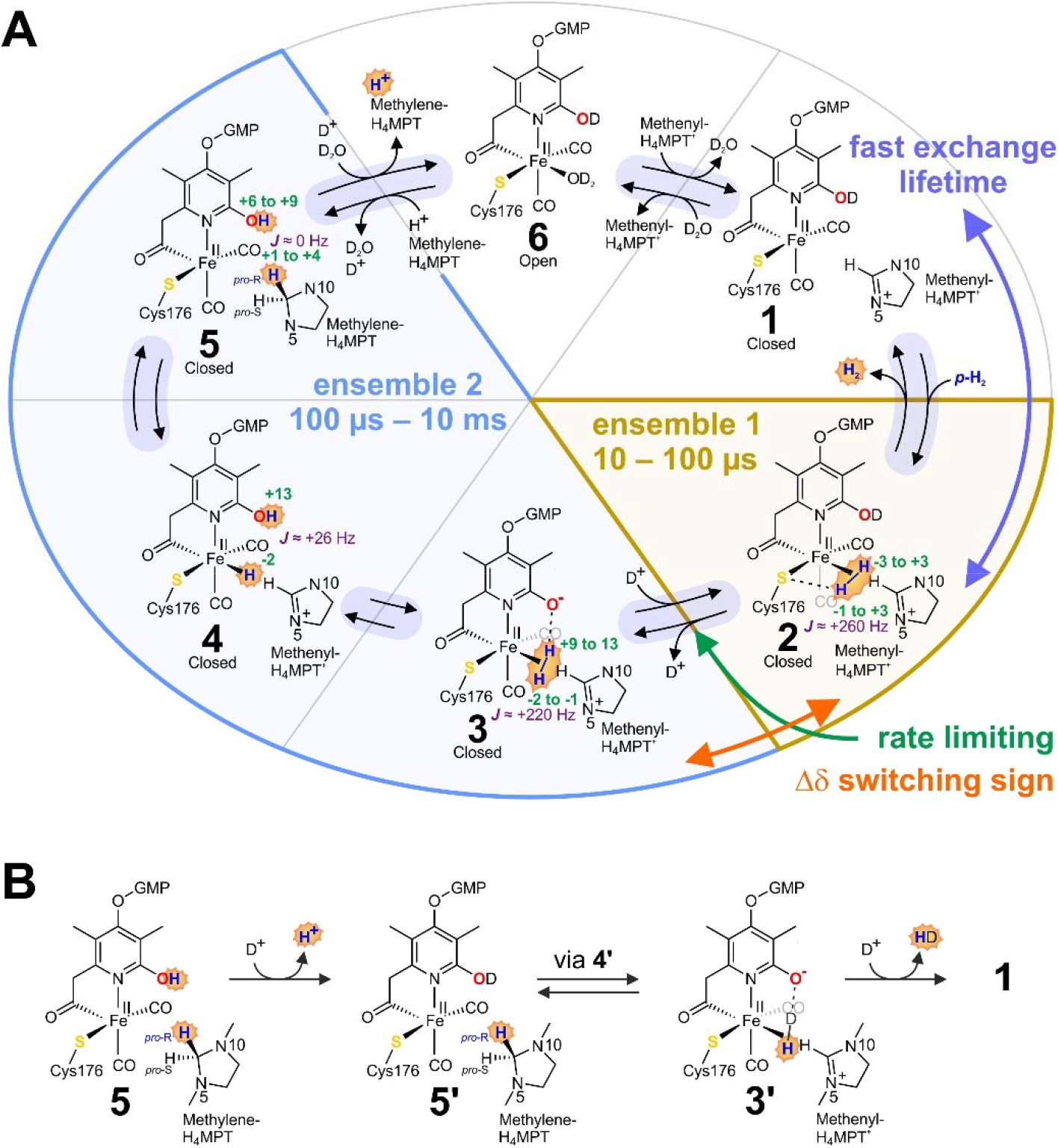
Proposed catalytic mechanism for the creation of PHIP effects by Hmd. (**A**) The suggested catalytic cycle, adjusted for use with *p*-H_2_ in D_2_O buffer. NMR parameters obtained from structural modeling are indicated. Green: Chemical shift ranges from different models [ppm], purple: typical *J*-coupling [Hz]. (**B**) The mechanism proposed for the isotope exchange forming hyperpolarized HD and H^+^. Primes indicate different hydrogen isotope patterns.

In contrast, the HD-PHIP is caused by coherent two-spin evolution under strong coupling in a transiently formed intermediate with a lifetime of *τ*_*HD-PHIP*_ *100 μs - 10 ms* (SI 2.9, Figs. S23 & S24, Table S6), coupled with subsequent hydrogen isotope exchange with the solvent (Fig. 3B). This interpretation is supported by the field dependence of the effect at low magnetic fields (SI 2.4 & 2.5, Fig. S18), which is consistent with a strong coupling-mediated mechanism (SI 2.5).^28,33,34,35^

Comparison between modeled and experimental spectra suggests that multiple intermediate states contribute to the enhancement effects (SI 2.8), which is expected for the facile reversibility of the catalytic reactions and the flat free energy profiles predicted for Hmd.^16,21,36^ The PNL and HD-PHIP effects observed for the same reaction can only be understood when at least two ensembles of bound states are included in the modelling, with one of the ensembles requiring an exchange of protons and deuterium. Whereas PNL and HD-PHIP may be caused by the same or two different *J*-coupled intermediates, placing the H^+^ *⇄* D^+^ exchange in the intermediate causing the PNL is incompatible with the observed isotope exchange kinetics (SI 2.8, Fig. S20).

However, when two bound-state ensembles are included, ^1^H-PHIP spectra (Fig. 2C) and isotope exchange kinetics (SI 2.1-2.3, Fig. S12-S17, Table S4) can be simultaneously reproduced by kinetic modelling. The *J*-coupled intermediate causing the PNL must be placed in the early intermediate, separated from the isotope exchange reaction by the rate-limiting step. Different modeling scenarios for the HD-PHIP are compatible with the data (SI 2.10, Tables S5-S6, Figs. 2D & S26-S28). The scenario requiring only one large, positive *J*-coupling in ensemble 1, and a switch in sign of the chemical shift difference *Δδ* between the two ensembles (Table S6, scenario A) is shown in Fig. 2D.

Only when we request that enhancement effects are predominantly created in an intermediate, where the FeGP cofactor is in the protonated pyridinol form, and when we request that the conformational changes required for pyridinol deprotonation are rate limiting for isotope exchange, a switch in sign of *Δδ* can be explained mechanistically. The new catalytic cycle (Fig. 3A) has to include an interchange of the deprotonation and H_2_-binding steps between **1** and **3**. The change in sign for *Δδ*, derived from the spectra, occurs only if these mechanistic steps are interchanged.^16^ Consequently, intermediate **2** differs from the previously proposed structure (ref 16), which represented the FeGP with vacant binding site in the deprotonated oxypyridine form. In the H_2_-bound state **2** with the FeGP cofactor in the pyridinol-form, the H_2_ orbitals are polarized mainly by interaction with the thiolate ligand (S-Cys176). Deprotonation of the pyridinol switches the sign of *Δδ* because the oxypyridine is the stronger base. This implies that both thiolate and oxypyridine are involved in the catalysis.

Using PNL in chemical exchange saturation transfer (CEST) experiments combined with para-hydrogen is promising with regard to detecting unknown catalytic species.^31^ CEST indirectly detects transiently formed species of low concentration through chemical exchange with an abundant species, i.e. between the enzyme and signal-enhanced hydrogen.^31,37^ Using the combination of PHIP and CEST (PHIP-CEST) for the Hmd system (Figs. S2&S3), we detected CEST effects on signals of the H_2_-PHIP and the HD-PHIP.^31^ The effects manifest in altering signal integrals, measured for H_2_ and HD, as a function of CEST offset (Fig. 4 A&B and Figs. S29-S31); and in changes of signal shape for the H_2_-PHIP-CEST (inserts in Fig. 4B). The CEST effects saturate at spin-lock fields between 0.5 and 1 kHz and thus, provide estimates for the lifetime of the intermediates of *τ*_*PHIP-CEST*_ *1 − 2* ms.

**Fig. 4.**
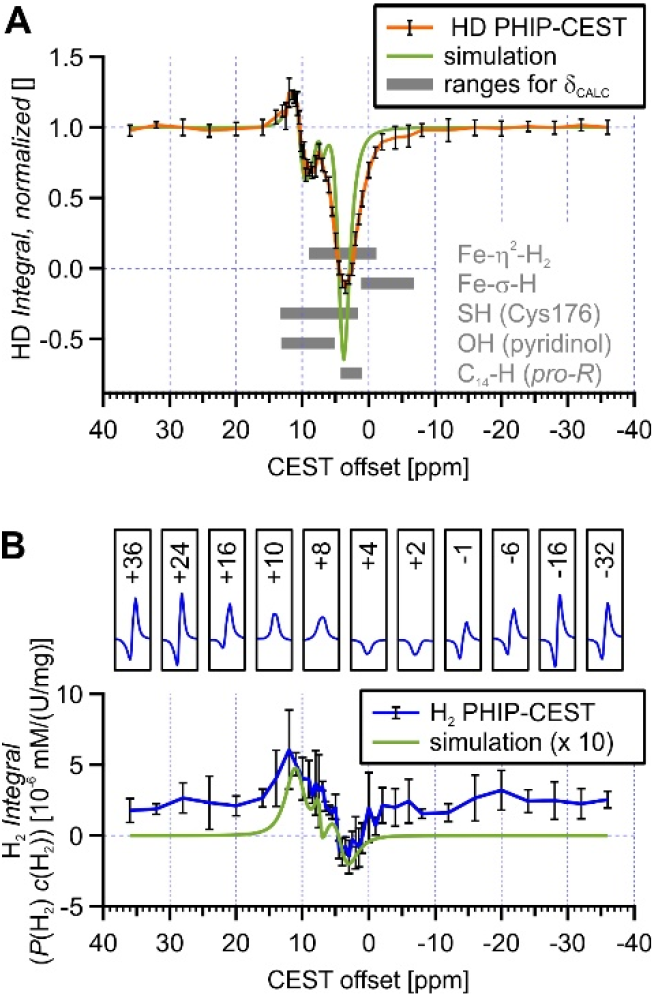
Hyperpolarized CEST experiments for the identification of reaction intermediates. PHIP-CEST experiments performed at 309 K and 14.1 T (600 MHz), with sample conditions as for Fig. 2. (**A**) HD-PHIP-CEST profile obtained from multiple-quantum filtered experiments (Fig. S3), using an 8 s saturation with *γB*_1_ = 666 Hz. The HD signal integral, normalized to the off-resonance integral, measured at -36 ppm, is shown in orange. Data from five samples was averaged, error bars indicate standard deviations. A simulated HD-PHIP-CEST profile (SI 2.11 & Table S7) is overlaid. In grey, the ranges of computed chemical shifts (*δ*_CALC_) obtained from multiple different QM/MM models (S26) are indicated. (**B**) H_2_-PHIP-CEST experiment according to Fig. S2, using a 2 s saturation with *γB*_1_ = 1333 Hz. Top: H_2_ lineshape observed after irradiating at different offsets. Bottom: H_2_-line integral (*polarization?concentration*), normalized by sample activity for hydrogen isotope exchange (activity range: 68 U/mg – 41 U/mg). The average from three samples is shown with standard deviations as error bars. A simulated profile obtained using the same simulation parameters as in panel **A** is shown in green. Simulations underestimate the H_2_-PHIP-CEST effect, and simulated data is up-scaled 10-fold for better visualization of the qualitative agreement obtained.

The PHIP-CEST curves can be reproduced qualitatively in simulations, i.e. assuming chemical shifts of 10.5 ppm (±0.5 ppm) and 4 ppm (±2 ppm) for one of the ensembles, with hydrogen isotope exchange with D_2_O happening at 10.5 ppm (SI 2.11, Table S7). The 10.5 ppm fall into the chemical shift ranges estimated for the pyridinol position of FeGP as well as for previously suggested thiol-ligand intermediates (H-S-Cys176) (SI 2.12-2.14, Tables S9 & S11).^20,21^ However, given the 100-fold reduction in H^+^ *⇄* D^+^ exchange activity in the H14A-jHmd mutant, the pyridinol position appears to be the reasonable choice, (Table S3). For the 4 ppm position, it appears that it is the H_*proR*_ position of CH_2_=H_4_MPT in structure **5** that falls into the expected range, which is in agreement with the model we propose (Fig. 3).

In summary, we demonstrate that sensitivity-enhanced NMR reveals hitherto undetectable bound hydrogen intermediates of hydrogenases. The NMR technique, we first applied here to study a diamagnetic metalloenzyme, is particularly suitable for investigating transient hydrogen intermediates during catalysis, because it specifically and only enhances the signals of bound hydrogen species. We could detect a Fe-*η*^2^-H_2_ species, which is in rapid equilibrium with free H_2_, and we could characterize it in addition to intermediates. Given the possibility to conduct experiments under turnover-conditions at physiological concentrations at unprecedented NMR sensitivity, information that is complementary to other spectroscopic techniques is obtained, i.e. showing catalytic intermediates that can be detected by this approach. Therefore, these techniques can now be used as a general approach for unravelling the catalytic mechanisms of all hydrogenases and their modeled catalysts. In addition, given the high sensitivity of this technology, it holds promise regarding its application to microbial cells, and with regard to exploring new hydrogen metabolisms *in vivo*. Ultimately, all these findings have opened up new insights about the hydrogenases that can now be investigated for its translatability into efficient hydrogenation catalysts that will help to solve our societies’ energy challenges in the future.

## Acknowledgments

The authors thank Prof. Dr. Markus Reiher and Dr. Matthew Woodrich for providing the structural models for Hmd published in ref. 16 and ref. 10, respectively, and for helpful discussions. We thank Prof. Dr. Dorothea Becker for feedback on the manuscript and editing.

## Funding

Max Planck Society (L.K., M.G., G.H., G.L.S., F.N., A.A.A., C.G., S.S. and S.G) German Research Foundation (DFG) grants SPP 1927: SH 87/1-1 and SH 87/1-2 (S.S.) and grants PR 1868/3-1, HO-4602/2-2, HO-4602/3, GRK2154-2019, EXC2167, FOR5042, SFB1479 and TRR287 (A.P. and J.B.H.) German Federal Ministry of Education and Research (BMBF), framework of the e:Med research and funding concept (01ZX1915C) (A.P.) European Regional Development Fund (ERDF) (J.B.H.) Zukunftsprogramm Wirtschaft of Schleswig-Holstein (Project no. 122-09-053) (J.B.H.)

## Author contributions

C.G., A.P. S.G. and S.S. initially conceived the work. M.G. and G.H. performed protein and cofactor preparation. NMR experiments were designed by L.K., A.P., C.G. and S.G.. L.K. performed NMR experiments, data analysis and visualization thereof. A.P., L.K. and M.K. performed kinetics and spin-dynamics modeling. C.R., G.L.S., F.N. and A.A.A. performed structural modeling and I and *J* computations. All authors contributed to data interpretation, L.K. prepared the initial manuscript with contributions from M.G., S.G., M.K. and A.P and all authors contributed to its revision.

## Competing interests

Authors declare that they have no competing interests.

## Notes

### Competing Interest Statement

The authors have declared no competing interest.

