## Supplementary material for "Sensitivity-enhanced magnetic resonance reveals hydrogen intermediates during active [Fe]-hydrogenase catalysis": SI

### 1. Materials and Methods

#### 1.1 Cultivation of *Methanothermobacter marburgensis*

*M. marburgensis* was cultivated anaerobically in a 10-l fermenter under continuous flow of a gas mixture composed of $H_2/CO_2/H_2S$ (80%/20%/0.1%) (*38*).The medium consists of 40 mM $NH_4Cl$, 50 mM $KH_2PO_4$, 24 mM $Na_2CO_3$, 0.5 mM nitrilotriacetic acid (NTA), 0.2 mM $MgCl_2·6\,H_2O$, 1 µM $CoCl_2·6\,H_2O$, 1 µM $Na_2MoO_4·2\,H_2O$, 50 µM $FeCl_2$, 5 µM $NiCl_2$ and 20 µM resazurin (final concentrations) (*38*). To isolate $H_4MPT$ from the cells, *M. marburgensis* was cultivated in the full medium. In the case of Hmd purification, *M. marburgensis* was cultivated under nickel-limiting conditions, where $NiCl_2$ was omitted from the medium. In the nickel-limiting culture, a trace amount of nickel is supplied by erosion from the metal parts of the fermenter. For the preparation of $^{57}$Fe-enriched FeGP cofactor, 50 µM [$^{57}$Fe]-$FeCl_2$ were added to the medium instead of non-enriched $FeCl_2$. [$^{57}$Fe]-$FeCl_2$ was prepared by the treatment of $^{57}$Fe-enriched metal (96% enrichment) in 38% HCl solution. In the nickel-sufficient culture, when the culture reached an optical density (OD) of ~6–7 at the late exponential growth phase, the cells were harvested. In the case of nickel-limiting condition, the culture growth, which started from 5% inoculation of pre-culture, became slower after overnight culture at OD ~4. The slow growth of the culture with doubling time of ~11 h under the nickel-limiting conditions continued until OD ~5–6. The culture in the fermenter was cooled down by circulating ice water and then anaerobically harvested via continuous-flow centrifugation and the cells were stored at −75 °C.

#### 1.2 Purification of Hmd from *M. marburgensis*

Purification of Hmd from *M. marburgensis* (mHmd) was performed under strictly anaerobic conditions in an anaerobic glove box (Coy Laboratories, Grass Lake, MI). Centrifugation was performed using a plastic tube with a screw cap and a rubber O-ring. Around 100 g of *M. marburgensis* cells were suspended in 200 ml of 50 mM potassium phosphate buffer (KPP) pH 7.0 and sonicated (80% power of 100 W with 6 times of 8 min on/7 min off cycles) with SONOPLUS HD 200 from BANDELIN (Berlin) using a VS 70 T tip. The crude extract was centrifuged for 30 min at 140 000 × g and 4 °C. Ammonium sulfate powder was added to the supernatant until 60 % saturation. After 20 min incubation on ice, the supernatant was centrifuged for 20 min at 13 000 × g and 4 °C, and then ammonium sulfate powder was added to the supernatant until 90 % saturation. After another 20 min incubation on ice, the suspension was centrifuged for 20 min at 13 000 × g and 4 °C. Afterwards, the pellet was suspended in 15 ml of 50 mM 3-(*N*-morpholino)propanesulfonic acid (MOPS)/KOH pH 7.0. The suspension was dialyzed at 4 °C overnight against 50 mM citrate/NaOH pH 5.0. The dialyzed solution was centrifuged for 20 min at 17 000 × g and 4 °C, and the supernatant was applied to a Source 30Q column (300 ml column volume) equilibrated with 50 mM citrate/NaOH pH 5.0. The column was washed with 50 mM citrate/NaOH pH 5.0 containing 200 mM NaCl. Elution took place with a linear gradient from 200 mM to 500 mM NaCl in 500 ml. Fractions of 10 ml were immediately neutralized with 1.0 ml of 1 M MOPS/KOH pH 7.0 and 0.06 ml 1 M NaOH. Fractions containing mHmd were pooled and concentrated via an Amicon ultrafilter (30 kDa cut off). Afterwards, the concentrated solution was applied to a HiPrep 26/10 desalting column equilibrated with $H_2O$. The elution was carried out with $H_2O$ and the fractions containing mHmd were pooled and concentrated with an Amicon ultrafilter (30 kDa cut off). Finally, purified mHmd was flash frozen in liquid nitrogen and stored at –75 °C.

1.3 Extraction of the FeGP cofactor

The FeGP cofactor was extracted from 100 mg of mHmd by incubation in 60% MeOH, 1 mM 2-mercaptoethanol and 1% $NH_3$ in a final volume of 12 ml for 15 min at 40 °C. Subsequently, the FeGP cofactor was separated from the denatured protein trough filtration with an Amicon filter (10 kDa cut off). The filtrate was collected and evaporated at 4 °C . The concentrated FeGP cofactor solution (~50 µl) was diluted to 1 ml with 10 mM ammonium carbonate pH 9 containing 1 mM 2-mercaptoethanol. The five aliquots of 200 µl were stored in liquid nitrogen.

1.4 Production of jHmd apoenzyme

The apoenzymes of Hmd from *Methanocaldococcus jannaschii* (jHmd); the wild type and the H14A mutant were produced in *E. coli* BL21(DE3) (*19*). For a pre-culture, 100 ml Luria-Bertani (LB) medium with 30 µg/ml kanamycin were inoculated with a frozen glycerol stock of the *E. coli* cells harboring the corresponding plasmid. The pre-culture was shaken at 37 °C overnight and used to inoculate 2 l of tryptone-phosphate (TP) medium (*39*), which contained 30 µg/ml kanamycin. When the culture reached an $OD_{600}$ of 1, the expression of *jHmd* was induced with a final concentration of 1 mM isopropyl β-D-1-thiogalactopyranoside (IPTG). After 3 h of expression, the cells were harvested by centrifugation for 20 min at 13 000 × g and 4 °C. The cells were suspended in 50 mM MOPS/KOH pH 7.0 containing 1 mM dithiothreitol (DTT) and disrupted by sonication (80% power of 100 W with 5 times of 4-min on/4-min off cycles) using a MS 72 tip. The debris was removed via ultracentrifugation for 40 min at 130 000 × g and 4 °C, and the supernatant was heated for 15 min at 70 °C to denature the *E. coli* proteins. By centrifugation for 20 min at 13 000 × g and 4 °C, the denatured proteins were removed. Afterwards, ammonium sulfate powder was slowly added to a final concentration of 2 M. Then the precipitated proteins were removed again via centrifugation for 20 min at 13 000 × g and 4 °C. The supernatant was applied to a Phenyl-Sepharose column (50 ml column volume) equilibrated with 50 mM MOPS/KOH pH7 containing 1 mM DTT and 2 M ammonium sulfate. The proteins were eluted with a 200 ml linear gradient from 2 M to 0 M ammonium sulfate. Each 10 ml was fractionated and analyzed by sodium dodecyl sulfate–polyacrylamide gel electrophoresis (SDS-PAGE). The fractions containing jHmd were pooled and concentrated to 10 ml with an Amicon ultrafilter (30 kDa cut off). Then the solution was desalted by a HiPrep 26/10 desalting column equilibrated with 50 mM MOPS/KOH pH 7.0 containing 1 mM DTT. The apoenzyme was flash frozen and then stored at –75 °C.

1.5 Reconstitution of the jHmd holoenzyme

To reconstitute the jHmd holoenzyme, 125 µM of the apoenzyme (either wild type or H14A mutant) were mixed with 175 µM of the purified FeGP cofactor. To get rid of the non-incorporated FeGP cofactor, the solution was applied to a HiPrep 26/10 desalting column equilibrated with 10 mM MOPS/KOH pH 7.0. All procedures were performed under yellow light in an anaerobic tent.

1.6 Purification of $H_4MPT$ and conversion to $CH_2$=$H_4MPT$, $^{13}CH_2$=$H_4MPT$, $CD_2$=$H_4MPT$

For the purification of $H_4MPT$, 130 g of *M. marburgensis* cells (nickel-sufficient growth condition) were suspended in 130 ml of 50 mM MOPS/NaOH pH 6.8 and heated to 60°C in a water bath. *N*,*N*,*N*-trimethylhexadecan-1-aminium bromide (CTAB) was added to a final concentration of 1% and the suspension was incubated for 6 min at 60 °C. Afterwards, the suspension was cooled down in an ice bath for 30 min. In an anaerobic chamber, the pH was adjusted to 3 using 100% formic acid and the suspension was centrifuged for 60 min at 6800 × g and 4 °C. The supernatant was separated from the pellet and put on a Serdolit PAD II column (SERVA, Heidelberg, Germany) equilibrated with XAD buffer ($H_2O$-formic acid buffer (69:1), pH 3 adjusted by NaOH). The column was washed with XAD buffer and eluted with 15% methanol in XAD buffer. The $H_4MPT$-containing fractions were pooled and evaporated. Subsequently, the lyophilized preparation was solubilized in 50 ml of $H_2O$ and pH was adjusted with 100% formic acid to 3. The solution was loaded on a Serdolit PAD I column (SERVA, Heidelberg, Germany) equilibrated with XAD buffer. The column was washed with 0.1% formic acid in $H_2O$ and eluted with 30% methanol containing 0.1% formic acid in $H_2O$. The $H_4MPT$-containing fractions were pooled, lyophilized and the concentration was adjusted by adding $H_2O$. For the conversion of $H_4MPT$ to $CH_2$=$H_4MPT$, $^{13}CH_2$=$H_4MPT$ and $CD_2$=$H_4MPT$, 500 µl of 2 mM $H_4MPT$ were mixed with 15 µl 200 mM HCHO, [$^{13}C$]-HCHO or [$^{2}H_2$]-HCHO. The conversion to $CD_2$=$H_4MPT$ took place in $D_2O$. The converted solutions were evaporated and the concentrations were adjusted with $H_2O$ or $D_2O$. [$^{13}C$]-HCHO or [$^{2}H_2$]-HCHO and $D_2O$ were purchased from Cambridge Isotope Laboratories (Tewksbury, MA).

1.7 Activity Assay for methylene-$H_4MPT$ to methenyl-$H_4MPT^+$ conversion using UV/Vis spectroscopy

A 1 ml quartz cuvette containing 680 µl of anoxic 120 mM potassium phosphate buffer pH 6 with 1 mM EDTA under 100% $N_2$ was shielded with a rubber stopper and pre-incubated at 40 °C for 5 min. Seven µl of 2 mM methylene-$H_4MPT$ was added and then the assay was started by the addition of 10 µl of the enzyme solution. The increase of the absorbance of methenyl-$H_4MPT^+$ at 336 nm ($\varepsilon_{336}$ = 21.6 $mM^{-1}\cdot cm^{-1}$) at 40 °C was measured to calculate the enzyme activity. One unit (U) of the enzyme activity is the amount catalyzing the formation of 1 $\mu mol\cdot min^{-1}$ methenyl-$H_4MPT^+$ (*40*). The specific activity ($\mu mol\cdot min^{-1}\cdot mg^{-1}$) was calculated using the enzyme concentration in the assay cuvette. The enzyme concentration was determined by the Bradford method using bovine serum albumin as standard (*41*). The measured activity under these conditions was 400 $U\cdot mg^{-1}$ for the reconstituted wild-type jHmd.

1.8 Activity Assay for hydrogen isotope exchange activity using NMR

To measure the sample activity *in-situ* in the NMR, the sample activity for hydrogen isotope exchange ($H_2 \rightleftarrows HD \rightleftarrows D_2$) was determined. Samples were bubbled with 7 bars (gauge pressure) of *n*-$H_2$ and the concentrations of $H_2$ and HD in solution were measured by series of small flip-angle $^{1}H$ experiments, according toFig. S1. The disappearance of the $H_2$ signal, as well as the appearance and fading of the HD signal were observed in series of small flip-angle experiments. For details of sample preparation and measurement, see sections S1.9 - S1.11.

Spectra were processed, as described in section S1.11, and the $H_2$ integrals were extracted using a localized 3rd order polynomial baseline correction close to the $H_2$ line (to reduce bias from the nearby water). The decay of the $H_2$-concentration was fitted to

$$[H_2](t) = [H_2]_0 \exp(-v_0 t), \quad (1)$$

keeping the hydrogen concentration measured in the first spectrum after bubbling $[H_2]_0 = [H_2](t = 0)$ as a free fit parameter. From the fitted parameters, the activities stated were computed as

$$v_0(H_2\ consumption) = \frac{[H_2]_0 v_0}{[Hmd]_0 M(Hmd)}. \quad (2)$$

$[Hmd]_0$ was computed from the enzyme concentration determination via the Bradford method (see section 1.7), previously performed for the same stocks. For these activity tests, one unit (U) of the enzyme activity thus is the amount catalyzing the consumption of 1 $\mu mol \cdot min^{-1}$ $H_2$.

All values stated refer to measurements at 309 K, for samples prepared from 1 µM *j*Hmd and 3 µM $^{13}CH_2$=$H_4$MPT in 600µL $D_2O$-buffer, pD 6.0, 1 mM EDTA, 120 mM potassium phosphate. The measured specific activity for fresh stocks of wild-type jHmd was 80 U $mg^{-1}$.

An exception to the stated condition applies to the values stated in Table S3, which were measured at pD 7.0.

### 1.9 NMR Sample preparation

NMR samples were prepared by mixing stocks of the corresponding forms of Hmd and metheylene-$H_4$MPT with $D_2O$-buffer inside 5 mm NMR tubes under a stream of $N_2$ or Argon. All stocks were handled in brown-glass vials under inert atmosphere, using microliter syringes. All sample handling was performed in the dark. Used sample concentrations are stated in the Figure and Table captions. After mounting the samples to the systems for bubbling NMR samples inside the NMR spectrometers (described in section 1.10), all samples were bubbled with $N_2$ for at least 1 minute, to convert the metheylene-$H_4$MPT used for sample preparation to methenyl-$H_4$MPT *in situ*.

Deuterated 120 mM potassium phosphate buffers containing 1 mM EDTA were prepared at varying *p*D (see figure captions) and degassed before use by bubbling with $N_2$. Since EDTA was used as internal concentration reference for all experiments, the EDTA concentration was checked for all buffer stocks after preparation, against an internal standard (maleic acid). During experiments with Hmd, no maleic acid was present.

The chemical shift referencing to DSS (3-(Trimethylsilyl)-1-propansulfonic acid) was performed by adding 1 mM DSS to the buffers prepared, and tabulating the HDO chemical shift for all buffer *p*D values and temperatures used. For measurements with Hmd, the HDO signal was used as chemical shift reference, using the tabulated values.

$D_2O$ (99.9%), $K_3PO_4$, DSS, and EDTA-dianhydride were purchased from Sigma Aldrich. $KD_2PO_4$ was either also purchased from Sigma Aldrich, or prepared from $KH_2PO_4$ (Carl Roth), using isotope exchange with $D_2O$.

1.10 NMR instrumentation

NMR experiments were performed on four different spectrometers equipped with home-built bubbling setups, which provide the ability to handle the samples under inert atmosphere, and saturate the solutions with different gases within a few seconds (5 – 15 s) by gas bubbling, while the sample resides inside the spectrometer. The bubbling setups were equipped with three gas channels ($N_2$, *p*-$H_2$ and the third channel used for *n*-$H_2$ or mixtures of $N_2$ and *n*-$H_2$) with flow control by needle valves, and magnet valves switched during the NMR pulse sequence, controlled by the TTL output signals of the spectrometer. Gas pressure inside the sample volume was cycled between room pressure and 7 bar (gauge pressure), using pressure regulators at the inlets of the three gas channels and a backpressure regulator at the gas outlet. For experiments where mixtures of $N_2$ and *n*-$H_2$ were used, the two gases were pre-mixed at defined ratios in a storage container. *p*-$H_2$ was produced from *n*-$H_2$ by a Bruker parahydrogen generator (BPHG 90) or (for Fig. S9 and Table S3 only) by a parahydrogen generator from ColdEdge International Inc., operating at 20 K for 99% *p*-$H_2$ enrichment. *P*-$H_2$ was either directly supplied to the bubbling setup, or stored in 1 L aluminum bottles for transport, and supplied to the bubbling setups from the aluminum vessels. For experiments using the transport bottles, the *p*-$H_2$ content was measured before and after every experimental series by saturating samples containing the neat buffer either with *n*-$H_2$ or with *p*-$H_2$ at 7 bars and comparing the thermally polarized $^1$H signals for $H_2$ in both cases. For the *p*-$H_2$ produced by the Bruker parahydrogen generator (BPHG 90) the enrichment was measured in regular intervals, and was found to be (87 ± 2)%. Measured *p*-$H_2$ enrichments are given with all figures.

Experiments at 14.10 T ($^1$H: 599.9 MHz) were performed on an Avance III HD narrow bore system (Bruker Biospin GmbH, Karlsruhe), equipped with a 5 mm inverse quadruple resonance cryoprobe (QCI, $^1$H/$^2$H/$^{13}$C/$^{15}$N/$^{31}$P) with z-Gradient. 90° pulses on this system were 11.25 µs for $^1$H and 120 µs for $^2$H. A BCU-II chiller (Bruker Biospin, Karlsruhe) was used for temperature regulation. TopSpin 3.6 patchlevel 2 was used for acquisition.
Experiments at 21.15 T ($^1$H: 900.1 MHz) were performed on an Avance III HD narrow bore system (Bruker Biospin GmbH, Karlsruhe), equipped with a 5 mm inverse triple resonance cryoprobe (TCI, $^1$H/$^2$H/$^{13}$C/$^{15}$N) with z-Gradient. The 90° pulse for $^1$H on this system was 12.5 µs. A BCU-II chiller (Bruker Biospin, Karlsruhe) was used for temperature regulation. TopSpin 3.5 patchlevel 7 was used for acquisition.
Experiments at 7.05 T were performed at two different instruments, both equipped with 5 mm direct double resonance room-temperature probes (BBFO, BB/$^1$H/$^2$H). The first system was an Avance III HD narrow bore system (Bruker Biospin GmbH, Karlsruhe) operated at 300.29 MHz $^1$H frequency, equipped with a BCU-I chiller (Bruker Biospin GmbH, Karlsruhe). The 90° pulse for $^1$H on this system was 14.5 µs and TopSpin 3.5 patchlevel 7 was used for acquisition. The second system was an Avance IV (NEO) wide bore system (Bruker Biospin GmbH, Karlsruhe) operated at 300.13 MHz $^1$H frequency, equipped with a narrow bore shim insert (BOSS/3). This system was not equipped with a chiller. The 90° pulses for on this system were 10.25 µs for $^1$H and 230 µs for $^2$H. TopSpin 4.0 patchlevel 7 was used for acquisition on this system.

For T < 330K, sample temperatures were calibrated at each instrument using an externally calibrated Pt100 resistance thermometer, and were cross-checked with a methanol-$d_4$ sample (99.8%) (*42*). Agreement between both methods of temperature calibration was better than

± 0.2 K. Fig. S6 & Fig. S7 and Table S1 were acquired prior to subjecting all instruments to the same temperature calibration, and the sample temperatures stated were back-calculated, using the above-mentioned calibration, from the difference of HDO and HD chemical shifts. For T ≥ 330K, 80% 1,2-ethanediol in DMSO-$d_6$ was used for cross-checking, which yielded ± 0.5 K offset from the Pt100 results.

1.11 NMR experiments

NMR experiments with sample bubbling were performed according to the experiment schemes in Figs. S1 – S4. Towards the start of all experiments, the samples were bubbled with $N_2$ for 30 s and subsequently with $p$-$H_2$, $n$-$H_2$, or mixtures of $n$-$H_2$ and $N_2$ for bubbling periods of $\tau_{\text{bubbling}}$ of 8 s (7 T instruments) or 15 s (14 T & 21 T instruments), with intermittent pressure release to atmospheric pressure. $p$-$H_2$ was used for PHIP experiments, $n$-$H_2$ was used for measurement of sample activity and reference experiments, and mixtures of $n$-$H_2$ and $N_2$ were used for the enzyme kinetics analysis shown in chapter SI S2.2, Table S4 and Fig. S14. A 1 s delay (2 s for experiments with manual field-cycling) was introduced between the endpoint of $H_2$-bubbling and the beginning of pulsing and acquisition to wait for bubbles to leave the detection volume. Sample pressure was maintained constant at 7.0 bars during acquisition.

All $^1$H-PHIP spectra shown were acquired with a single-scan using 16° - 45° hard pulses, on-resonance with the water signal. All $^1$H-PHIP spectra shown are the first spectrum after bubbling $p$-$H_2$, acquired in a series of single scan spectra ($n$ = 8 - 128), according to Fig. S1. Isotope exchange kinetics were monitored with series of pulse-acquisition-gradient experiments after bubbling $n$-$H_2$ or mixtures of $n$-$H_2$ and $N_2$ according to the same experimental scheme. For sample activity measurements at 309 K, isotope exchange was followed for 256 s at 7.05 T or for 512 s at 14.10 T and 21.15 T with a 4 s time increment, one scan per time-point and a single time of hydrogen bubbling for the activity measurement.

FIDs were acquired for 3.28 s (exception: Fig. S6 & Fig. S7 with 0.82 s) length were acquired, collecting 16384 complex data-points with a dwell-time of 200 µs at 7.05 T or 32768 complex data-points with a dwell-time of 100 µs at 14.10 T and 21.15 T (exception: Fig. S6 & Fig. S7 with 16384 complex data-points at 7.05 T and 8192 complex data-points at 21.15 T). Data was apodized with a monoexponentially decaying apodization function with 0.3 Hz linebroadening (1.0 Hz for Fig. S6 & Fig. S7), zero-filled to double the amount of time-points and Fourier-transformed. Prior to the integration of the signals close to the HDO residual peak, localized base-line corrections with a 3rd order polynomial were performed, to remove bias from the foot of the water signal.

PHIP-CEST data was acquired using the pulse sequence schemes shown in Fig. S2 (for $H_2$-PHIP-CEST) and Fig. S3 (for HD-PHIP-CEST). All PHIP-CEST spectra were acquired as single scan spectra without phase cycling and with all pulse phases at zero. CEST saturation was applied after stopping the bubbling and after a 1 s delay for the sample to settle, for either 2 s ($H_2$-PHIP-CEST) or 8 s (HD-PHIP-CEST). Nominal spin-lock-field amplitudes ($\gamma_H B_1$) were varied between 5 Hz and 3 kHz, as indicated in the figure legends. Spin-lock-field offsets were varied according to randomly shuffled lists of 20 - 51 offset points dispersed over the range of -36 ppm to +36 ppm relative to the DSS resonance.

For HD-PHIP-CEST experiments, profiles were obtained by direct integration of the two low-field peaks of the HD triplet. The high-field peak is overlapped with artifacts from non-ideal water suppression. For $H_2$-PHIP-CEST experiments, an automated $0^{th}$ order phase correction was applied to all spectra and a localized automated $3^{rd}$ order baseline correction was applied to a 0.15 ppm region was applied around the $H_2$ signal region prior to integration of the $H_2$ peak, to reduce the impact of the HDO signal base on the measured integral of $H_2$. In all cases, integrals over the real part of the spectrum are reported.

The PHIP-CEST curves shown are averages of measurements from one to five samples measured, as stated in the figure legends. Appropriate correction for differences in sample activity were crucial, thus the following, detailed description: Before and after each set of PHIP-CEST experiments, the sample activity for hydrogen isotope exchange (see section S2.3) was monitored using the experimental scheme shown in Fig. S1. Usually sample activity decreased by about ~20% during the collection of the 20-64 different PHIP experiments. Sample activities ranged from 82 µmol $H_2$ $min^{-1}$ $mg^{-1}$ to 19 µmol $H_2$ $min^{-1}$ $mg^{-1}$, depending on the age and the storage conditions of the stocks. Integrals measured PHIP-CEST experiments were scaled by the measured activity, assuming a time-proportional decrease of the sample activity during measurement, between the activity measurement before and after the PHIP-CEST experiments. An exemption are three out of the five experiments used for Fig. 4A of the main text, and the experiments shown in Fig. S31. For these experiments, the list of sampled CEST offset-points was interleaved with regular measurements of spectra with irradiation at -36 ppm. These measurements were used to fit a monoexponential decay with offset to the measured signal integrals (reflecting sample activity), to account for a rapid decrease of the sample activity typically observed during the first experiments, before sample deactivation slowed down.

Field-cycling data was collected using manual field-cycling, according to the scheme shown in Fig. S4. Manual field-cycling was performed between the end of the $p$-$H_2$-bubbling and prior to the $^1H$-pulse and took less than 2 s. Field-cycling was performed by lifting the sample out of the magnet center on a stick to marked positions, which indicated previously deteremined low-field positions between $B_{bubbling} = 1\ mT - 6\ T$. For measurements at $B_{bubbling} = 7\ T$, the sample was not shuttled, but resided in the probe. A single scan was acquired for each spectrum. To determine the locations of the low field positions, the field profile inside of the magnet bore was mapped in the $1\ mT - 3.5\ T$ range using a Gaussmeter, and the locations of the field positions between 3.5 $T$ and 7 $T$ were interpolated, fitting a sigmoidal function to the measured field profile.

To avoid possible temperature gradients along the field-cycling paths, the temperature control of the spectrometer was shut off prior to the experiment overnight, and the field cycling data was acquired at room-temperature (292.1 K, measured by the thermocouple of the probe, 292.5 K measured at 1 mT position, which is still inside the bore). Hydrogen isotope exchange kinetics were characterized at 292.2 K after the measurement, using the pulse sequence of Fig. S1, collecting single-can spectra in intervals of 4 s over 128 s after bubbling $n$-$H_2$.

### 1.12 NMR data representation: Field independent scales for intensities and integrals

NMR data in this paper is represented on field-independent absolute scales for signal intensity or integral, rather than the usual representation in arbitrary units. These scales were introduced, to facilitate the comparison of signal intensities and integrals measured at different fields.

For idealized single-pulse experiments, the NMR signal integral $I_k$ for a group of spin-1/2 nuclei $k$ is proportional to the product of their concentration $c_k$ and their polarization $P_k$ prior to the pulse.

$$I_k \propto c_k P_k \tag{3}$$

For thermally polarized samples, $P_k$ equals the thermal equilibrium polarization $P_0$.

$$P_0 \approx \frac{\gamma_k \hbar B_0}{2 k_B T} \tag{4}$$

Using the 1 mM EDTA present in all buffers as an internal concentration reference, a field independent representation of the signal integrals is possible, according to

$$c_k P_k = c_{ref} P_0 \frac{\tilde{I}_k}{\tilde{I}_{ref}}. \tag{5}$$

This assumes, that the nuclear spins of the concentration reference were at thermal equilibrium prior to the experiment, and that a simple pulse-acquisition experiment was performed.

For convenience, we introduce the proportionality constant $u$ that relates the signal integrals $\tilde{I}$ given in arbitrary units with the product $c_k P_k$.

$$u = \frac{c_k P_k}{\tilde{I}_k} = \frac{c_{ref} P_0}{\tilde{I}_{ref}} \tag{6}$$

The intensity axis scaling used e.g. in Fig. 2 of the main article can be obtained with knowledge of $u$ from

$$\frac{c_k P_k}{\nu} = u * \frac{SI}{SW} * Intensity\ [a.u.], \tag{7}$$

where $Intensity\ [a.u.]$ is the signal intensity provided by the instrument in arbitrary units, $SW$ is the spectral width of the spectrum and $SI$ is the number of complex points used for discrete Fourier transformation.

### 1.13 Simulation of NMR spectra and CEST curves

Nuclear spin dynamics calculations were performed in MATLAB® R2020b, using the MOIN spin simulation library (*43*). The details of the simulations (equations and explanatory illustrations) are given in the chapters 2.5 – 2.10 of the supplementary text.

1.14 Structural Modeling & Chemical Shift Computation

Chemical shifts and *J*-couplings were computed for a series of QM/MM models derived from two different sources: Firstly, QM/MM models were constructed based on the high-resolution closed-conformation crystal structure (PDB: 6hav) published in ref. (*10*). Secondly, the QM/MM models published in figures 9 & 11 of ref. (*16*), which are derived from MD-simulation snapshots, were used, which were kindly supplied by the authors. The QM/MM models from both sources were (re)optimized and used to construct a series of possible hydrogen-bound intermediates of the Hmd active site, summarized in Tables S8 - S11.

The QM region used in this work is sketched in Fig. S32.Fig. S32. It includes the FeGP cofactor up to the phosphate linker, the side chain of Cys176 coordinating to Fe of FeGP, the pterine, imidazoline and phenyl part of the methenyl-/methylene-$H_4MPT$, as well as the hydrogens originating from $H_2$, which are modeled into the active site. This equals the QM region previously used in ref. (*16*). For the models based on the crystal structure (models E and G), the His14 residue was included into the QM region, due to the close proximity of the $N_\varepsilon$ of this residue to the oxygen at the 2-oxypyridine position of FeGP (3.3 Å).

The active region for geometry optimization was built around the Fe center. It includes all atoms that have a distance of less than 5 Å to the iron center of FeGP, plus the backbone or sidechain (for proteic residues) or full molecules (for non-proteic groups) that these atoms belong to. For the crystal structure derived models E and G, this includes full molecules of FeGP, methenyl-/methylene-$H_4MPT$ and waters Wat598 and Wat731, the full residues of Cys176, Pro202, Val205, and Pro206, and the side chain atoms of His14, Trp148 and His201. For the MD-derived models A and C, this includes the full molecules of FeGP, methenyl-/methylene-$H_4MPT$ and waters Wat1344 and Wat1070, the full residues of Pro202 and Val205, and the side chain atoms of Trp148, Cys176 and His201.

QM/MM calculations were carried out using the ORCA software (*44*). ORCA's default QM/MM settings were used: Additive QM/MM with electrostatic embedding (*45*), link atom approach and using the charge shifting scheme (*46*) to avoid overpolarization of the electron density at the QM-MM boundary. For the MM part the AMBER topology published in ref. (*16*) was used for models A & C, and was prepared using the open forcefield toolkit for models D to G (after conversion to the prms format as required by the ORCA software using the orca_mm module).

During geometry optimization only the atomic positions of the active region were optimized, while the positions of all other atoms were kept frozen. The TPSS density functional (*47*) together with Grimme's D3BJ dispersion correction (*48, 49*) was used in conjunction with the def2-TZVP basis set (*50*) and the def2/J auxiliary basis (*51*).

NMR shielding calculations were performed at the DFT level (*52*), using the TPSS functional (*47*), the pcSseg-2 basis set (*53*) (abbreviated to "pS2" below), and def2/JK auxiliary basis (*51*). Only atoms in the QM region were treated at this level, while the MM region was included as point charges. $^1H$ chemical shifts were calculated with respect to tetramethylsilane (TMS), whose geometry was optimized at the TPSS-D3BJ/def2-TZVP/CPCM(water) level and NMR shieldings calculated at the TPSS/pS2/CPCM(water) level. Gauge-including atomic orbitals (GIAOs) were employed in all shielding calculations and the ad-hoc gauge-invariant treatment of the kinetic energy density τ was used (ORCA keyword "TAU=GI") (*54*).

Indirect nuclear spin-spin coupling constants were calculated using the PBE0 hybrid functional (*55*) and the pcJ-2 basis set (*56*) (abbreviated to “pJ2” below), together with the def2-TZVPP basis set (*50*) for Fe. The isotropic parts of the full coupling tensors are reported as scalar *J*-couplings. Once again, electrostatic QM/MM embedding was applied. All contributions to the spin-spin coupling (Fermi contact, spin-dipole, diamagnetic and paramagnetic spin-orbit) were included in the calculations.

In order to gauge the uncertainty of the calculated NMR properties, several calculations using different density functionals, basis sets, and treatments of the environment on a few arbitrarily chosen models were performed, as described in section S2.12 (Tables S12 – S15).

### 2. Supplementary Text

#### 2.1 Properties of the PHIP effects

No PHIP effects were observed in the absence of *j*Hmd and methenyl-$H_4MPT^+$ (see Fig. S5), or with the H14A-*j*Hmd mutant (see Fig. S9Fig. S9. and Table S3), which is about 100 fold less active than the wild type (*11*), in the presence of methenyl-$H_4MPT^+$. The generation of PHIP effects thus requires the active enzyme substrate complex.

The PHIP experiments can be repeated for a single sample (within the limits of sample stability), simply by bubbling *p*-$H_2$ through the sample again. Fig. S6Fig. S6. shows the overlay of six experiments performed with interleafed bubbling of *p*-$H_2$ and *n*-$H_2$. The HD-PHIP signal intensity is largest close to the conditions of maximum enzyme activity, which are around 330 K (see Fig. S11Fig. S11.), whereas the maximum out-of-phase PHIP signal for for $H_2$ is found at around 320 K. Both PHIP intensities peak around pD 6 (see Fig. S10Fig. S10.), which is also around the pD of maximum sample activity.

While higher *j*Hmd concentrations and higher temperatures produce bigger PHIP signals for HD (compare Fig. S7Fig. S7. and Fig. S8), the enzyme kinetics are quickly becoming too fast for convenient monitoring via NMR. In this limit, fast isotope exchange according to the reaction shown in Fig. 1C of the main article also significantly reduces the PHIP signal intensities observed, since the hyperpolarized species $H_2$ and HD are quickly getting converted to $D_2$.

To narrow down which mechanisms could be causing the PHIP effects, we performed isotope labeling experiments (see Fig. S9 and Table S3) and field dependent PHIP experiments (Fig. S7Fig. S7. & Fig. S8 and Tables S1 & S2). Isotope labeling excludes possible three spin interactions between *p*-$H_2$ and the methenyl-group of methenyl-$H_4MPT^+$ or between *p*-$H_2$ and $^{57}Fe$ as the source of the PHIP effects (see Fig. S9 and Table S3).

Field dependent PHIP experiments show a monotonic increase of the $H_2$-PHIP effect in the range of 7.0 – 21.1 T. This is in line with the behavior expected for PNL lines in the limit of fast dissociation, as discussed in section S2.6. The observed lineshape for the PNL fits well the expectations for the PNL-effect, which provides strong support that this is the dominant PHIP mechanism for $H_2$. For the HD-PHIP, field cycling experiments probing the field dependence of the HD-PHIP effect in the 1 mT – 7.0 T range gave strong support for a strong coupling driven mechanism in this field range, as outlined in (see sections S2.4 & S2.5). The observed slight increase of the HD-PHIP from 14.1 T to 21.1 T may hint at mixed contributions from different PHIP mechanisms at high fields, yet, given the estimated error margins, this is speculative.

2.2 Fitting of Enzyme kinetics

The kinetics of the hydrogen isotope exchange reactions (Fig. 1C of the main article) catalyzed by Hmd can be monitored conveniently using the bubbling-acquisition experiment shown in Fig. S1.Fig. S1. For monitoring the isotope exchange kinetics, samples prepared in $D_2O$ buffer were bubbled with $n$-$H_2$ or mixtures of $n$-$H_2$ and $N_2$, and the signal integrals of $H_2$ and HD were measured in series of small flip-angle $^1$H-experiments after stopping the bubbling, as detailed in section 1.11. $H_2$ and HD concentrations were computed from the measured integrals, compared to the integrals of the internal standard (1 mM EDTA (see section 1.12)), after correction for signal attenuation by fast pulsing.

For fitting the observed kinetics, we chose the kinetic model shown in Fig. S12Fig. S12. The model is a variation of the model recently discussed by Leroux *et al.* (*57*), which explicitly incorportates preferential isotope exchange at one of the two binding sites and site exchange via $k_{ex}$ between these two binding sites. In the limit $k_{ex} \rightarrow \infty$, our model is equivalent to the model by Leroux *et al.* (*57*).

All data fitting was performed by numerically solving the full model shown in Fig. S12 using the ode45 solver implemented in Matlab® 2020b. For deriving relations between kinetic models with one and two bound state geometries (Fig. S17) used in sections 2.7 & 2.8, steady state analyses were performed to two simplified models (Fig. S13), as detailed in section 2.3.

The model used for numerical fitting (Fig. S12) assumes a single geometry (P-XY) for the hydrogen bound state, which forms in the different isotopomers available (P-HH, P-DH, P-HD, P-DD). Hydrogen association and dissociation proceed with rate constants $k_a$ and $k_d$, respectively, and hydrogen isotope exchange proceeds through $k_{HD}$, with the fraction of solvend deuteration $f_D$ as a known parameter. For the bound state, it is assumed that only one of the two positions in the hydrogen bound state (highlighted in red) is able to undergo hydrogen isotope exchange, and that mutual site exchange between the two bound positions within the bound intermediate can proceed via direct site exchange (rate constant $k_{ex}$) or via dissociation and reassociation. Kinetic isotope effects for the rate constants $k_a$, $k_d$, $k_{HD}$ and $k_{ex}$ are neglected for simplicity, relying on the previous observation, that at pH 6.0, the kinetic isotope effects are small for the isotope exchange reaction between hydrogen and water (*14*).

Hydrogen mass transport from the gas phase to the liquid phase was modeled with rate constants $k_{in}$ and $k_{out}$. To good approximation, mass transport between the liquid and the gas phase can be neglected during the kinetics measured, which were of < 10 min lenght. In the absence of bubbling, we measured $k_{out} = (0.084 \pm 0.006)\ h^{-1}$ and $k_{in} = (0.52 \pm 0.05)\ mM\ h^{-1}$.

From a single kinetic run according toFig. S1, the sample activity can be determined (see section 1.8), but only two linearly independent kinetics parameters can be fit. To be able to fit the three parameters $k_a$, $k_d$ and $k_{HD}$, we performed multiple kinetic runs, with variation of the total concentration of dissolved hydrogen $c_0 = [H_2] + [HD] + [D_2] \approx [H_2]_0$. To achieve this variation of $c_0$, we performed series of kinetic runs with varying mixtures of $n$-$H_2$ and $N_2$ in the range of 10% - 100% $n$-$H_2$. In all cases, bubbling was performed at 7 bars (gauge) of total pressure, to retain the same bubbling characteristics.

With this approach, it is possible to extract the rate constants $k_a$, $k_d$ and $k_{HD}$, if the limit $k_{ex} \rightarrow \infty$ is assumed. In the limit $k_{ex} \rightarrow \infty$, our model is equivalent to the model by Leroux *et al.* (*57*). Best fit parameters are listed in Table S4.

In the context of the PHIP effects studied, it is very useful in addition to discuss the limit of finite $k_{ex}$, since fast site exchange via $k_{ex}$ would quench both of the observed PHIP effects. While fitting of all four rate constants ($k_a$, $k_d$, $k_{HD}$ and $k_{ex}$) is not possible from our data, it is possible to estimate the lower bounds $k_{ex,min} = (90 \pm 10)\ s^{-1}$ and $k_{HD,min} = (178 \pm 13)\ s^{-1}$ which are compatible with our model (see also equation ( 23 ) in section 2.3.1), and to find best-fit solutions for $k_a$, $k_d$ and $k_{HD}$ for chosen values of $k_{ex}$. Over the whole range of $k_{ex}$ compatible with our data, best fit solutions for $k_a$ and $k_d$ deviate by less than 20%, whereas the best fit values for $k_{HD}$ show a strong dependence on the choice of $k_{ex}$, as expected from the steady state analysis.

Since for $k_{ex}$ and $k_{HD}$ only lower bound estimates can be extracted from the data, only an upper estimate for the bound state lifetime

$$\tau = (k_d + k_{HD} + k_{ex})^{-1} \qquad (8)$$

can be obtained in which isotope exchange occurs. Form the kinetic data measured at 309 K, we obtain $\tau_{max,isotope\ exchange} \approx 1.6\ ms$.

While isotope exchange kintetics are capable of providing an upper limit for these lifetimes, the PHIP experiments will further set a lower bound to these values, since both PHIP effects will vanish in the limit of short bound state lifetimes.

### 2.3 Steady state kinetics analysis of models with one and two bound state geometries

To derive approximate relations between the rate laws of the models with one or two bound state geometries (Fig. S13 A & B), we performed a steady-state analysis of the two kinetic models similar to the analysis in (*57*), which assumes a steady state for the bound intermediates ($[P]_0 \ll [H_2]_0$). For the analysis, it is further assumed, that the bubbling has already stopped ($k_{in} = 0$, $k_{out} = 0$), and that back reactions are negligible, due to a high degree of deuteration ($f_D = 1$). The model with two bound states (Fig. S13 B) further assumes that hydrogen isotope exchange only occurs in the bound states P-H-H and P-H-D ($k_{HD1} = 0$).

#### 2.3.1 Steady state analysis of model with one bound state geometry

Steady-state analysis of the model with one bound state geometry (Fig. S13 A) yields

$$\frac{d[H_2]}{dt} = -[P]_0 \frac{k_d k_{HD}}{k_d + k_{HD}} \frac{[H_2]}{c_0 + \frac{k_d}{k_a}} \quad (9)$$

from which the initial rate of $H_2$ consumption can be obtained.

$$\left.\frac{d[H_2]}{dt}\right|_{t=0} \approx -[P]_0 \frac{k_d k_{HD}}{k_d + k_{HD}} \frac{[H_2]_0}{[H_2]_0 + \frac{k_d}{k_a}} \quad (10)$$

Comparing to Menten equation

$$v_0 = v_{max} \frac{[S]}{[S] + K_m} \quad (11)$$

yields

$$\left.\frac{d[H_2]}{dt}\right|_{t=0} \approx -v_{max,1} \frac{[H_2]_0}{[H_2]_0 + K_m}, \quad (12)$$

with

$$K_{m,1} = \frac{k_d}{k_a} \quad (13)$$

and

$$v_{max,1} = k_a K_{m,1} [P]_0 \frac{1}{1 + \frac{k_d}{k_{HD}}}. \quad (14)$$

Note, that we are defining $v_0$ here by the initial rate of educt consumption, rather than by the initial rate of product formation.
Equations ( 13 ) and ( 14 ) were used to obtain the Menten parameters stated in Table 4.

The index $i = 1, 2$ was herein introduced to facilitate the discrimination between the analytical expressions for $K_{m,i}$ and $v_{max,i}$ referring to the model with one bound state geometry (Fig. S13A, $i = 1$) and the model with two bound state geometries (Fig. S13B, $i = 2$).

Further, we find in steady state:

$$[P] = [P]_0 \frac{K_m}{K_m + c_0} \quad (15)$$

Under the assumptions made, the integrated rate-law for $[H_2]$ is a simple monoexponential

$$[H_2](t) = [H_2]_0 e^{-At}, \quad (16)$$

with

$$A = [P]_0 \frac{k_a k_d k_{HD}}{(c_0 k_a + k_d)(k_d + k_{HD})} = k_a [P]_0 \frac{K_{m,1}}{c_0 + K_{m,1}} \frac{k_{HD}}{(k_d + k_{HD})} = \frac{v_{max,1}}{c_0 + K_{m,1}}. \quad (17)$$

Further introducing

$$B = \frac{1}{2}[P]_0 \frac{k_a k_d k_{HD}}{(c_0 k_a + k_d)\left[k_d + k_{HD}\frac{k_d + k_{ex}}{k_d + 2k_{ex}}\right]}$$
$$= \frac{1}{2} k_a [P]_0 \frac{K_{m,1}}{c_0 + K_{m,1}} \frac{k_{HD}}{\left[k_d + k_{HD}\frac{k_d + k_{ex}}{k_d + 2k_{ex}}\right]} \quad (18)$$

and

$$C = 2\left(1 - \frac{k_{ex} k_{HD}}{(k_d + k_{HD})(\frac{1}{2}k_d + k_{ex})}\right), \quad (19)$$

the rate law for $[HD]$ can be expressed as

$$\frac{d[HD]}{dt} = BC[H_2]_0 e^{-At} - B[HD] \quad (20)$$

from which

$$[HD](t) = 2[H_2]_0 (e^{-Bt} - e^{-At}). \quad (21)$$

(note: $BC/(A - B) = 2$).

The HD concentration peaks at

$$t_{max} = \frac{ln(A/B\ )}{A - B} \quad (22)$$

from which the ratio of the maximum HD concentration $[HD]_{max}$ and the initial $H_2$ concentration $[H_2]_0$ is obtained as

$$\frac{[HD]_{max}}{[H_2]_0} = 2\left(\left(\frac{A}{B}\right)^{-\frac{B}{A-B}} - \left(\frac{A}{B}\right)^{-\frac{A}{A-B}}\right), \qquad (23)$$

which is only depends on the ratios $k_d/k_{ex}$ and $k_d/k_{HD}$.

$$\frac{A}{B} = 2 - \frac{1}{\left(\frac{1}{2}k_d/k_{ex} + 1\right)(k_d/k_{HD} + 1)} \qquad (24)$$

$$\frac{B}{A-B} = \frac{(k_d/k_{HD} + 1)\left(\frac{1}{2}k_d/k_{ex} + 1\right)}{(k_d/k_{HD} + 1)\left(\frac{1}{2}k_d/k_{ex} + 1\right) - 1} \qquad (25)$$

$$\frac{A}{A-B} = \frac{2(k_d/k_{HD} + 1)\left(\frac{1}{2}k_d/k_{ex} + 1\right) - 1}{(k_d/k_{HD} + 1)\left(\frac{1}{2}k_d/k_{ex} + 1\right) - 1} \qquad (26)$$

The ratio $[HD]_{max}/[H_2]_0$ is experimentally easily accessible, and enables a straightforward upper bound estimate of the ratios $k_d/k_{HD}$ and $k_d/k_{ex}$. The contours for different ratios of $[HD]_{max}/[H_2]_0$ are shown in Fig. S15. Experimentally we observe $[HD]_{max}/[H_2]_0 \approx 0.1$ from which we obtain $k_d/k_{HD} \leq 0.2$ and $k_d/k_{ex} \leq 0.4$. With $k_d = (35 \pm 2)\ s^{-1}$, as obtained from fitting, we get $k_{HD,min} = (175 \pm 10)\ s^{-1}$ and $k_{ex,min} = (88 \pm 5)\ s^{-1}$.

2.3.2 <u>Steady state analysis of model with two bound state geometries</u>

Steady-state analysis of the model with two bound-state geometries (Fig. S13B) yields:

$$\frac{d[H_2]}{dt} = -[P]_0 k_{d1} \frac{k_{d2}}{k_{a2} + k_{d2}} \frac{1}{1 + \frac{k_{d1}}{k_{a2}}\left(1 + \frac{k_{d2}}{k_{HD2}}\right)} \frac{[H_2]}{c_0 + \frac{k_{d1}}{k_{a1}} \frac{k_{d2}}{k_{a2} + k_{d2}}} \qquad (27)$$

Again setting $c_0|_{t=0} \approx [H_2]_0$ and comparing to Menten equation ( 11 ) yields

$$\left.\frac{d[H_2]}{dt}\right|_{t=0} \approx -v_{max,2} \frac{[H_2]}{[H_2] + K_{m,2}}, \qquad (28)$$

with

$$K_{m,2} = \frac{k_{d1}}{k_{a1}} \frac{k_{d2}}{k_{a2} + k_{d2}} \qquad (29)$$

and

$$v_{max,2} = k_{a1} K_{m,2} [P]_0 \frac{1}{1 + \frac{k_{d1}}{k_{a2}}\left(1 + \frac{k_{d2}}{k_{HD2}}\right)} \qquad (30)$$

for this model.

In the steady state, we again find

$$[P] = [P]_0 \frac{K_m}{K_m + c_0} \tag{15}$$

and further we find

$$\frac{[\text{P-H-H}]}{[\text{P-HH}]} = \frac{k_{a2}}{k_{d2} + k_{HD2}} \tag{31}$$

and

$$\frac{[\text{P-H-H}]}{[\text{P-H-H}] + [\text{P-HH}]} = \frac{k_{a2}}{k_{a2} + k_{d2} + k_{HD2}}. \tag{32}$$

Again, we can express the rate laws in equations ( 16 ), ( 20 ) and ( 21 ) in terms of constants $A$, $B$ and $C$, which for the model with two bound states (Fig. S13B) take the form

$$A = k_{a1}[P]_0 \frac{K_{m,2}}{c_0 + K_{m,2}} \frac{1}{1 + \frac{k_{d1}}{k_{a2}}\left(1 + \frac{k_{d2}}{k_{HD2}}\right)} = \frac{v_{max,2}}{c_0 + K_{m,2}}, \tag{33}$$

$$B = \frac{1}{2} k_{a1} k_{a2} [P]_0 \frac{K_{m,2}}{c_0 + K_{m,2}}$$

$$\cdot \frac{k_{HD2}}{\left[k_{d1}k_{d2} + \frac{k_{HD2}\left(k_{d1}(k_{d1} + 2k_{ex1})(k_{d2} + k_{ex2}) + k_{a2}\left((k_{d1} + k_{ex1})(k_{d2} + 2k_{ex2}) + k_{a2}k_{ex2}\right)\right)}{\left((k_{d1} + 2k_{ex1})(k_{d2} + 2k_{ex2}) + 2k_{a2}k_{ex2}\right)}\right]} \tag{34}$$

and

$$C = \frac{k_{d1}k_{d2}}{k_{d1}(k_{d2} + k_{HD2}) + k_{a2}k_{HD2}}$$

$$\cdot \left(\frac{(k_{d1} + 2k_{ex1})\left((k_{d2} + 2k_{ex2}) + k_{HD2}\right) + k_{a2}(2k_{ex2} + k_{HD2})}{(k_{d1} + 2k_{ex1})\left(\frac{1}{2}k_{d2} + k_{ex2}\right) + k_{a2}k_{ex2}}\right). \tag{35}$$

In the limit $k_{d1} \gg k_{a2}$ discussed later (section 2.7), these relations simplify to

$$A|_{k_{d1} \gg k_{a2}} \approx \frac{k_{a1}k_{a2}}{k_{d1}} [P]_0 \frac{K_{m,2}}{c_0 + K_{m,2}} \frac{k_{HD2}}{k_{HD2} + k_{d2}}, \tag{36}$$

$$B|_{k_{d1} \gg k_{a2}} \approx \frac{1}{2} \frac{k_{a1}k_{a2}}{k_{d1}} [P]_0 \frac{K_{m,2}}{c_0 + K_{m,2}} \frac{k_{HD2}}{\left[k_{d2} + k_{HD2} \frac{(k_{d2} + k_{ex2})}{(k_{d2} + 2k_{ex2})}\right]} \tag{37}$$

and

$$C|_{k_{d1}\gg k_{a2}} \approx 2\left(1-\frac{k_{ex2}k_{HD2}}{(k_{d2}+k_{HD2})\left(\frac{1}{2}k_{d2}+k_{ex2}\right)}\right). \qquad (38)$$

Comparing to equations ( 17 ) - ( 19 ) shows that in the limit $k_{d1} \gg k_{a2}$, the net kinetics of the model with two bound state geometries (Fig. S13B) behave like those of the model with one bound state geometry (Fig. S13A), with an effective association constant

$$k_a|_{k_{d1}\gg k_{a2}} \approx \frac{k_{a1}k_{a2}}{k_{d1}}, \qquad (39)$$

and with

$$k_{d2}|_{k_{d1}\gg k_{a2}} \approx k_d, \qquad (40)$$

$$k_{HD2}|_{k_{d1}\gg k_{a2}} \approx k_{HD} \qquad (41)$$

and

$$k_{ex2}|_{k_{d1}\gg k_{a2}} \approx k_{ex}. \qquad (42)$$

### 2.4 An analytical Model for the HD-PHIP

We attribute the HD-PHIP effects observed to simple coherent evolution of the initial singlet state of $p$-$H_2$ under the effects of a chemical shift difference and a $J$-coupling in the bound state. From the spin dynamics perspective, this effect is closely related to the oneH-PHIP (*58*) and the NEPTUN effect (*33, 34, 59*).

The simplest mechanistic model in which this effect can be understood is shown in Fig. S16A. In this model, dissolved $p$-$H_2$ can reversibly bind to a catalytic site, and it can undergo hydrogen isotope exchange at catalytic site, to form HD. In the bound state, the two atoms stemming from $p$-$H_2$ experience a difference of their chemical shifts $\Delta\delta = \delta_{\mathrm{I}} - \delta_{\mathrm{S}}$ and a non-zero mutual $J$-coupling $J_{HH} \neq 0$. Spin evolution in the bound state builds up magnetization on the two nuclei in an $\frac{1}{2}(\hat{I}_z - \hat{S}_z)$ term, as detailed below. This magnetization becomes observable on HD and $H^+$ after $H^+ \rightarrow D^+$ exchange and subsequent release from the enzyme.

For the mechanism to produce observable magnetization it is required, that one of the two binding sites $I$ and $S$, has a higher propensity for $H^+ \rightarrow D^+$ exchange. For simplicity we assume, that $H^+ \rightarrow D^+$ exchange only occurs at site $S$ (highlighted in red), so that it is the $\hat{S}_z$ term that will produce hyperpolarized $H^+$ and the $\hat{I}_z$ term, that will produce hyperpolarized HD.

Spin evolution from the singlet state of $p$-$H_2$ into the magnetization observable later on $H^+$ and HD happens in the bound state P-HH. The relevant evolution of the singlet state under the action of $\Delta\delta$ and $J_{HH}$ in an asymmetric bound state has been reviewed by Natterer and Bargon (equations 7-18 of the cited review) (*28*) and will be briefly recaptured in equations ( 44 ) – ( 60 ). We will extend the discussion to cases, in which this bound state only exists during a short binding period:

When $p$-$H_2$ undergoes a reaction forming a reaction product in which the two hydrogen atoms have non-zero chemical shift difference and $J$-coupling, the singlet state initially present

$$\hat{\rho}_0 = |S_0\rangle\langle S_0| = \frac{1}{2}(|\alpha\beta\rangle - |\beta\alpha\rangle)(\langle\alpha\beta| - \langle\beta\alpha|) = \frac{1}{4}\mathbb{1} - \hat{I}_x\hat{S}_x - \hat{I}_y\hat{S}_y - \hat{I}_z\hat{S}_z \quad (43)$$

starts evolving under the action of $\Delta\delta$ and $J_{HH}$. If the system is treated as isolated 2-spin-1/2 system in isotropic solution and neglecting relaxation, the Hamiltonian can be split into two terms, of which only $\hat{H}_1$ determines the spin evolution of $|S_0\rangle\langle S_0|$ (*28*).

$$\hat{H} = \hat{H}_o + \hat{H}_1 \quad (44)$$

with

$$\hat{H}_0 = \pi\, \nu_\Sigma(\hat{I}_z + \hat{S}_z) + 2\pi\, J_{IS}\hat{I}_z\hat{S}_z \quad (45)$$

and

$$\hat{H}_1 = \pi\, \Delta\nu(\hat{I}_z - \hat{S}_z) + 2\pi\, J_{IS}(\hat{I}_x\hat{S}_x + \hat{I}_y\hat{S}_y) \quad (46)$$

and with the frequencies

$$\nu_\Sigma = \nu_I + \nu_S \quad (47)$$

and

$$\Delta\nu = \nu_I - \nu_S, \quad (48)$$

and with

$$\nu_{\mathrm{i}} = -\frac{\gamma B_0}{2\pi}(1+\delta_i) \qquad (i = I, S). \tag{49}$$

Evolution of $\hat{\rho}_0 = |S_0\rangle\langle S_0|$ over a time period $t$ under the action of $\widehat{H}_1$, yields the density operator

$$\hat{\rho}(t) = \tfrac{1}{4}\hat{\mathbb{1}} - \hat{I}_z\hat{S}_z - \mathrm{a}(t)\widehat{ZQ}_x - \mathrm{b}(t)\widehat{ZQ}_y - \mathrm{c}(t)\tfrac{1}{2}\left(\hat{I}_z - \hat{S}_z\right) \tag{50}$$

for the bound state, with the zero-quantum terms

$$\widehat{ZQ}_x = \hat{I}_x\hat{S}_x + \hat{I}_y\hat{S}_y \tag{51}$$

and

$$\widehat{ZQ}_y = \hat{I}_y\hat{S}_x - \hat{I}_x\hat{S}_y. \tag{52}$$

The coefficients $\mathrm{a}(t)$, $\mathrm{b}(t)$ and $\mathrm{c}(t)$ hereby are given by (*28*)

$$a(t) = \frac{1}{\xi^2+1}\left(1+\xi^2\cos(kt)\right) \tag{53}$$

$$b(t) = \frac{\xi}{\sqrt{\xi^2+1}}\sin(kt) \tag{54}$$

$$c(t) = \frac{\xi}{\xi^2+1}\left(1-\cos(kt)\right) \tag{55}$$

with

$$k = 2\pi\sqrt{(\nu_I - \nu_S)^2 + J_{HH}^2} \tag{56}$$

and

$$\xi = \frac{\nu_I - \nu_S}{J_{HH}}. \tag{57}$$

To obtain the observable density operator, we have to perform appropriate averaging of the coefficients $\mathrm{a}(t)$, $\mathrm{b}(t)$ and $\mathrm{c}(t)$ over the reaction duration.

The scenario usually discussed is a one-way addition reaction to a substrate P

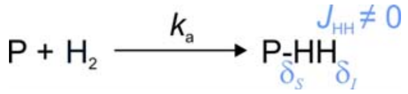


( 58 )

which is slow relative to spin evolution ($k \gg k_a[S]$). In this case, the time dependent factors average to zero and one obtains (*28*)

$$\bar{\rho}_{slow} = \tfrac{1}{4}\hat{\mathbb{1}} - \hat{I}_z\hat{S}_z - \frac{1}{1+\xi^2}\widehat{ZQ}_x - \frac{\xi}{1+\xi^2}\tfrac{1}{2}\left(\hat{I}_z - \hat{S}_z\right), \tag{59}$$

which shows the well-known $B_0$-dependence of the $\tfrac{1}{2}\left(\hat{I}_z - \hat{S}_z\right)$ operator amplitude

$$\boldsymbol{Tr}\left(\tfrac{1}{2}(\hat{I}_z - \hat{S}_z)\bar{\rho}_{slow}\right) = -\bar{c}_{slow} = -\frac{\xi}{1+\xi^2}. \tag{60}$$

If, however, we consider the case of transient catalyst binding that happens at rates comparable to $k$, we need to reevaluate the ensemble averages of $\mathrm{a}(t)$, $\mathrm{b}(t)$ and $\mathrm{c}(t)$. To do so, we consider a bound state P-HH of characteristic lifetime $\tau$, during which the singlet state can evolve under the action of $\widehat{H}_1$. Hereby, we make the following assumptions:

1. We assume that the hydrogen forming P-HH is in a pure singlet state and that the density matrix $\hat{\rho}_{\text{P-HH}}(0)$ is given by equation ( 43 ). This implies the assumptions, that
   a. The hydrogen added enriched to 100% in $p$-$H_2$.
   b. Bubbling is sufficiently fast, that right after bubbling, the solution is saturated in pure $p$-$H_2$. This applies for $k_{out} = k_{in}/(p(p\text{-H}_2)H^{cp}) \gg k_a[P]$, where $k_{in}$ is the rate of $p$-$H_2$ inflow (mM/s), $k_{\text{out}}$ is the rate of $p$-$H_2$ exhaust ($s^{-1}$), $H^{cp}$ is Henry's constant (mM/bar) and $p(p\text{-H}_2)$ is the $p$-$H_2$ partial pressure applied during a bubbling (bar).
   c. We are only considering timepoints, where the singlet-to-triplet conversion is negligible and where hydrogen isotope exchange can be neglected. This is the case for timepoints with $k_a[P]t \ll 1$, where $t$ is the time measured from the end of the bubbling period.
2. We neglect possible mutual site exchange between the two hydrogens in the bound state P-HH ($k_{ex} = 0$).
3. We treat relaxation in the bound state phenomenologically by introducing a uniform relaxation rate $R_{\text{P-HH}}$ for all coherences in the bound state P-HH and we neglect cross-relaxation. Further it is assumed, that relaxation drives the system to $\rho(\infty) = 0$, rather than to thermal equilibrium.

The inverse of this characteristic lifetime is the sum of all first order reaction rates leading to loss of P-HH, which in the model given in Fig. S16A equates to

$$\tau^{-1} = k_\Sigma = k_d + k_{cat} \qquad (assuming\ k_{ex} = 0). \tag{61}$$

Assuming only first order reactions of loss, the probability density for P-HH to have a given lifetime $t$ is

$$p(t) = k_\Sigma e^{-k_\Sigma t}. \tag{62}$$

Ensemble averaging for $a(t)$ over the bound state lifetime via

$$\bar{a}_{transient} = \frac{\int_0^\infty p(t)a(t)e^{-R_{\text{P-HH}}t}\,dt}{\int_0^\infty p(t)\,dt} = \frac{\int_0^\infty p(t)a(t)e^{-R_{\text{P-HH}}t}\,dt}{\int_0^\infty p(t)\,dt} \tag{63}$$

yields

$$\bar{a}_{transient} = \frac{1}{1+\xi^2}\left[\frac{k_\Sigma}{k_\Sigma + R_{\text{P-HH}}} + \xi^2\frac{k_\Sigma(k_\Sigma + R_{\text{P-HH}})}{(k_\Sigma + R_{\text{P-HH}})^2 + k^2}\right], \tag{64}$$

and similarily, we obtain

$$\bar{b}_{transient} = \frac{\xi}{\sqrt{1+\xi^2}} \frac{k_\Sigma k}{(k_\Sigma + R_{\text{P-HH}})^2 + k^2}. \tag{65}$$

and

$$\bar{c}_{transient} = \frac{\xi}{1+\xi^2} \frac{k_\Sigma}{k_\Sigma + R_{\text{P-HH}}} \frac{k^2}{(k_\Sigma + R_{\text{P-HH}})^2 + k^2}. \tag{66}$$

The field dependence of $\bar{c}$ is thus modified in the case of transient binding, as compared to eq. ( 60 ). In addition to the term $\xi/(1+\xi^2)$ already contained in eq. ( 60 ), there is an additional term $k^2/[(k_\Sigma + R_{\text{P-HH}})^2 + k^2]$, which shifts the maximum of $\bar{c}$ towards higher fields and decreases the maximum value for $\bar{c}$, as soon as the bound-state lifetime $k_\Sigma^{-1}$ or the relaxation rate in the bound state $R_{\text{P-HH}}$ are comparable to the spin-state evolution rate $k$. From the shape of the field dependence of $\bar{c}$, the case of transient binding cannot be discriminated from the case of a slow one-way reaction, since in both cases a dispersive lorentzian profile is observed. Inserting ( 56 ) and ( 57 ) into ( 60 ) or ( 66 ), we obtain

$$\bar{c}_{slow} = \frac{(2\pi)^2 J_{HH}\,\Delta\nu}{(2\pi\,J_{HH})^2 + (2\pi\,\Delta\nu)^2}, \tag{67}$$

whereas

$$\bar{c}_{transient} = \frac{k_\Sigma}{k_\Sigma + R_{\text{P-HH}}} \frac{(2\pi)^2 J_{HH}\,\Delta\nu}{(k_\Sigma + R_{\text{P-HH}})^2 + (2\pi\,J_{HH})^2 + (2\pi\,\Delta\nu)^2}, \tag{68}$$

where only $\Delta\nu$ is field dependent. The expression for the magnetic field at which $\bar{c}_{transient}$ is maximized thus takes the simple form:

$$B_{\max,\bar{c}} = \sqrt{\frac{(k_\Sigma + R_{\text{P-HH}})^2 + (2\pi J_{HH})^2}{\gamma^2(\delta_I - \delta_S)^2}}. \tag{69}$$

Exemplary field profiles for $\bar{c}_{transient}$ are shown in Fig. S18A.

To estimate an upper-bound limit for the amplitude of the HD-PHIP effect, we assume a time-averaged density operator $\bar{\hat{\rho}}_{\text{P-HH}}$ for the bound state intermediate P-HH of

$$\lim_{t\to 0} \bar{\hat{\rho}}_{\text{P-HH}} \approx \tfrac{1}{4}\hat{\mathbb{1}} - \hat{I}_z\hat{S}_z - \bar{a}_{transient}\widehat{ZQ}_x - \bar{b}_{transient}\widehat{ZQ}_y - \bar{c}_{transient}\tfrac{1}{2}(\hat{I}_z - \hat{S}_z). \tag{70}$$

This assumption is only valid for early timepoints, where singlet-to-triplet conversion as well as hydrogen isotope exchange can be neglected ($k_a[P]t \ll 1$).

Isotope exchange leading to the formation of P-DH will quench the $\hat{I}_z\hat{S}_z$ term, the $\widehat{ZQ}_x$ term and the $\widehat{ZQ}_y$ term, whereas the longitudinal magnetization terms $\hat{I}_z$ and $\hat{S}_z$ remain on the reaction products after isotope exchange. With the hydrogen site that undergoes $H^+ \rightarrow D^+$ exchange in P-HH labeled $S$ (as stated above), only the $\hat{I}_z$ term will be transferred to HD. Following the kinetic scheme Fig. S16A, and again using a simplified treatment of relaxation, we approximate the time evolution of $\hat{\rho}_{\text{HD}}$ as

$$\dot{\hat{\rho}}_{\text{HD}} \approx -\left(i\hat{\hat{H}}_{\text{HD}} + R_{\text{HD}}\right)\hat{\rho}_{\text{HD}} + k_{cat}\boldsymbol{Tr}\left(\hat{I}_z\bar{\hat{\rho}}_{\text{P-HH}}\right)\hat{I}_z \tag{71}$$

For the "high-field" limit we are interested in ($B_0 \geq 1$ mT), the steady-state solution for $\hat{\rho}_{\text{HD}}$ is easily found (setting $\dot{\hat{\rho}}_{\text{HD}} \approx 0$), since to good approximation, $\hat{I}_z$ is an eigenstate of $\hat{\hat{H}}_{\text{HD}}$ in this field range.

$$\hat{\rho}_{\text{HD}} \approx \frac{k_{cat}}{R_{\text{HD}}} \boldsymbol{Tr}\left(\hat{I}_z \bar{\hat{\rho}}_{\text{P-HH}}\right) \hat{I}_z \tag{72}$$

In the limit, where the hydrogen forming P-HH mostly is *p*-H2, we thus find

$$\lim_{t \to 0} \hat{\rho}_{\text{HD}} \approx -K_{S19D} \frac{1}{2} \bar{c}_{transient} \hat{I}_z \tag{73}$$

with

$$K_{S19D} = \frac{k_{cat}}{R_{\text{HD}}}, \tag{74}$$

where the constants that depend on the details of the kinetic model assumed are condensed into a field-independent constant $K_{S19D}$, whereas the field-dependence of the HD-PHIP is contained in $\bar{c}_{transient}$ only. The polarization $P(HD)$ of HD expected in this model can be obtained from the expectation value of $\hat{I}_z$ as

$$\lim_{t \to 0} P(HD) = \boldsymbol{Tr}\left(\hat{I}_z \hat{\rho}_{\text{HD}}\right) \approx -K_{S19D} \frac{1}{2} \bar{c}_{transient} \tag{75}$$

In models in which coherence evolution through $\nu_I - \nu_S$ and $J_{HH}$ takes place in the same intermediate where isotope exchange is taking place, as is the case in Fig. S16A, the sign of the HD-PHIP polarization directly reflects the the sign of $\xi$, and thus allows to relate the sign of $J_{HH}$ to the position of HD-exchange. With the position of preferential $H^+ \to D^+$ exchange defined as *S*, and noting equations ( 49 ), ( 57 ), ( 66 ) & ( 80 ), one easily finds

$$\begin{aligned} \text{Sign(HD-PHIP)} &= \text{Sign}\left(-\frac{\nu_{\text{I}} - \nu_{\text{S}}}{J_{IS}}\right) = \text{Sign}\left(\frac{\gamma_H B_0(\delta_I - \delta_S)}{J_{IS}}\right) \\ &= \text{Sign}\left(\frac{\gamma_H B_0\left(\delta_{not\ exchanging} - \delta_{exchanging}\right)}{J_{IS}}\right). \end{aligned} \tag{76}$$

Since $\gamma_H > 0$ and $B_0 > 0$, for protons we can summarize

| | $J_{IS} > 0$ | $J_{IS} < 0$ |
|---|---|---|
| exchange at the high chemical shift position ($\delta_I - \delta_S < 0$) | negative HD-PHIP | positive HD-PHIP |
| exchange at the low chemical shift position ($\delta_I - \delta_S > 0$) | positive HD-PHIP | negative HD-PHIP |

Assuming a model in which coherence evolution through $\nu_I - \nu_S$ and $J_{HH}$ takes place in the same intermediate where isotope exchange is taking place, and noting that we experimentally observe positive hyperpolarization for HD, we would thus conclude that we either have $J_{IS} > 0$ and isotope exchange at the low chemical shift position, or $J_{IS} < 0$ and isotope exchange at the high chemical shift position. As outlined in the main article, however our data is incompatible with models assuming only one bound state, and thus the rules mentioned here do not necessarily apply to our case, since $\Delta\delta = \delta_I - \delta_S$ can switch sign in-between different bound intermediates. The model outlined in Fig. 3A of the main article assumes such a switch of sign for $\Delta\delta$.

For fitting the field-cycling data discussed in section 2.5, we used the kinetic model in Fig. S16B, which comes closer to the full kinetic model used for extracting rate constants (Fig. S12). The model shown in Fig. S16B in itself contains an inconsistency, which we accepted for simpler treatment: For P-HH we assume that no mutual site exchange is taking place ($k_{ex} = 0$), whereas for the net rate from P-DH to P-DD, we assume a value of $\frac{1}{2}k_{HD}$, which is the limit of fast mutual exchange. For rigorous treatment of exchange in P-HH, analytical expressions become so lengthy that numerical simulations are preferable.

Following the kinetic scheme Fig. S16B, and once again using a simplified treatment of relaxation, we approximate the time evolution of $\hat{\rho}_{\text{P-DH}}$ and $\hat{\rho}_{\text{HD}}$ as

$$\dot{\hat{\rho}}_{\text{P-DH}} \approx -\left(i\hat{\hat{H}}_{\text{P-DH}} + R_{\text{P-DH}} + k_d + \tfrac{1}{2}k_{HD}\right)\hat{\rho}_{\text{P-DH}} + k_a[P]\hat{\rho}_{\text{HD}} + k_{HD}\boldsymbol{Tr}\left(\hat{I}_z\bar{\hat{\rho}}_{\text{P-HH}}\right)\hat{I}_z \qquad (77)$$

$$\dot{\hat{\rho}}_{\text{HD}} \approx -\left(i\hat{\hat{H}}_{\text{HD}} + R_{\text{HD}} + k_a[P]\right)\hat{\rho}_{\text{HD}} + k_d\hat{\rho}_{\text{P-DH}}. \qquad (78)$$

For the “high-field” limit ($B_0 \geq 1$ mT), $\hat{I}_z$ again is an eigenstate of $\hat{\hat{H}}_{\text{P-DH}}$ and $\hat{\hat{H}}_{\text{HD}}$, thus the steady-state solutions for $\hat{\rho}_{\text{P-DH}}$ and $\hat{\rho}_{\text{HD}}$ are easily found as well (setting $\dot{\hat{\rho}}_{\text{P-DH}} \approx 0$ and $\dot{\hat{\rho}}_{\text{HD}} \approx 0$).

$$\hat{\rho}_{\text{P-DH}} \approx \frac{k_a[P]\hat{\rho}_{\text{HD}} + k_{HD}\boldsymbol{Tr}\left(\hat{I}_z\bar{\hat{\rho}}_{\text{P-HH}}\right)\hat{I}_z}{R_{\text{P-DH}} + k_d + \frac{1}{2}k_{HD}} \qquad (79)$$

$$\hat{\rho}_{\text{HD}} \approx \frac{k_d}{(R_{\text{HD}} + k_a[P])}\hat{\rho}_{\text{P-DH}}, \qquad (80)$$

In the limit where $p$-$H_2$ makes up for nearly all $H_2$ dissolved ($k_a[P]t \ll 1$, full enrichment after bubbling) we obtain

$$\lim_{t\to 0}\hat{\rho}_{\text{HD}} \approx -K_{S19E}\frac{1}{2}\bar{c}_{transient}\hat{I}_z. \qquad (81)$$

where in this case the constant $K$ containing the details of the kinetic model is

$$K_{S19E} = \frac{k_d k_{HD}}{(R_{\text{HD}} + k_a[P])\left(R_{\text{P-DH}} + \frac{1}{2}k_{HD}\right) + k_d R_{\text{HD}}}. \qquad (82)$$

From this, the polarization $P(HD)$ of HD at early timepoints during the kinetics can be obtained from the expectation value of $\hat{I}_z$ as

$$\lim_{t\to 0}P(HD) = \lim_{t\to 0}\boldsymbol{Tr}\left(\hat{I}_z\hat{\rho}_{\text{HD}}\right) \approx -\frac{1}{2}K_{S19E}\frac{\xi}{1+\xi^2}\frac{k_\Sigma}{k_\Sigma + R_{\text{P-HH}}}\frac{k^2}{(k_\Sigma + R_{\text{P-HH}})^2 + k^2} \qquad (83)$$

Estimating the HD-concentration from eqs. ( 17 ), ( 18 ) & ( 21 ) for the limit $k_{ex} \to \infty$

$$[HD](t) = 2[H_2]_0(e^{-Bt} - e^{-At}). \quad (21)$$

$$\lim_{t\to 0}[HD](t) \approx \frac{[H_2]_0 k_a k_d k_{HD}}{[H_2]_0 k_a + k_d} \frac{k_{HD}}{\left(k_d + \frac{1}{2}k_{HD}\right)(k_d + k_{HD})} t, \quad (84)$$

the experimentally observed signal integral $[HD]\ P(HD)$ can be estimated from the HD signal integral $I(HD)$ and the integral $I(ref)$ of a thermally polarized signal of reference with known concentration.

$$[HD]\ P(HD) = [ref]\ P_0(ref)\frac{I(HD)}{I(ref)} \quad (85)$$

with

$$P_0(ref) \approx \frac{\gamma_k \hbar B_0}{2k_B T} \quad (4)$$

for spin-1/2 nuclei.

2.5 Field cycling experiments

To characterize the origin of the HD-PHIP, we performed field cycling experiments with manual shuttling, as described in section 1.11 according to the scheme shown in Fig. S4. In brief, samples were bubbled at low-field positions, realized by lifting the sample to specific positions along the magnet bore prior to bubbling, and readout of the HD polarization produced at different fields was accomplished by manually shuttling the sample into the probe after bubbling had stopped, where the HD polarization produced was read out by a pulse acquisition scheme. Since no active temperature control was available at the low field positions, all experiments were performed at room temperature. Performing experiments at room temperature also ensured, that the limit of small $p$-$H_2$ consumption/conversion ($k_a[P]t \ll 1$) assumed in the model outlined in section 2.4 was well fulfilled during the experiments.

The HD-PHIP polarization produced (Fig. S18B) has a defined maximum at $B_{max} = (1.9 \pm 0.2)\ T$ and shows the field dependence expected from the model outlined in section 2.4 (i.e. dispersive lorentzian shape, see equation ( 68 )). This suggests that coherent spin evolution under strong coupling in a transiently formed intermediate is the dominant mechanism for creation of the HD-PHIP over the field range sampled (1 mT – 7 T).

To fit the HD-PHIP signal intensity experimentally observed, rate constants for the net isotope exchange reaction were determined for the experiment temperature ($T = 292\ K$), assuming the kinetic model shown in Fig. S12 with the limit $k_{ex} \to \infty$. We obtained $k_a = (7 \pm 4)\ mM^{-1}s^{-1}, k_d = (4.6 \pm 0.3)\ s^{-1}$ and $k_{HD} = (35 \pm 5)\ s^{-1}$ (lower bound estimate for $k_{HD}$), and in addition we obtain $[H_2]_0 \approx (5.4 \pm 1.2)\ mM$. From these parameters, we get a rough estimate of the HD concentration $[HD](2\ s) \approx 18 \pm 6\ \mu M$, according to equation ( 85 ). With this concentration estimate, the maximum polarization for HD can be estimated as $P(HD, 1.9\ T) = (1.0 \pm 0.6)\%$, corresponding to an enhancement of $\varepsilon = (410 \pm 250)$ over the thermal polarization at 7 T.

A fit of the experimentally observed HD-PHIP intensity was performed according to equation ( 83 ) using the above-mentioned kinetic parameters, and assuming $R_{HD} = (8\ s)^{-1}$, $R_{P-HH} = (1\ s)^{-1}$ and $R_{P-DH} = (8\ s)^{-1}$. We obtain $\Delta\delta = \mp(0.080 \pm 0.014)\ ppm$ and $J_{HH} = \pm(0.005 \pm 0.001)\ Hz$. The small magnitude of these fit results most likely is due to an overestimation of the lifetime of the bound species producing the HD-PHIP. Such an overestimation appears reasonable, since it may have several origins:

1. The analysis of the overall isotope exchange kinetics according to the model in Fig. S12 assumes a single bound species. The true catalytic mechanism must however be comprised of multiple catalytic steps, which may have significantly shorter lifetime.
2. As outlined in section 2.3.1, analysis of the overall isotope exchange kinetics only enables lower limit estimates for $k_{HD}$ and $k_{ex}$. Since $k_{HD}$ limits the lifetime of the bound species, an overestimation of its lifetime is clearly possible.
3. For simpler analytic treatment, we neglected the effects of mutual site exchange via $k_{ex}$ for the bound state.

In addition, the relaxation rates in the bound states may also have been underestimated. Our analysis thus supports the model of coherent spin evolution in the bound state as the cause of the HD-PHIP, and it suggests that the species in which spin evolution is taking place likely is shorter lived than the upper-bound bound state lifetime estimated from the overall isotope exchange kinetics.

2.6 Spin Dynamics Models for Numerical Simulations

Numerical spin dynamics simulations were performed using three different kinetic networks shown in Fig. S17:

- **Model-1b** (Fig. S17A): a model that assumes a single bound state (1b) without isotope exchange:
- **Model-1b-HDex** (Fig. S17B): a model assuming a single bound state (1b) where a hydrogen isotope exchange can take place (HDex): $H_2 \rightleftarrows HD \rightleftarrows D_2$.
- **Model-2b-2HDex** (Fig. S17C): a model assuming two different bound states (2b). Each such state can result in a hydrogen isotope exchange (2HDex).

Each model is an extension of the previous model, therefore we will describe them sequentially increasing complexity.

To calculate the spin state of the exchanging system we will use an approach described before and used to simulate Signal Amplification By Reversible Exchange of parahydrogen (SABRE) (*43, 60*). The master equation of this method is

$$\frac{d\hat{\rho}}{dt} = \left(\hat{\hat{L}} + \hat{\hat{K}}\right)\hat{\rho} + \widehat{W}, \tag{86}$$

where $\hat{\rho}$ is the generalized density matrix, $\hat{\hat{L}} = -i\hat{\hat{H}} + \hat{\hat{R}}$ is generalized Liouvillian superoperator that includes Hamiltonian superoperators, $\hat{\hat{H}} = [\widehat{H},\cdot]$ and relaxation superoperator $\hat{\hat{R}}$ of all exchanging intermediates. We used only a local fluctuating magnetic field relaxation mechanism for all species except $H_2$. The step-by-step description of computation of the local fluctuating field mechanism is given in Ref. (*61*) and needs only a single parameter of $T_1$ relaxation time ($T_1$). To describe the relaxation of $H_2$ we used a combination of local fluctuating fields with an intramolecular dipolar-dipole relaxation as was proposed before (*62*) to take into account the long lifetime of para-ortho conversion (POC). Two parameters are needed: $T_1$ and $T_{POC}$.

Each intermediate in our case was consisting only of two spins, however, the same approach was used for more spins e.g. in Ref. (*63*), and it is straightforward to extend it to more spins. $\widehat{W}$ is the source of spin order or molecules. Below we will detail the structure of these operators for the used models.

**Model-1b.** There are two chemical species in this model: $H_2$ in a solution and bound species P-HH with two protons A and B. The used NMR parameters are:

$H_2$: ($\delta$ = 4.655 ppm, $T_1$ = 5 s, $T_{POC}$ = 500 s, $J_{HH}$ = 280 Hz).

P-HH: ($\delta_A$, $\delta_B$, $\Delta\delta = \delta_B - \delta_A$, $T_1$ = 1 s, $J_{HH}$).

If the parameters are not specified then they were varied and are given where necessary. The necessary density matrices and superoperators of this model have the following structure:

$$\hat{\rho} = \begin{pmatrix} \hat{\rho}_{HH} \\ \hat{\rho}_{P-HH} \end{pmatrix}, \tag{87}$$

$$\hat{\hat{L}} = \begin{pmatrix} \hat{\hat{L}}_{HH} & 0 \\ 0 & \hat{\hat{L}}_{P-HH} \end{pmatrix}, \tag{88}$$

$$\hat{\hat{K}} = \begin{pmatrix} -(k_a' + k_{out})\hat{\hat{1}}_{HH} & k_d\hat{\hat{1}}_{HH} \\ 0.5k_a'\left(\hat{\hat{1}}_{HH} + \hat{\hat{Q}}_{H1\leftrightarrow H2}\right) & -k_d\hat{\hat{1}}_{HH} \end{pmatrix}, \tag{89}$$

$$\hat{W} = \begin{pmatrix} k_{in}\hat{\rho}_{\mathrm{HH}^*} \\ 0 \end{pmatrix}, \quad (90)$$

where all "0" in the equation are assumed to be zero matrices or vectors of the proper size.

The density matrix of 100% p-$H_2$ written in a square form is

$$\hat{\rho}_{\mathrm{HH}^*} = \frac{1}{2}\begin{pmatrix} 0 & 0 & 0 & 0 \\ 0 & 1 & -1 & 0 \\ 0 & -1 & 1 & 0 \\ 0 & 0 & 0 & 0 \end{pmatrix}, \quad (91)$$

the unitary operator is

$$\hat{\hat{1}}_{\mathrm{HH}} = \hat{1}_{\mathrm{HH}} \otimes \hat{1}_{\mathrm{HH}}, \text{ with } \hat{1}_{\mathrm{HH}} = \begin{pmatrix} 1 & 0 & 0 & 0 \\ 0 & 1 & 0 & 0 \\ 0 & 0 & 1 & 0 \\ 0 & 0 & 0 & 1 \end{pmatrix} \quad (92)$$

And the operator of exchange of two spin-1/2 nuclei is

$$\hat{\hat{Q}}_{\mathrm{H1}\leftrightarrow\mathrm{H2}} = \hat{Q}_{\mathrm{H1}\leftrightarrow\mathrm{H2}} \otimes \hat{Q}_{\mathrm{H1}\leftrightarrow\mathrm{H2}}, \text{ with } \hat{Q}_{\mathrm{H1}\leftrightarrow\mathrm{H2}} = \begin{pmatrix} 1 & 0 & 0 & 0 \\ 0 & 0 & 1 & 0 \\ 0 & 1 & 0 & 0 \\ 0 & 0 & 0 & 1 \end{pmatrix}. \quad (93)$$

Association is treated as pseudo-first-order process with the pseudo-first-order association rate $k'_a = k_a[P]$ with the units s$^{-1}$, wherein $[P]$ is the steady-state concentration value (see equation ( **15** )) and $k_a$ is an association rate. $k_d$ is the dissociation rate (s$^{-1}$), $k_{in}$ is the rate of $p$-$H_2$ inflow (mM/s) and $k_{\mathrm{out}}$ is the rate of $p$-$H_2$ exhaust (s$^{-1}$).

Initial conditions: all simulations were started from $\hat{\rho} = 0$ and two time evolution periods were explicitly simulated: A bubbling period, during which hydrogen mass transport between gas-phase and liquid phase proceed through $k_{in}$ and $k_{out}$ and the period after bubbling during mass transport between the gas and the liquid phase is assumed to be negligible ($k_{in} = 0$ and $k_{out} = 0$). During bubbling, we generally assumed $k_{in} = 10$ mM/s and $k_{out} = k_{in}/p(p\text{-H}_2)H^{cp} \approx 2s^{-1}$, $H^{cp}$ is Henry's constant and $p(p\text{-H}_2)$ is the $p$-$H_2$ partial pressure applied during a bubbling.

**Model-1b-HDex.** This is the extension of the Model-1b with an isotopic exchange in the bound form. In the bound states, hydrogen isotope exchange is assumed to only occur for the A-protons which have a chemical shift $\delta_{\mathrm{A}}$. The isotope exchange rates are $k_{\mathrm{HD}}$ (position highlighted in red). The second B-proton with chemical shift $\delta_{\mathrm{B}}$ does not participate in the HD exchange. We define the chemical shift difference as $\Delta\delta = \delta_{\mathrm{B}} - \delta_{\mathrm{A}}$, so it is negative if the exchange is happening at the position with a higher chemical shift. Only forward exchange $H_2 \rightarrow HD \rightarrow D_2$ was incorporated in the models for spin-dynamics simulations, and the reverse isotope exchange was neglected assuming the water deuteration factor $f_D = 1$ (measured values: $f_D \approx 96.5\% - 98.4\%$). $k_{ex}$ is a hydrogen site-exchange rate constant. The used NMR parameters are:

$H_2$: (δ = 4.655 ppm, $T_1$ = 5 s, $T_{\mathrm{POC}}$ = 500 s, $J$ = 280 Hz).

P-HH: ($\delta_{\mathrm{A}}$, $\delta_{\mathrm{B}}$, $\Delta\delta = \delta_{\mathrm{B}} - \delta_{\mathrm{A}}$, $T_1$ = 1 s, $J_{\mathrm{HH}}$).

DH: (δ = 4.6025 ppm, $T_1$ = 8 s, $J$ = 43 Hz).

P-DH: ($\delta_A$ (deuterium), $\delta_B$, $T_1$ = 1 s, $\frac{\gamma_D}{\gamma_H}J_{HH}$).

P-HD: ($\delta_A$, $\delta_B$ (deuterium), $T_1$ = 1 s, $\frac{\gamma_D}{\gamma_H}J_{HH}$).

The necessary density matrices and superoperators of this model are given in equations ( 96 ) - ( 99 ), on the next page. For expressing $\widehat{\widehat{K}}$, we made use of the operator of mutual exchange of spin-1 and spin-1/2 in a two-spin system with the first spin being spin-1

$$\widehat{\widehat{\mathrm{Q}}}_{\mathrm{D1\leftrightarrow H2}} = \widehat{\mathrm{Q}}_{\mathrm{D1\leftrightarrow H2}} \otimes \widehat{\mathrm{Q}}_{\mathrm{D1\leftrightarrow H2}}, \text{with } \widehat{\mathrm{Q}}_{\mathrm{D1\leftrightarrow H2}} = \begin{pmatrix} 1 & 0 & 0 & 0 & 0 & 0 \\ 0 & 0 & 0 & 1 & 0 & 0 \\ 0 & 1 & 0 & 0 & 0 & 0 \\ 0 & 0 & 0 & 0 & 1 & 0 \\ 0 & 0 & 1 & 0 & 0 & 0 \\ 0 & 0 & 0 & 0 & 0 & 1 \end{pmatrix} \quad (94)$$

and of the operator of mutual exchange of spin-½ and spin-1 in a two-spin system with the first spin being spin-½

$$\widehat{\widehat{\mathrm{Q}}}_{\mathrm{H1\leftrightarrow D2}} = \widehat{\mathrm{Q}}_{\mathrm{H1\leftrightarrow D2}} \otimes \widehat{\mathrm{Q}}_{\mathrm{H1\leftrightarrow D2}}, \text{with } \widehat{\mathrm{Q}}_{\mathrm{H1\leftrightarrow D2}} = \widehat{\mathrm{Q}}^{\mathrm{T}}_{\mathrm{D1\leftrightarrow H2}}. \quad (95)$$

The forms of the operator of direct product $\widehat{\widehat{Dir}}$, trace $\widehat{\widehat{Tr}}$ can be found in Ref. (*60*).

$$\hat{\rho} = \begin{pmatrix} \hat{\rho}_{\mathrm{HH}} \\ \hat{\rho}_{\mathrm{P-HH}} \\ \hat{\rho}_{\mathrm{DH}} \\ \hat{\rho}_{\mathrm{P-DH}} \\ \hat{\rho}_{\mathrm{P-HD}} \end{pmatrix} \tag{96}$$

$$\hat{\hat{L}} = \begin{pmatrix} \hat{\hat{L}}_{\mathrm{HH}} & 0 & 0 & 0 & 0 \\ 0 & \hat{\hat{L}}_{\mathrm{P-HH}} & 0 & 0 & 0 \\ 0 & 0 & \hat{\hat{L}}_{\mathrm{DH}} & 0 & 0 \\ 0 & 0 & 0 & \hat{\hat{L}}_{\mathrm{P-DH}} & 0 \\ 0 & 0 & 0 & 0 & \hat{\hat{L}}_{\mathrm{P-HD}} \end{pmatrix} \tag{97}$$

$$\begin{aligned}\hat{\hat{K}} \\ = \end{aligned}\begin{pmatrix} -(k'_a + k_{\mathrm{out}})\hat{\hat{1}}_{\mathrm{HH}} & k_{\mathrm{d}}\hat{\hat{1}}_{\mathrm{HH}} & 0 & 0 & 0 \\ \frac{1}{2}k'_{\mathrm{a}}\left(\hat{\hat{1}}_{\mathrm{HH}} + \hat{\hat{Q}}_{\mathrm{H1\leftrightarrow H2}}\right) & -(k_d + k_{HD})\hat{\hat{1}}_{\mathrm{HH}} + k_{\mathrm{ex}}\left(\hat{\hat{Q}}_{\mathrm{H1\leftrightarrow H2}} - \hat{\hat{1}}_{\mathrm{HH}}\right) & 0 & 0 & 0 \\ 0 & 0 & -(k'_{\mathrm{a}} + k_{\mathrm{out}})\hat{\hat{1}}_{\mathrm{DH}} & k_d\hat{\hat{1}}_{\mathrm{DH}} & k_d\hat{\hat{1}}_{\mathrm{DH}} \\ 0 & k_{\mathrm{HD}}\widehat{\widehat{Dir}}_{\mathrm{D}}\widehat{\widehat{Tr}}_1^{\mathrm{P-HH}} & 0.5k'_{\mathrm{a}}\hat{\hat{1}}_{\mathrm{DH}} & -(k_{\mathrm{d}} + k_{\mathrm{ex}})\hat{\hat{1}}_{\mathrm{DH}} & k_{\mathrm{ex}}\hat{\hat{Q}}_{\mathrm{H1\leftrightarrow D2}} \\ 0 & 0 & 0.5k'_{\mathrm{a}}\hat{\hat{Q}}_{\mathrm{D1\leftrightarrow H2}} & k_{\mathrm{ex}}\hat{\hat{Q}}_{\mathrm{D1\leftrightarrow H2}} & -(k_{\mathrm{d}} + k_{\mathrm{HD}} + k_{\mathrm{ex}})\hat{\hat{1}}_{\mathrm{DH}} \end{pmatrix} \tag{98}$$

$$\hat{W} = \begin{pmatrix} k_{\mathrm{in}}\hat{\rho}_{\mathrm{HH^*}} \\ 0 \\ 0 \\ 0 \\ 0 \end{pmatrix} \tag{99}$$

**Model-2b-2HDex.** This is the extension of the Model-1b-1HDex with two bound states where an isotopic exchange happens on both bound states. Therefore, the amount of some system parameters is doubled: $\delta_{\mathrm{A}i}$, $\delta_{\mathrm{B}i}$, $\Delta\delta_i$, $k_{\mathrm{HD}i}$, $k_{\mathrm{ex}i}$, where $i$ =1, 2.

The used NMR parameters are:

$H_2$: (4.655 ppm, $T_1$=5 s, $T_{POC}$=500 s, $J$=280 Hz).

DH: (4.6025 ppm (deuterium), 4.6025 ppm, $T_1$=8 s, $J$=43 Hz).

P-HH: ($\delta_{\mathrm{A}1}$, $\delta_{\mathrm{B}1}$, $\Delta\delta_1 = \delta_{\mathrm{B}1} - \delta_{\mathrm{A}1}$, $T_1$=1 s, $J_1$).

P-H-H: ($\delta_{\mathrm{A}2}$, $\delta_{\mathrm{B}2}$, $\Delta\delta_2 = \delta_{\mathrm{B}2} - \delta_{\mathrm{A}2}$, $T_1$=1 s, $J_2$).

P-DH: ($\delta_{\mathrm{A}1}$ (deuterium), $\delta_{\mathrm{B}1}$, $T_1$=1 s, $\frac{\gamma_D}{\gamma_H}J_1$).

P-D-H: ($\delta_{\mathrm{A}2}$ (deuterium), $\delta_{\mathrm{B}2}$, $T_1$=1 s, $\frac{\gamma_D}{\gamma_H}J_2$).

P-HD: ($\delta_{\mathrm{A}1}$, $\delta_{\mathrm{B}1}$ (deuterium), $T_1$=1 s, $\frac{\gamma_D}{\gamma_H}J_1$).

P-H-D: ($\delta_{\mathrm{A}2}$, $\delta_{\mathrm{B}2}$ (deuterium), $T_1$=1 s, $\frac{\gamma_D}{\gamma_H}J_2$).

The necessary density matrices and superoperators of this model are given in equations ( 100 ) – ( 103 ). All operators used here have been introduced before in the text. Exchange rates are specified in Fig. S17C.

$$\hat{\rho} = \begin{pmatrix} \hat{\rho}_{\mathrm{HH}} \\ \hat{\rho}_{\mathrm{P-HH}} \\ \hat{\rho}_{\mathrm{P-H-H}} \\ \hat{\rho}_{\mathrm{DH}} \\ \hat{\rho}_{\mathrm{P-DH}} \\ \hat{\rho}_{\mathrm{P-D-H}} \\ \hat{\rho}_{\mathrm{P-HD}} \\ \hat{\rho}_{\mathrm{P-H-D}} \end{pmatrix} \qquad (100)$$

$$
\hat{\hat{L}} = \begin{pmatrix}
\hat{\hat{L}}_{\mathrm{HH}} & 0 & 0 & 0 & 0 & 0 & 0 & 0 \\
0 & \hat{\hat{L}}_{\mathrm{P-HH}} & 0 & 0 & 0 & 0 & 0 & 0 \\
0 & 0 & \hat{\hat{L}}_{\mathrm{P-H-H}} & 0 & 0 & 0 & 0 & 0 \\
0 & 0 & 0 & \hat{\hat{L}}_{\mathrm{DH}} & 0 & 0 & 0 & 0 \\
0 & 0 & 0 & 0 & \hat{\hat{L}}_{\mathrm{P-DH}} & 0 & 0 & 0 \\
0 & 0 & 0 & 0 & 0 & \hat{\hat{L}}_{\mathrm{P-D-H}} & 0 & 0 \\
0 & 0 & 0 & 0 & 0 & 0 & \hat{\hat{L}}_{\mathrm{P-HD}} & 0 \\
0 & 0 & 0 & 0 & 0 & 0 & 0 & \hat{\hat{L}}_{\mathrm{P-H-D}}
\end{pmatrix} \tag{101}
$$

$$
\hat{\hat{\boldsymbol{K}}} = \begin{pmatrix}
-(\boldsymbol{k}'_{\mathrm{a1}} + \boldsymbol{k}_{out})\mathbf{1}_{HH} & \boldsymbol{k}_{\mathrm{d1}}\hat{\hat{\mathbf{1}}}_{\mathrm{HH}} & \mathbf{0} & \mathbf{0} & \mathbf{0} & \mathbf{0} & \mathbf{0} & \mathbf{0} \\
\frac{\mathbf{1}}{\mathbf{2}}\boldsymbol{k}_{a1}\left(\hat{\hat{\mathbf{1}}}_{\mathrm{HH}} + \hat{\hat{\boldsymbol{Q}}}_{\mathrm{H1\leftrightarrow H2}}\right) & -(\boldsymbol{k}_{\mathrm{d1}} + \boldsymbol{k}_{\mathrm{a2}} + \boldsymbol{k}_{\mathrm{HD1}})\hat{\hat{\mathbf{1}}}_{\mathrm{HH}} + \boldsymbol{k}_{\mathrm{ex1}}\left(\hat{\hat{\boldsymbol{Q}}}_{\mathrm{H1\leftrightarrow H2}} - \hat{\hat{\mathbf{1}}}_{\mathrm{HH}}\right) & \boldsymbol{k}_{\mathrm{d2}}\hat{\hat{\mathbf{1}}}_{\mathrm{HH}} & \mathbf{0} & \mathbf{0} & \mathbf{0} & \mathbf{0} & \mathbf{0} \\
\mathbf{0} & \boldsymbol{k}_{\mathrm{a2}}\hat{\hat{\mathbf{1}}}_{\mathrm{HH}} & -(\boldsymbol{k}_{\mathrm{d2}} + \boldsymbol{k}_{\mathrm{HD2}})\hat{\hat{\mathbf{1}}}_{\mathrm{HH}} + \boldsymbol{k}_{\mathrm{ex2}}\left(\hat{\hat{\boldsymbol{Q}}}_{\mathrm{H1\leftrightarrow H2}} - \hat{\hat{\mathbf{1}}}_{\mathrm{HH}}\right) & \mathbf{0} & \mathbf{0} & \mathbf{0} & \mathbf{0} & \mathbf{0} \\
\mathbf{0} & \mathbf{0} & \mathbf{0} & -(\boldsymbol{k}'_{\mathrm{a1}} + \boldsymbol{k}_{\mathrm{out}})\hat{\hat{\mathbf{1}}}_{\mathrm{DH}} & \boldsymbol{k}_{d1}\hat{\hat{\mathbf{1}}}_{\mathrm{DH}} & \mathbf{0} & \boldsymbol{k}_{\mathrm{d1}}\hat{\hat{\mathbf{1}}}_{\mathrm{DH}} & \mathbf{0} \\
\mathbf{0} & \boldsymbol{k}_{\mathrm{HD1}}\widehat{\widehat{\boldsymbol{Dir}}}_{\mathrm{D}}\widehat{\widehat{\boldsymbol{Tr}}}_{\mathbf{1}}^{\mathrm{P-HH}} & \mathbf{0} & \mathbf{0.5}\boldsymbol{k}'_{\mathrm{a1}}\hat{\hat{\mathbf{1}}}_{\mathrm{DH}} & -(\boldsymbol{k}_{\mathrm{d1}} + \boldsymbol{k}_{\mathrm{a2}} + \boldsymbol{k}_{\mathrm{ex1}})\hat{\hat{\mathbf{1}}}_{\mathrm{DH}} & \boldsymbol{k}_{\mathrm{d2}}\hat{\hat{\mathbf{1}}}_{\mathrm{DH}} & \boldsymbol{k}_{\mathrm{ex1}}\hat{\hat{\boldsymbol{Q}}}_{\mathrm{H1\leftrightarrow D2}} & \mathbf{0} \\
\mathbf{0} & \mathbf{0} & \boldsymbol{k}_{\mathrm{HD2}}\widehat{\widehat{\boldsymbol{Dir}}}_{\mathrm{D}}\widehat{\widehat{\boldsymbol{Tr}}}_{\mathbf{1}}^{\mathrm{P-H-H}} & \mathbf{0} & \boldsymbol{k}_{\mathrm{a2}}\hat{\hat{\mathbf{1}}}_{\mathrm{DH}} & -(\boldsymbol{k}_{\mathrm{d2}} + \boldsymbol{k}_{\mathrm{ex2}})\hat{\hat{\mathbf{1}}}_{\mathrm{DH}} & \mathbf{0} & \boldsymbol{k}_{\mathrm{ex2}}\hat{\hat{\boldsymbol{Q}}}_{\mathrm{H1\leftrightarrow D2}} \\
\mathbf{0} & \mathbf{0} & \mathbf{0} & \mathbf{0.5}\boldsymbol{k}'_{\mathrm{a1}}\hat{\hat{\mathbf{1}}}_{\mathrm{DH}} & \boldsymbol{k}_{\mathrm{ex1}}\hat{\hat{\boldsymbol{Q}}}_{\mathrm{D1\leftrightarrow H2}} & \mathbf{0} & -(\boldsymbol{k}_{\mathrm{d1}} + \boldsymbol{k}_{\mathrm{a2}} + \boldsymbol{k}_{\mathrm{HD1}} + \boldsymbol{k}_{\mathrm{ex1}})\hat{\hat{\mathbf{1}}}_{\mathrm{DH}} & \boldsymbol{k}_{\mathrm{d2}}\hat{\hat{\mathbf{1}}}_{\mathrm{DH}} \\
\mathbf{0} & \mathbf{0} & \mathbf{0} & \mathbf{0} & \mathbf{0} & \boldsymbol{k}_{\mathrm{ex2}}\hat{\hat{\boldsymbol{Q}}}_{\mathrm{D1\leftrightarrow H2}} & \boldsymbol{k}_{\mathrm{a2}}\hat{\hat{\mathbf{1}}}_{\mathrm{DH}} & -(\boldsymbol{k}_{\mathrm{d2}} + \boldsymbol{k}_{\mathrm{HD2}} + \boldsymbol{k}_{\mathrm{ex2}})\hat{\hat{\mathbf{1}}}_{\mathrm{DH}}
\end{pmatrix} \tag{102}
$$

$$
\widehat{W} = \begin{pmatrix}
k_{in}\hat{\rho}_{\mathrm{HH}^*} \\ 0 \\ 0 \\ 0 \\ 0 \\ 0 \\ 0 \\ 0
\end{pmatrix} \tag{103}
$$

2.7 Field-dependent modeling of the HD-PHIP and the PNL

To determine the parameter ranges compatible with the experimentally observed field-dependence of the HD-PHIP and the PNL shown in Fig. 2C of the main article, we first modelled both effects separately, before attempting the joint modeling of both effects.

The experimental spectra of reference were picked from the spectra presented in Fig. S8, which shows the overlay of three $^1$H-PHIP spectra and three reference spectra with $n$-$H_2$ bubbling at each of the three static fields used for the experiments. At each field, the $^1$H-PHIP spectrum with the best shimming was picked for comparison with simulated spectra and for presentation in Fig. 2C of the main article. The average signal integrals for $H_2$ and HD observed in the spectra shown in Fig. S8 are summarized in Table S2.

For both the HD-PHIP and the PNL signal, the available parameter space was first searched by manual comparison of the experimental and the simulated spectra, and in the following, the parameter space spanned by $\Delta\delta$ and $J_{HH1}$ was subject to a grid search, with optimization of kinetic parameters at each grid point to obtain the minimum RMSD in the signal region between experimental and simulated spectra.

All modeling parameters other than $\Delta\delta$, $J_{HH1}$ and selected kinetic parameters were kept constant during all simulations (see Table S5). Simulations were performed for the data at 309 K. NMR experiment parameters for the simulations were taken from the experimental datasets, with two exceptions: 1) The line broadening applied during apodization with a monoexponentially decaying function was increased to reproduce the experimentally observed linewidths (experiments: 0.3 Hz, simulations: 0.7 Hz). 2) The simulated spectral width was reduced 16x with respect to the experiments, to reduce computational cost. No aliasing occurred due to the reduced spectral width.

A short summary of the compatible parameter ranges obtained from the systematic scanning is summarized in Fig. S25. For the model in Fig. S17B, we define the lifetime of the bound state, listed as $\tau_{PNL}$ or $\tau_{HD-PHIP}$, as $\tau = (k_d + k_{HD} + k_{ex})^{-1}$. Details of the systematic parameter scanning are described in the following two sections (2.8 & 2.9). Joint simulations of PNL and HD-PHIP are described in section 2.10.

### 2.8 Simulation of the PNL lineshape

For modeling of the PNL, it was attempted to reproduce the three spectra shown in Fig. 2C of the main article. The experimental data was collected at three different $B_0$-fields (7.05 T, 14.10 T & 21.15 T) using samples with 1 μM *j*Hmd and 3 μM $^{13}CH_2$=$H_4$MPT. Samples were prepared in $D_2O$-buffer (95.4 – 98.4% deuteration) with *p*D 6.0, 1 mM EDTA and 120 mM potassium phosphate. The enrichment of $p$-$H_2$ was 87%.

Under the experimental conditions, the PNL behaves as follows:

- The PNL signal monotonically increases with increasing field (7 T to 21 T)
- At all three fields, the PNL signal shape fits an overlay of two fully absorptive lorentizians with opposing phase, with negative part at high δ and positive part at low δ. Dispersive parts of the PNL line have small amplitudes in the real part of the spectrum.
- The PNL signal position does not notably shift away from the position of free $H_2$.

It was first attempted to reproduce these qualitative features in numerical simulations. For simplicity, simulations of the PNL line shape were performed in the following using the model in Fig. S17A (only one bound intermediate, no isotope exchange).

From the increase of the PNL with field, it was concluded, that dissociation from the bound state must be faster than one chemical shift evolution period, even at the highest field tested. Indeed, by testing different ratios of simulation parameters, it was found that

$$(5 * \tau_{PNL})^{-1} > |\Delta\nu(21.1T)| > |\Delta\nu(7.0T)| \gg |J_{HH}|, \quad (104)$$

with $\tau_{PNL} = (k_d + k_{ex})^{-1}$, was required to qualitatively reproduce the observed increase of the PNL with field. The factor $5^{-1}$ ensures, that the intensity ratios at the three fields roughly match the observation. In this limit, the phase of the PNL signal is fixed by the sign of $J_{HH}$, and

$$J_{HH} > 0 \quad (105)$$

was found to be required to obtain the observed signal sign. Exemplary plots showing the general trends of the PNL on $J_{HH}$ and bound state lifetime $\tau_{PNL} = (k_d + k_{ex})^{-1}$ are shown in Fig. S19.

As will be shown in section 2.7, the HD-PHIP can be reproduced using the kinetic parameters $k_a$, $k_d$ and $k_{HD}$ obtained from the kinetics analysis (Table S4 & section 2.2). This was not the case for the PNL. No combination of reasonable $J$ and $\Delta\delta$ was found, which provided a good fit to the PNL-line using the kinetic parameters stated in Table S4, when we turned to the model in Fig. S17B. To reproduce the PNL signals observed, much faster association and dissociation rates were required ($k_d \approx 10^4 - 10^6\ s^{-1}, k_a \approx 10^4 - 10^6\ mM^{-1}\ s^{-1}$), than those obtained from the fitting of the net isotope exchange kinetics ($k_d \approx (35 \pm 2)\ s^{-1}, k_a \approx (40 \pm 16)\ mM^{-1}\ s^{-1}$). Two exemplary simulations are compared in Fig. S20. For fast association and dissociation, the kinetics need to be rate limited by $k_{HD}$ to fulfill the net $[H_2]$ consumption kinetics (see equation ( 10 )). With $k_d \gg k_{HD}$, however, it is unavoidable, that the ratio $[HD]_{max}/[H_2]_0$ is comes close to 0.5 (see Fig. S13 and equation ( 23 )), whereas we experimentally observe $[HD]_{max}/[H_2]_0 \approx 0.1$. In Fig. S20D this results in a strong overestimation of the HD concentration. Therefore we concluded, that it is impossible to simultaneously reproduce the PNL data and the net isotope exchange kinetics within the model assuming only one bound-state geometry (Fig. S17B). From this finding we hypothised, that the bound state causing the PNL is not the bound state in which hydrogen isotope exchange predominantly occurs.

The PNL peaks at lower temperature than the HD-PHIP (Fig. S11), which could imply that the PNL is created in an earlier catalytic intermediate than the HD-PHIP. We thus assumed a mechanism with two bound state geometries (Fig. S13C), with the PNL being created in the early intermediate P-HH, and the isotope exchange happening in the late intermediate P-H-H. In this model it was possible to simultaneously reproduce the PNL and the isotope exchange kinetics, if a fast pre-equilibrium was assumed ($k_{d1} \gg k_{a2}$) and using $k_{HD1} = 0$ (isotope exchange in the late intermediate).

To systematically search for the parameter ranges compatible with the PNL effects observed, we returned to the model with only one bound intermediate and without no isotope exchange (Fig. S17A), since if a fast pre-equilibrium is assumed ($k_{d1} \gg k_{a2}$), the late intermediate (P-H-H) will not have significant impact on the PNL observed. The much simpler model in Fig. S17A was thus used.
The parameter space spanned by $J$ and $\Delta\delta$ was systematically scanned and at each point $k_d$ was optimized to yield the best possible fit between the experimental spectra and the spectra simulated with the model in Fig. S17A. Our simulations can only provide an upper bound limit for the possible PHIP intensities via a given mechanism, since many possible signal loss mechanisms remain uncharacterized. Therefore during fitting, the spectra were simulated for a given choice of $J$, $\Delta\delta$ using an upper bound value for $k_a'$ and the best fit value for $k_d$, and the spectrum was uniformly downscaled by $f_{downscaling} = [0; 1]$ to the intensity that yielded the minimum RMSD between the experimental spectra and the downscaled simulated spectra.

Access to the upper bound value for $k_a'$, which determines the maximum possible PHIP intensity for a given $k_d$, was obtained from the steady state analysis of the model with two bound state intermediate geometries (Fig. S13B) presented in section 2.3.2. From equation ( 29 )

$$\text{(for model Fig. S13B):} \quad K_m = \frac{k_{d1}}{k_{a1}} \frac{k_{d2}}{k_{d2} + k_{a2}}, \tag{29}$$

We obtain

$$\text{(for model Fig. S13B):} \quad k_{a1} \leq \frac{k_{d1}}{K_m}. \tag{106}$$

With this constraint, and remembering our assumption that PNL is formed in the early intermediate P-HH, we estimate $k_a' = k_a[P]$ via

$$k_a' \leq \frac{k_d}{K_m}[P] = [P]_0 \frac{k_d}{K_m + [H_2]_0} \approx \frac{k_d}{(6.4 \pm 0.4) * 10^3}, \tag{107}$$

using $[H_2]_0 \approx 5.5 mM$, $[P]_0 = 1 \mu M$ and $K_m \approx (0.9 \pm 0.4)\ mM$. We thus fixed the upper bound estimate $k_a' = k_d/(6.0 * 10^3)$. Multiple parameter scans were performed for different choices of $\delta_{av}$ and $k_{ex}$. Exemparily, the parameter scan for $\delta_{av} = 4.3\ ppm$ and $k_{ex} = 0$, which yields the best fit result, is given in Fig. S21.

The best fits in all cases were obtaind for $J = +300\ Hz$, which is the upper bound limit of the range of $J$ values we scanned. Good fits were obtained only for $J \geq 50Hz$ and $|\Delta\delta| \geq 0.5ppm$ in the bound state. The average chemical shift in the bound state seems to be around 3.5 $ppm\ <$ $\delta_{av} < 5.0\ ppm$ (best fit: $\delta_{av} = 4.3\ ppm$, see Fig. S22). Lifetimes of the bound state causing the PNL in the range of $\tau_{PNL} = (k_d + k_{ex})^{-1} = 1\mu s - 100\mu s$ can reproduce the data, and good fits were only obtained for $k_{ex} < 10\ k_d$. The values for $J$ and $\Delta\delta$ used in the simulations should reflect the time averaged NMR parameters of an ensemble of intermediates exchanging faster than $\tau_{PNL}$.

The large positive *J*-coupling ($J \geq 50Hz$) required implies, that a $^{1}J_{HH}$ coupling is present in the bound state ensemble causing the PNL. A non-classical hydride with Fe-$\eta^2$-$H_2$ type binding appears the only reasonable choice.

### 2.9 Simulation of the HD-PHIP in two different modeling scenarios

In section 2.5 it is shown that an analytical model based on a simplified kinetic scheme and a single bound state is able to describe the field dependence of the HD-PHIP in the range of 10 mT – 7 T. This strongly supports a coherent *J*-coupling based mechanism.

Since the PNL effect observed, however, requires a kinetic model with at least two distinct bound states (Fig. S17C) as discussed in section 2.6, it is a priori not known, which of the two bound state ensembles contains the *J*-coupling causing the HD-PHIP. The analysis of the PNL (section 2.6) strongly suggests, that the isotope exchange is happening in ensemble 2 (as assumed in Fig. S17C). We therefore investigated two scenarios in numerical simulations:

Scenario A): The *J*-coupling causing the HD-PHIP is in ensemble 1, whereas isotope exchange happens in ensemble 2.

Scenario B): The *J*-coupling causing the HD-PHIP is in ensemble 2, where isotope exchange also is happening.

For both scenarios, it was again estimated which bound state parameters are compatible with the three spectra shown in Fig. 2C of the main article, and with the isotope exchange kinetics measured.

In scenario B), only ensemble 2 is involved in the creation of the HD-PHIP effect. We thus neglected ensemble 1 during modeling of this scenario, and performed numerical simulations in the framework of a model with only one bound state (Fig. S17B), were the bound state was assumed to represent ensemble 2. As discussed in section 2.3.2, in the limit $k_{d1} \gg k_{a2}$ a simplified treatment of the isotope exchange kinetics is possible, only considering the second bound state (P-H-H).

For this bound state, the kinetic parameters were thus approximated according to equations equations ( 39 ) – ( 42 ) from the parameters extracted from the isotope exchange kinetics. We fixed $k_d = 35\ s^{-1}$, $k_{HD} = 257\ s^{-1}$ and we optimized $k_{ex}$ for every point in the $\Delta\delta - J$ plane to yield the smallest RMSD between the experimental and simulated HD signals.

For $k_a'$, we chose the value that best reproduces the isotope exchange kinetics for the samples used ($k_a' = 8.0 * 10^{-3}\ s^{-1}$; $k_a = 48\ mM^{-1}\ s^{-1}$), whereas from the values in Table S4, one would estimate $k_a' \approx k_d[P]_0/(K_m + c_0) \approx (6.5 \pm 1.6) * 10^{-3}\ s^{-1}$ (assuming $\Delta[P]_0 \approx 0.2\ \mu M$ and $\Delta c_0 \approx 0.5\ mM$). For fitting, a lower bound for $k_{ex}$ of 70 $s^{-1}$ was used. To avoid problems with signal overlap with water, only the middle and the low-$\delta$ multiplet peak of HD were used to compute the RMSD.

In the model with one bound state geometry only (Fig. S17B), sign rules link the sign of $J_{HH}$ to the sign of $\Delta\delta$ (see section S2.4). In this model, the positive HD-PHIP intensity observed thus requires $\Delta\delta * J_{HH} > 0$, i.e. that for positive $J_{HH}$, isotope exchange must be happening at the low chemical shift position ($\Delta\delta > 0$), whereas for negative $J_{HH}$ it must be happening at the high chemical shift position ($\Delta\delta < 0$). In simulations, both scenarios yield identical simulated spectra, so only the quadrant $\Delta\delta > 0$ and $J_{HH} > 0$ was scanned, while the and results are equally valid for the quadrant $\Delta\delta < 0$ and $J_{HH} < 0$. Results are independent of the average chemical shift $\delta_{av1} = (\delta_{A1} + \delta_{B1})/2$ during simulation.

For all $|J| > 3\ Hz$, fits of essentially equal quality to the best fit data can be found. All these best fit solutions are in a fairly narrow range for the chemical shift difference $|\Delta\delta|$ of $0.1\ ppm \leq |\Delta\delta| \leq 0.3\ ppm$. These results unfortunately provide very little structural insight into the intermediate causing the HD-PHIP, since many plausible intermedeates feature $|J| > 3\ Hz$ for the computed *J*-couplings. In the computed structural models (Tables S8 – S11), in most cases bigger chemical shift differences were obtained from computations. The small value for $|\Delta\delta|$ estimated from the data may result from chemical shifts that are similar by chance, or it may be a hint, that more sophisticated underlying enzyme kinetics are relevant for the HD-PHIP creation.

Scenario A) is more sophisticated to model, since it requires the model with two distinct bound state geometries (Fig. S17C), for which the relevant kinetic rate constants are underdetermined by the isotope exchange kinetics measured.

The parameter space relevant to be searched can however be notably restricted by the isotope exchange kinetics, if the following assumptions are made:

1. We assumed a fast preequilibrium between free hydrogen and the bound state ensemble 1 ($k_{d1} \gg k_{a2}$), as implied by our results obtained from the PNL analysis (section 2.6).
2. We assumed no isotope exchange in ensemble 1 ($k_{HD1} = 0$), as also implied by our results obtained from the PNL analysis (section 2.6).
3. We further assumed no position exchange in ensemble 1 ($k_{ex1} = 0$), which requires some explanation:
    a. In the limit $k_{d1} \gg k_{a2}$, $k_{ex1}$ is not constrained by the observed isotope exchange kinetics.
    b. We know that $k_{ex2}$ has to be non-zero in the assumed limit $k_{d1} \gg k_{a2}$: A lower level constraint for $k_{ex2}$ can be obtained from the ratio $[HD]_{max}/[H_2]_0$ in this limit. The observed ratio $[HD]_{max}/[H_2]_0 \approx 0.1$ requires that $k_{d2}/k_{ex2} \leq 0.4$ under the assumption $k_{d1} \gg k_{a2}$.
    c. In the limit $k_{d1} \gg k_{a2}$, $k_{ex1} > 0$ only leads to a downscaling of the observed PHIP effects. The desired upper limit estimate for the possible PHIP intensities are obtained for $k_{ex1} = 0$.

With assumptions 1 – 3, the isotope exchange kinetics obtained from the steady state analysis of the model with two bound state geometries (Fig. S17C) reduces to the steady state soultions obtained from the model with one bound state geometry (Fig. S17B), with an effective association constant

$$k_a|_{k_{d1} \gg k_{a2},\ k_{ex1}=0} \approx \frac{k_{a1}k_{a2}}{k_{d1}}. \quad (39)$$

Thus, the kinetic rates extracted from isotope exchange kinetics using a model with one bound state geometry (Table S4) characterize ensemble 2 well. We used the best-fit values to fix $k_{d2} = k_d = 35\ s^{-1}$, $k_{HD2} = k_{HD} = 257\ s^{-1}$ and $k_{ex2} = k_{ex} = 300\ s^{-1}$, according to equations ( 40 ) - ( 42 ).

$$k_{d2}|_{k_{d1} \gg k_{a2}} \approx k_d, \quad (40)$$

$$k_{HD2}|_{k_{d1} \gg k_{a2}} \approx k_{HD} \quad (42)$$

and

$$k_{ex2}|_{k_{d1} \gg k_{a2}} \approx k_{ex}. \quad (42)$$

Then, for different fixed guess-values of $k_{a2}/k_{d1}$, we scanned the $\Delta\delta_1 - J_{HH1}$ parameter space, determining the value of $k_{d1}$ that provides the minimum RMSD between measured and simulated spectra at each point in the $\Delta\delta_1 - J_{HH1}$ plane. Again downscaling of the spectra by an arbitrary factor $f_{downscaling} = [0; 1]$ was used to account for the fact, that simulations could always be overestimating the PHIP intensity, due to uncharacterized loss processes. Guess values of $k_{a2}/k_{d1} = 10^{-1}, 10^{-2}, 10^{-3}, 10^{-4}, 10^{-5}$ and $10^{-6}$ were sampled in different scans. With $k_{d1}$ and $k_{a2}/k_{d1}$ given, we can estimate $k_{a1}$ from equation ( 39 ), fixing the last kinetic parameter missing.

As illustrated in Fig. S24, a range of parameters between ($\Delta\delta_1 = -0.03\ ppm$; $J_{HH1} = 10\ Hz$) and ($\Delta\delta_1 = -1.0\ ppm$; $J_{HH1} = 300\ Hz$) is capable of reproducing the experimental HD-PHIP data. If scenario A applies, The ratio $k_{a2}/k_{d1}$ should be bigger than $10^{-3}$, since for smaller ratios, the HD-PHIP data cannot be reproduced.

2.10 Joint simulation of PNL and HD-PHIP in models with two bound state geometries.

Based on the modeling restraints obtained from analysis of the HD-PHIP and the PNL, briefly summarized in Fig. S25, we attempted to find models, which simultaneously reproduce the HD-PHIP and the PNL observed. Since we found, that the spin dynamics model with one bound state geometry (Fig. S17B) is not able to reproduce the PNL and the isotope exchange kinetics simultaneously, we approached the joint simulation of the PNL, the HD-PHIP and the isotope exchange kinetics assuming the model with two different bound state geometries (Fig. S17C).

For this model, our preceeding analysis (section 2.6) suggests, that the PNL must result form fast exchange between free $H_2$ and the bound form of ensemble 1 (P-HH), with the need for a large positive $J_{HH1}$ in ensemble 1 (see section 2.6). The HD-PHIP, however could be created either in ensemble 1 (scenario A) or in ensemble 2 (scenario B), and the *J*-couplings causing the effect could be as small as $|J| > 3\ Hz$ (see section 2.7).

Thus three conceivable scenarios emerge, which are all equally capable of reproducing the $^1$H-spectra together with the observed isotope exchange kinetics:

A) Creation of both PNL and HD-PHIP within ensemble 1 by a large positive $J_{HH1}$ and isotope exchange in ensemble 2 in a completely dissociated ensemble ($J_{HH2} = 0$). This model requires a switch of sign for $\Delta\delta$ between the two ensembles.

B1) Creation of the PNL within ensemble 1 by a large positive $J_{HH1}$ and creation of the HD-PHIP in ensemble 2 via a small positive $J_{HH2}$.

B2) Creation of the PNL within ensemble 1 by a large positive $J_{HH1}$ and creation of the HD-PHIP in ensemble 2 via a small negative $J_{HH2}$.

Exemplary simulation parameters for these scenarios are listed in Table S6, and simulation results are shown in Fig. S27 and Fig. 2C of the main article, illustrating that agreement with the data is good in all three cases. Simulation parameters were manually adjusted to achieve good reproduction of the experimentally observed $^1$H spectra. Least squares fitting of the data was not performed.

Due to the large number of free modeling parameters, it was practically impossible to achieve a full sampling of the available parameter space for the model with chosen two different bound state geometries (Fig. S17C), so we cannot be sure that we sampled all the relevant parameter space, yet all parameter sets we found that were able to reproduce the PHIP spectra and the isotope exchange kinetics simultaneously share these common features:

1. A fast equilibrium between free $H_2$ and ensemble 1,
2. A large positive $J_{HH1}$ within ensemble 1, characteristic for an Fe-$\eta^2$-$H_2$ species,
3. Hydrogen isotope exchange happening in ensemble 2,
4. Rate limitation by the transition between ensembles 1 and 2.

Whereas for ensemble 1, a clear structural assignment can be made, due to the large positive $J_{HH1}$ required, the structural assignment for ensemble 2 is ambiguous: Ensemple 2 could be an intermediate, where the two atoms originating from $H_2$ are fully dissociated $J_{HH2} = 0$, or it could be an intermediate, where they still experience a sizable time-averaged *J*-coupling.

2.11 Simulation of PHIP-CEST curves

PHIP-CEST data was acquired at different static fields (7 T, 14 T & 21 T) and varying spin-lock field ($\gamma B_1$ = 42 Hz - 3000 Hz) as described in section 1.11. The $H_2$-PHIP-CEST curves obtained are shown in Fig. 4B of the main article, Fig. S29 and Fig. S30, whereas the HD-PHIP-CEST curves obtained are shown in Fig. 4A of the main article and Fig. S31. As apparent, both the $H_2$-PHIP-CEST effect and the HD-PHIP-CEST effect saturate at $\gamma B_1 \approx$ 1000 Hz, and they vanish as the spin-lock field range dropps below $\gamma B_1$ = 333 Hz. Thus we estimate the lifetime of the associated bound state(s) to be around $\tau_{PHIP-CEST} \approx 1 - 2$ ms, as stated in the main article.

Simulations of PHIP-CEST curves were performed, in an attempt to reproduce the data shown in Fig. 4 A & B of the main article. Simulations were performed using the model that assumes one bound state geometry (Fig. S17B) and irreversible hydrogen isotope exchange. Starting from $\hat{\rho} = 0$, the density operator was propagated through the bubbling period without application of an external spin lock pulse, and the resulting density operator was then subject to free evolution under spin lock irradiation with different offsets. For each offset, the resulting density operator was propagated through the spin-locking periods and at the end of the spin lock period, the expectation values for the $H_2$ and HD $^1$H-polarizations ($\langle \hat{I}_{z,1} \rangle + \langle \hat{I}_{z,2} \rangle$ for $H_2$ and $\langle \hat{I}_{z,1} \rangle$ for HD) was extracted and plotted against the spin lock offset, to create the simulated PHIP-CEST profiles. A perfectly homogeneous $B_1$ field was assumed for field locking.

For the model with one bound state geometry Fig. S17B, the measured PHIP-CEST profiles can be qualitatively reproduced with parameters that reproduce the overall isotope exchange kinetics, e.g. using the parameters listed in Table S7. From these simulations, chemical shifts for the bound species were extracted as (10.5 ± 0.5) ppm and (4 ± 2 ) ppm, by visually judging the agreement between experimental and simulated data. Confidence intervals were estimated by varying settings for one of the chemical shifts, while keeping all other parameters constant, at the values listed in Table S7. With these chemical shift values, a tentative assignment to structure **5** in Fig. 3A of the main article can be made.

To reproduce the HD-PHIP-CEST profile, the site of preferential hydrogen isotope exchange has to be placed at the high chemical shift position (10.5 ppm) and a negative *J*-coupling in between the two hydrogen atoms is required in the bound state, if the model shown in Fig. S17B is assumed. It should be mentioned, though, that no modeling parameters were found, which simultaneously reproduces the $H_2$-PHIP-CEST and the HD-PHIP-CEST effects observed in quantitative fashion. This indicates, that model with one bound state geometry (Fig. S17B) we chose for modeling is too simplistic and that more complex kinetic models need to be assumed, to properly reproduce the PHIP-CEST data.

### 2.12 Structural Modeling

Models of possible reaction intermediates were constructed starting from two different source structures: Structural models denoted **E** & **G** were constructed based on the high-resolution closed-conformation crystal structure (PDB: 6hav) published in ref. (*10*). Structural models denoted **A** & **C** were constructed starting from the QM/MM models published in figures 9 & 11 of ref. (*16*), which were kindly provided by Arndt R. Finkelmann and Markus Reiher.

We distinguish the model structures by the number and position of hydrogen atoms, considered relevant to the hydrogen splitting reaction. For the hydrogens stemming from *p*-$H_2$, it was assumed, that they can occupy the following bound state positions:

- The free coordination site at the iron, as a "non-classical hydride" with side-on binding, **Fe-$\eta^2$-$H_2$**.
- The free coordination site at the iron, as a "classical hydride", **Fe-$\sigma$-H**.
- The cysteine 176 sulfur atom, **SH (Cys176)**, pointing either towards or away from a nearby oxo group of the pterin ring.
- The 2-pyridinol oxygen atom of FeGP, **OH (pyridinol)**, pointing either towards or away from the active site.
- The *pro*-R position of the $C_{14}$ methylene group of $CH_2$=$H_4$MPT, **$C_{14}$-H (*pro*-R)**.

The models were categorized according to the number of these sites occupied by hydrogen atoms into "2H" models (in which two of these sites are occupied) and "3H" models (in which three of these sites are occupied).

An overview over the "2H" models computed, numbered **A1**-**A9** and **E1**-**E9**, is provided in Table S8. Likewise, an overview over the "3H" models computed, numbered **C1**-**C10** and **G1**-**G8**, is provided in Table S10.

All models were (re)optimized at the TPSS-D3BJ/def2-TZVP level, as described in section S1.14. Some models were not successfully optimized, but rather converged to one of the other structures, as indicated in the tables.

### 2.13 NMR parameter calculations

The computed NMR shifts and couplings are given in Table S9 for the "2H" models (**A1**-**A9** and **E1**-**E9**) and in Table S11 for the "3H" models (**C1**-**C10** and **G1**-**G8**). Computed chemical shift and *J*-coupling ranges are further graphically summarized in Figs. 3A and 4A of the main article.

Based on our benchmark calculations, discussed in section 2.14, we estimate that for a given structural model the computed $^1$H chemical shifts are accurate to about 1 ppm and *J*-couplings should be accurate to about 1-2 Hz. Deviations between analogous models (A6/7/8 vs E6/7/8 and C3/4/7/8 vs G3/4/7/8, respectively) are, however, much larger than the uncertainties estimated from the benchmarks (up to 3.6 ppm for chemical shifts and up to 9 Hz for *J*-couplings, for these models). This indicates that the biggest uncertainty for the computed NMR parameters lies in the uncertainty in the actual model geometries.

The NMR parameters compatible with the observed PNL effects fits well the computed NMR parameters of the Fe-$\eta^2$-$H_2$ intermediates (**A1**, **C1** & **G2**). For **C2**, the chemical shift difference obtained is too small to be compatible with the PNL, yet for ensemble 1 it can be expected that in addition to **C1**, **C2** & **G2** also the dissociated intermediates **C3-10** and **G3-8** contribute with lower population, which would increase the time-averaged chemical shift difference.

As discussed in section 2.8, the field dependent $^{1}$H-PHIP experiments data poorly restrains the NMR-parameters in ensemble 2. The *J*-coupling could be positive, negative or even zero for this ensemble. As illustrated in Table S6, ensemble 2 is thus compatible with diverse modeling scenarios and a clear assignment to a structural motive is not possible.

The chemical shift positions observed via PHIP-CEST (10.5 ppm (±0.5 ppm) and 4 ppm (±2 ppm)) are well covered by the chemical shift ranges computed, yet, no unique assignment is possible based on the chemical shifts. The position of $H^+ \rightleftarrows D^+$ exchange at 10.5 ppm is compatible with both the OH (pyridinol) and the SH (Cys176) computed chemical shifts. The 4 ppm (±2 ppm) chemical shift most likely corresponds to that s computed for the $C_{14}$-H (*pro*-R) position, although this assignment is not unique.

### 2.14 NMR parameter calculations: accuracy benchmark

We performed several calculations using different density functionals, basis sets, and treatments of the environment on a few arbitrarily chosen models, in order to gauge the uncertainty of the calculated properties. The effect of different functionals (TPSS (*47*) vs r$^2$SCAN (*64*)) and basis sets (pcSseg-*n* (*53*), denoted as "pS*n*") on the computed chemical shifts is shown in Table S12. Based on these results we estimate the basis set incompleteness errors to be below 0.3 ppm, while errors due to the choice of functional are of similar magnitude. The effect of the point charge embedding is shown in Table S13 by comparing to a calculation, in which the QM region is embedded in a CPCM implicit solvent model with a dielectric constant $\varepsilon$=4.0 (a value commonly used to model proteins). The deviations of up to 0.4 ppm signify that the MM point charge embedding is important, however, minor changes in the positions and magnitudes of the charges are unlikely to result in larger errors than this. Overall, errors due to the choice of functional, basis set, and particular embedding scheme can be expected to add up to as much as 0.5 ppm but should be lower than 1 ppm. Finally, a systematic deviation of the chemical shifts with respect to the experimental values can be expected due to the particular choice of reference – as the TMS system is treated somewhat differently, we cannot expect perfect error compensation. This deviation is difficult to estimate but is unlikely to be more than a few tenths of a ppm.

A similar analysis was performed for the indirect nuclear spin-spin coupling constants. A comparison using the pJ1-pJ3 basis sets (*56*) and the PBE (*65*) and PBE0 (*55*) functionals is shown in Table S14. The basis set incompleteness errors in this example are below 0.2 Hz, except for the large $^{1}J_{HH}$ coupling in the $H_2$ molecule. For another model, **C3**(OHtoH-FeH-SHtoO), the differences between pJ3 and pJ2 were between -0.08 and -0.34 Hz. The differences between PBE0 and PBE results are small (except for the $^{1}J_{HH}$ coupling in $H_2$), however they are a rather poor estimate of the expected deviation from experiment, which can be around 1-2 Hz, based on small molecule benchmarks (*66*). In addition, a calculation with a larger QM region was performed, whereby the Ala175, Thr177, His201, Pro202, Gly203, Cys204, Val205, Cys250, Asp251, and Met252 residues were also included and treated with the pcseg-1 basis set (*67*). Deviations due to the choice of QM region are under 0.3 Hz and fairly systematic. The effect of the QM/MM embedding, as opposed to a CPCM implicit solvent model, is shown in Table S15. Based on all of these calculations, we estimate that the "technical" errors in for the

${}^{1}J_{HH}$ couplings to be below 20 Hz and the "technical" error for long range coupling ${}^{n>1}J_{HH}$ calculations to be no more than 1 Hz, however deviations from experiment might be 2-3 times larger.

The influence of geometry changes in the structural models can be assessed by comparing the NMR parameters computed for models **A** & **C** derived from the crystal structure (*10*) with those for models **E** & **G** derived from the MD-trajectory(*16*). As previously stated, the deviations for the computed chemical shifts and indirect couplings between analogous models (**A6/7/8** vs **E6/7/8** and **C3/4/7/8** vs **G3/4/7/8**, respectively) are much larger than the uncertainties estimated from the benchmarks: Chemical shifts deviate up to 3.6 ppm and couplings up to 9 Hz for these models. Thus, differences in model geometries have a stronger influence on the NMR parameters computed than the different choices of DFT functional, basis set, or treatment of environment used for this this benchmark.

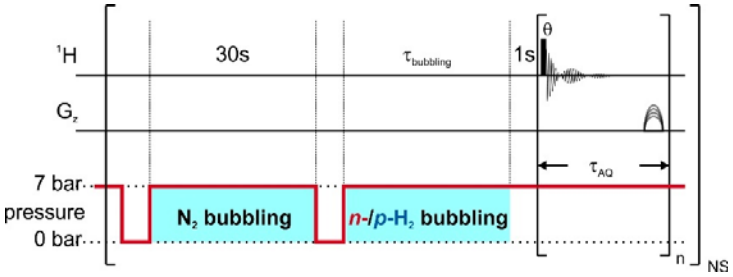


**Fig. S1.**
Excitation and acquisition scheme for $^1$H-NMR spectra series acquired after sample bubbling with $n$-$H_2$ or $p$-$H_2$ (used for Fig. 2A+B of the main article). Typically multiple acquisitions (n) were performed in fixed time intervals $\tau_{AQ}$ after $H_2$ bubbling. The red line indicates the idealized pressure profile throughout the experiment (gauge pressures stated). The filled bar indicates a hard pulse with angle $\theta$. The unfilled half-ellipsoids indicate purge-gradients of slightly varying amplitude for consecutive acquisitions. The pulse- and acquisition phase used was $(n-1)\frac{\pi}{2} + \Phi_{NS}$, where $\Phi_{NS} = \frac{\pi}{2}(0,\ 2,\ 2,\ 0,\ \ 1,\ 3,\ 3,\ 1)$ and where $\Phi_{NS}$ refers to the phase-cycling applied when collecting $n$-$H_2$ isotope exchange kinetics with repeated sample bubbling.

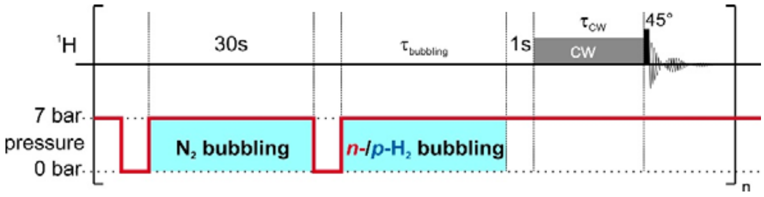


**Fig. S2.**

Excitation and acquisition scheme for PHIP-CEST experiment (used for Fig. 4B of the main article). The red line indicates the idealized pressure profile throughout the experiment (gauge pressures stated). The filled grey bar indicates a continuous-wave pulse applied at the desired CEST offset frequency and the black bar indicates a hard 45° pulse applied in resonance with water signal. Spin-lock and hard pulse were applied with phase 0, and all experiments were collected with a single acquisition per time-point (no phase cycling).

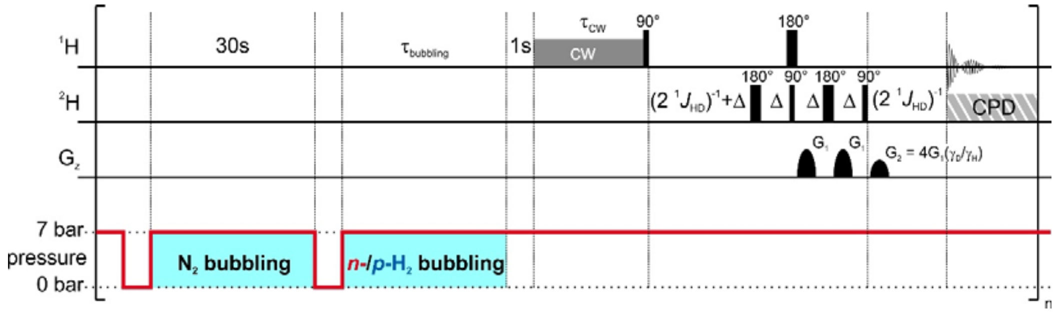


**Fig. S3.**

PHIP-CEST experiment with $^1$H-$^2$H HMQC acquisition filter (PHIP-CEST-HMQC; used for Fig. 4A of the main article). The red line indicates the idealized pressure profile throughout the experiment (gauge pressures stated). The filled grey bar indicates a continuous-wave pulse applied at the desired CEST offset frequency and black bars indicate hard pulses (with different flipping angle) applied in resonance with water signal. Filled half-ellipsoids indicate field gradient pulses and she striped grey block indicates optional heteronuclear decoupling during acquisition. Δ is a sufficiently long period to accommodate a pulsed field gradient, corresponding recovery delay and the delay for *J*-evolution equal to $(2 \cdot {}^1J_{HD})^{-1}$ with ${}^1J_{HD} = 43$ Hz. All pulse phases were 0 and spectra were collected in single scans. The striped grey box indicates GARP.4 composite pulse decoupling on the $^2$H-channel, which was applied during acquisition in some of the experiments.

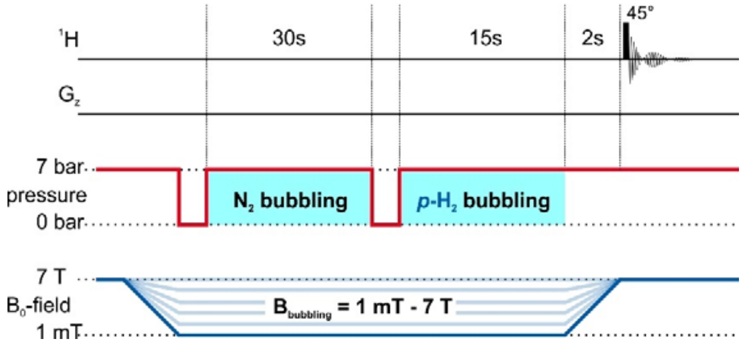


**Fig. S4.**

$^{1}$H-PHIP experiments with manual field-cycling between different low-field positions for $N_2$ and $p$-$H_2$ bubbling and acquisition at 7.05 T. Manual field-cycling was performed between the end of the $p$-$H_2$-bubbling and prior to the $^{1}$H-pulse and took less than 2 s. Idealized $B_0$-field profiles are indicated in blue. The true field profiles were non-linear and subject to the variability inherent to manual operation. An idealized pressure profile is indicated in red. No phase cycling was used, since only single scans wer acquired.

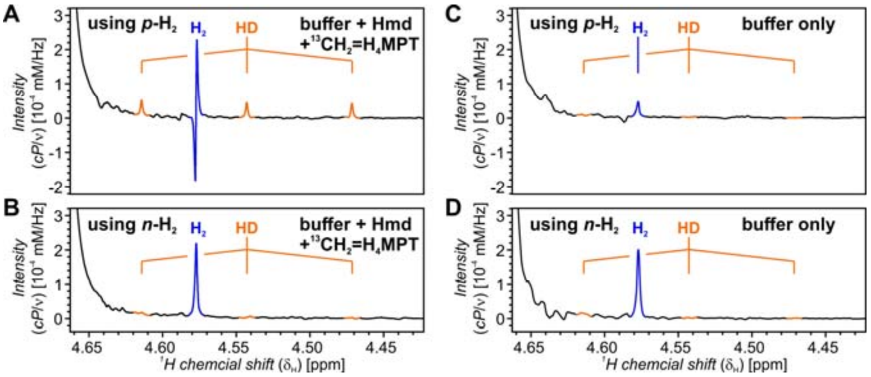


**Fig. S5.**

Test for PHIP effects in absence of Hmd. [1]H signal shapes for hydrogen obtained after bubbling *p*-$H_2$ (87% enrichment, top panels) or *n*-$H_2$ (bottom panels) through either the pure buffer (right panels) or a sample containing the buffer with 1 μM *j*Hmd and 3 μM $^{13}CH_2$=$H_4$MPT added (left panels). The $D_2O$-buffer had deuteration levels of 97% to 98%, a *p*D 6.0 and it contained 1 mM EDTA, 120 mM potassium phosphate.

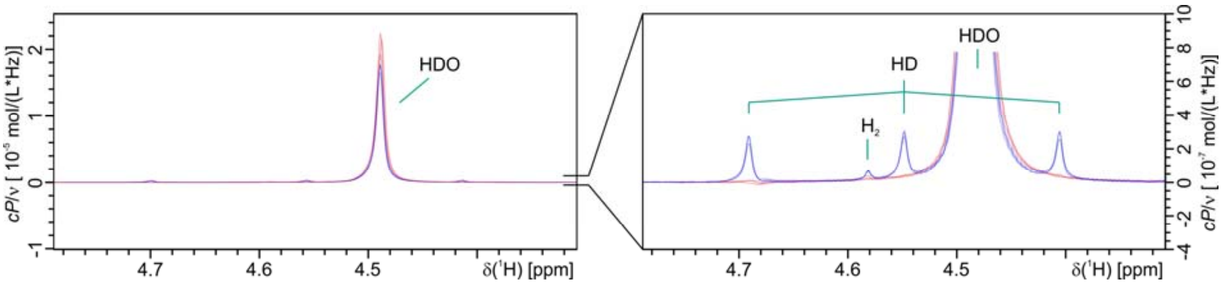


**Fig. S6.**

Spectrum overview and reproducibility of PHIP effects **for one sample.** Overlay of $^{1}$H-NMR spectra obtained after bubbling samples with *n*-$H_2$ (red) or with *p*-$H_2$ (blue) using the pulse sequence shown in Fig. S1. Three overlays are shown for bubbling with each *p*-$H_2$ and *n*-$H_2$. Spectra were acquired at 327.4 K using 11 µM *j*Hmd and 110 µM $^{13}CH_2$=$H_4$MPT in $D_2O$-buffer (98.3% deuteration), *p*D 7.0, 1 µM EDTA, 120 mM potassium phosphate.

The reduction of the water peak intensity, apparent only at high *j*Hmd concentration and high temperature, is visible in the left panel. Spectra are shown again in the 7.0 T row of Fig. S7, and signal integrals are listed the corresponding row in Table S1.

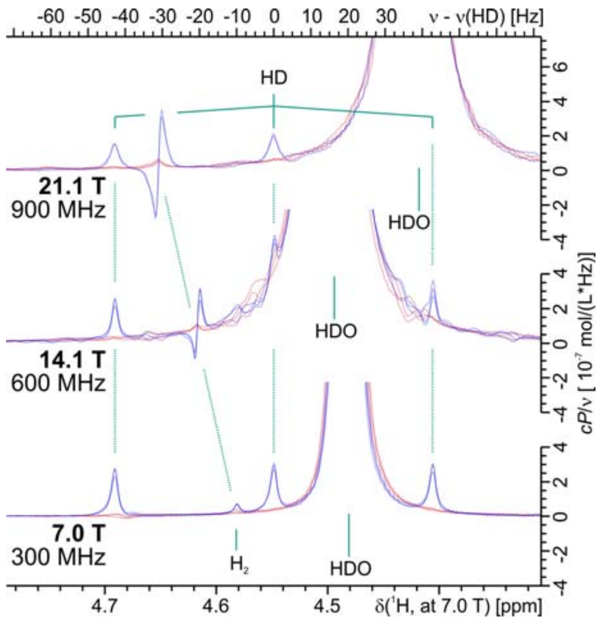


**Fig. S7.**

Reproducibility of PHIP effects **for one sample** and field dependence at high Hmd concentration and temperature. Overlay of $^{1}$H-NMR spectra obtained after bubbling samples with *n*-$H_2$ (red) or with *p*-$H_2$ (red) using the pulse sequence shown in Fig. S1. Spectra are shown for three magnetic field strengths and for all fields three overlays of each *p*-$H_2$ and *n*-$H_2$ spectra are shown. Spectra were acquired at ~325 K (7.0 T: 327.4 K, 14.1 T: 323.3 K, 21.1 T: 324.9 K, as obtained from the HDO peak position) using 11 µM *j*Hmd and 110 µM $^{13}CH_2=H_4MPT$ in $D_2O$-buffer (94.2 – 98.3% deuteration), pD 7.0, 1 mM EDTA, 120 mM potassium phosphate. For easier visualization, the spectra are plotted against a shared frequency scale in Hz (upper horizontal axis), relative to the HD resonance. The chemical shift axis (bottom horizontal axis) is only valid for the 7.0 T data.
The spectra shown for 7.0 T are also shown in Fig. S6.

**Table S1.**

Signal integrals obtained from the spectra shown in Fig. S8. At each magnetic field, **three spectra from the same sample** are overlaid. All integrals are reported as averages ± 1 standard deviation. The integrals for $H_2$ are reported as the integrals over the absolute values of the real part of the spectrum.

| Static magnetic field [*T*] | $P_0$ ($^1$H) [] | Bubbling | Integral (HD) (*cP*) [$10^{-3}$ mM] | Int.(abs(Re)) ($H_2$) (*cP*) [$10^{-3}$ mM] | Integral (HDO) (*cP*) [$10^{-3}$ mM] | Solvent deuteration [%] |
|---|---|---|---|---|---|---|
| 7.0 | 2.20*$10^{-5}$ | *para*-$H_2$ | 1.87 ± 0.12 | 0.10 ± 0.02 | 82 ± 2 | 96.4 ± 0.1 |
| | | *normal* $H_2$ | 0.00 ± 0.04 | 0.01 ± 0.01 | 88.2 ± 0.5 | |
| 14.1 | 4.43*$10^{-5}$ | *para*-$H_2$ | 1.17 ± 0.19 | 0.50 ± 0.09 | 273 ± 3 | 94.3 ± 0.2 |
| | | *normal* $H_2$ | 0.11 ± 0.09 | 0.05 ± 0.03 | 277 ± 4 | |
| 21.1 | 6.65*$10^{-5}$ | *para*-$H_2$ | 1.29 ± 0.08 | 1.21 ± 0.03 | 269 ± 7 | 96.3 ± 0.1 |
| | | *normal* $H_2$ | 0.02 ± 0.06 | 0.13 ± 0.04 | 268 ± 4 | |

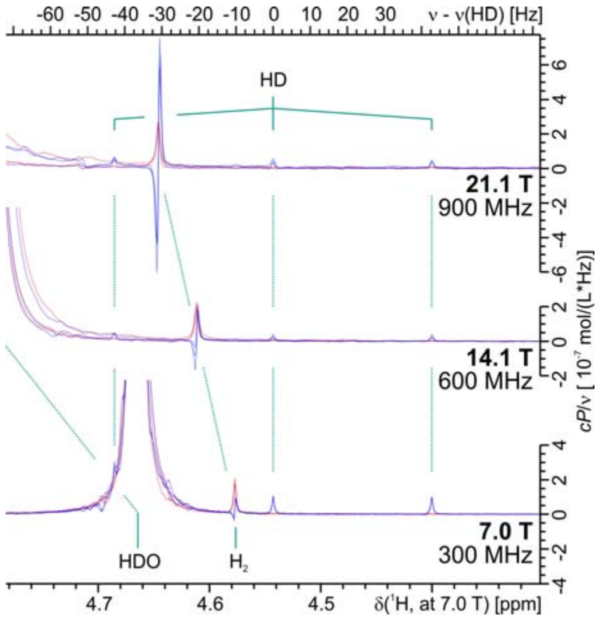


**Fig. S8.**

Reproducibility of PHIP effects **for different samples** and field dependence at low Hmd concentration and temperature. Overlay of $^1$H-NMR spectra obtained after bubbling samples with *n*-$H_2$ (red) or with *p*-$H_2$ (red) using the pulse sequence shown in Fig. S1Fig. S1. Spectra are shown for three magnetic field strengths and for all three fields three overlays of each *p*-$H_2$ and *n*-$H_2$ spectra are shown. At each field, the *p*-$H_2$ spectrum with the best shim was chosen for representation in Fig. 2C of the main article. Spectra were acquired at 309 K using 1 µM *j*Hmd and 3 µM $^{13}CH_2$=$H_4$MPT in $D_2O$-buffer (95.4 – 98.4% deuteration), *p*D 6.0, 1 mM EDTA, 120 mM potassium phosphate. For easier visualization, the spectra are plotted against a shared frequency scale in Hz (upper horizontal axis), relative to the HD resonance. The chemical shift axis (bottom horizontal axis) is only valid for the 7.0 T data.

**Table S2.**

Signal integrals obtained from the spectra shown in Fig. S8. At each magnetic field, **spectra from three different samples** are overlaid. The integrals for the HD and $H_2$ signals are reported as averages ± 1 standard deviation, whereas the HDO signal integral and degrees of deuteration are given as the range obtained in the three different sample preparations. The integrals for $H_2$ are reported as the integrals over the absolute values of the real part of the spectrum.

| Static magnetic field [*T*] | $P_0$ ($^1$H) [] | Bubbling | Integral (HD) (*cP*) [$10^{-3}$ mM] | Int.(abs(Re)) ($H_2$) (*cP*) [$10^{-3}$ mM] | Integral (HDO) (*cP*) [$10^{-3}$ mM] | Solvent deuteration [%] |
|---|---|---|---|---|---|---|
| 7.0 | $2.20*10^{-5}$ | *para*-$H_2$ | 0.26 ± 0.03 | 0.07 ± 0.01 | 82 – 84 | 96.7 – 96.9 |
| | | *normal* $H_2$ | 0.02 ± 0.02 | 0.19 ± 0.02 | | |
| 14.1 | $4.43*10^{-5}$ | *para*-$H_2$ | 0.10 ± 0.04 | 0.25 ± 0.08 | 88 – 183 | 96.4 – 98.3 |
| | | *normal* $H_2$ | 0.00 ± 0.03 | 0.34 ± 0.13 | | |
| 21.1 | $6.65*10^{-5}$ | *para*-$H_2$ | 0.15 ± 0.04 | 0.96 ± 0.05 | 114 – 358 | 95.0 – 98.5 |
| | | *normal* $H_2$ | 0.02 ± 0.03 | 0.37 ± 0.04 | | |

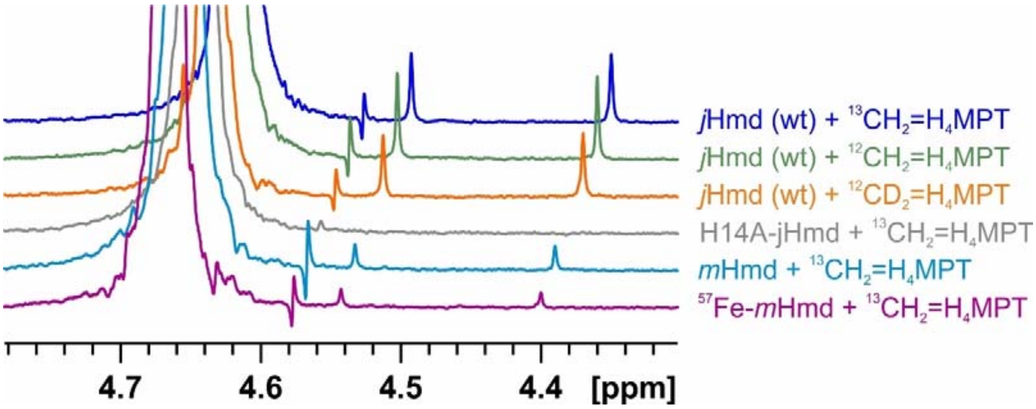


**Fig. S9.**

[1]H-PHIP experiments performed with differential isotopic labeling or with the H14A-mutation. The top three rows show that there is no significant impact of differential isotope labeling at the methylene-group of methylene-$H_4$MPT on the [1]H-PHIP effects. Row #4 vs row #1 (from top) shows the disappearance of the [1]H-PHIP effects of *j*Hmd, if the His14Ala mutation is introduced. The bottom two rows show, that $^{57}$Fe-labeling of the catalytically active iron has no significant impact on the [1]H-PHIP effects. Spectra are shown without scaling to the measured sample activities. Activity-normalized HD-PHIP intensities are given in Table S3. Samples were prepared from with 1 µM Hmd and 3 µM methylene-$H_4$MPT in $D_2O$-buffer (95.4 – 98.4% deuteration), *p*D 7.0, 1 mM EDTA, 120 mM potassium phosphate. Spectra were acquired at 309 K, after bubbling with *p*-$H_2$ of 99% enrichment.

**Table S3.**

Summary of the PHIP experiments with different isotope labeling schemes, as well as with the H14A mutant of *j*Hmd. **N**: Number of samples prepared. $\boldsymbol{v_0}$: activity for hydrogen isotope exchange. ***Int*(HD-PHIP)**: Integrals observed for the HD PHIP signals. ***Int*(HD-PHIP)/$\boldsymbol{v_0}$**: Integrals for the HD PHIP, scaled by the activity of the individual samples. Average values and standard deviations over all samples are reported. All experiments were performed at 7.0 T (300 MHz) and 309 K. Samples were prepared from 1 µM Hmd and 3 µM methylene-$H_4$MPT in $D_2O$-buffer (98.4 – 99.3% deuteration), *p*D 7.0, 1 mM EDTA, 120 mM potassium phosphate. The *p*-$H_2$ enrichments varied between 85% and 99%.

| | | N | $v_0$(309 K) [µmol min$^{-1}$ (mg protein)$^{-1}$] | *Int*(HD-PHIP) [mM (for 100% p$H_2$)] | *Int*(HD-PHIP)/$v_0$ [mM (for 100% p$H_2$)/(U/mg)] |
|---|---|---|---|---|---|
| *j*Hmd (wt) | $^{13}CH_2$=$H_4$MPT | 5 | 30 ± 9 | 11 ± 3 | 0.39 ± 0.11 |
| *j*Hmd (wt) | $^{12}CH_2$=$H_4$MPT | 4 | 43 ± 11 | 16 ± 6 | 0.37 ± 0.05 |
| *j*Hmd (wt) | $^{12}CD_2$=$H_4$MPT | 4 | 30 ± 7 | 12 ± 3 | 0.40 ± 0.07 |
| H14A-*j*Hmd | $^{13}CH_2$=$H_4$MPT | 3 | 0.3 ± 0.2 | not detected | not detected |
| *m*Hmd (wt) | $^{13}CH_2$=$H_4$MPT | 4 | 49 ± 10 | 4.2 ± 1.2 | 0.085 ± 0.011 |
| $^{57}$Fe-*m*Hmd | $^{13}CH_2$=$H_4$MPT | 5 | 32 ± 9 | 2.8 ± 0.6 | 0.090 ± 0.007 |

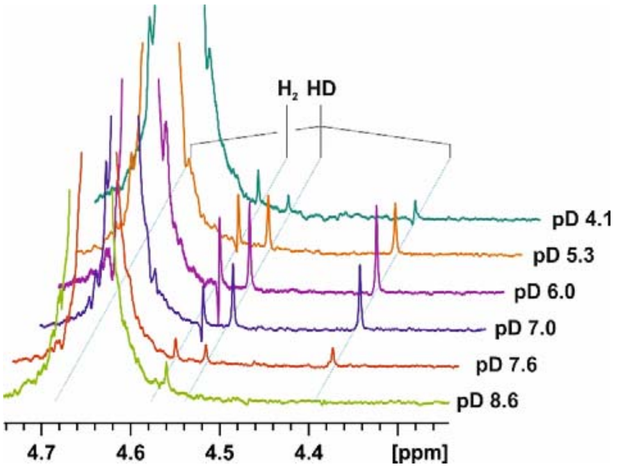


**Fig. S10.**

Dependence of the PHIP effects on the buffer pD. The hydrogen region of [1]H-PHIP experiments is shown. The residual water peak at 4.66 ppm has been truncated for better representation. Data was collected at 7.05 T and 309 K after bubbling *p*-$H_2$ (87% enrichment) through samples containing buffers of the corresponding pD with 1 µM *j*Hmd and 3 µM $^{13}CH_2$=$H_4$MPT added. The $D_2O$-buffers contained 1 mM EDTA and 120 mM potassium phosphate and were adjusted to the corresponding pD, using an electrode calibrated for pH-measurements and estimating the pD from the pH-meter reading pH*, using pD = pH* + 0.41 (*68*).

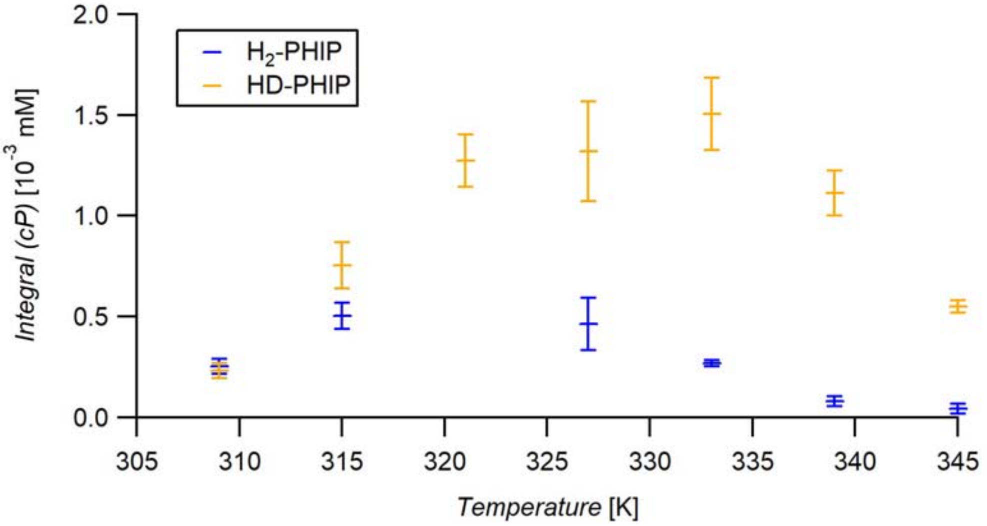


**Fig. S11.**

Dependence of the PHIP effects on temperature. Data was collected at 14.1 T after bubbling *p*-$H_2$ (87% enrichment) through samples containing 11 µM *j*Hmd and 110 µM $^{13}CH_2$=$H_4$MPT. The $D_2O$-buffer contained 1 mM EDTA and 120 mM potassium phosphate and was adjusted to pD 7.0. For HD, the integral over the real part of the spectrum is shown, for $H_2$, the integral over the absolute value of the real part of the spectrum is shown. Around 320 K the residual water peak of the buffer overlaps with the $H_2$ signal, so no reliable integration is possible here.

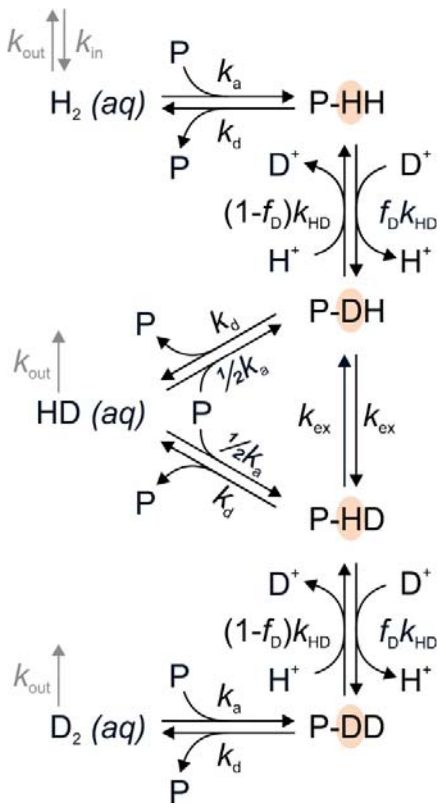


**Fig. S12.**

Kinetic model used for fitting the hydrogen isotope exchange catalyzed by Hmd for the extraction of kinetic parameters. The positions of isotope exchange are highlighted in red. Note that $k_{a1}$ is a second-order rate constant. $f_D$ is the fraction of deuteration of the buffer. Mass transport from the gas phase to the aqueous phase (reactions indicated in grey) was assumed to only occur during bubbling, and neglected thereafter.

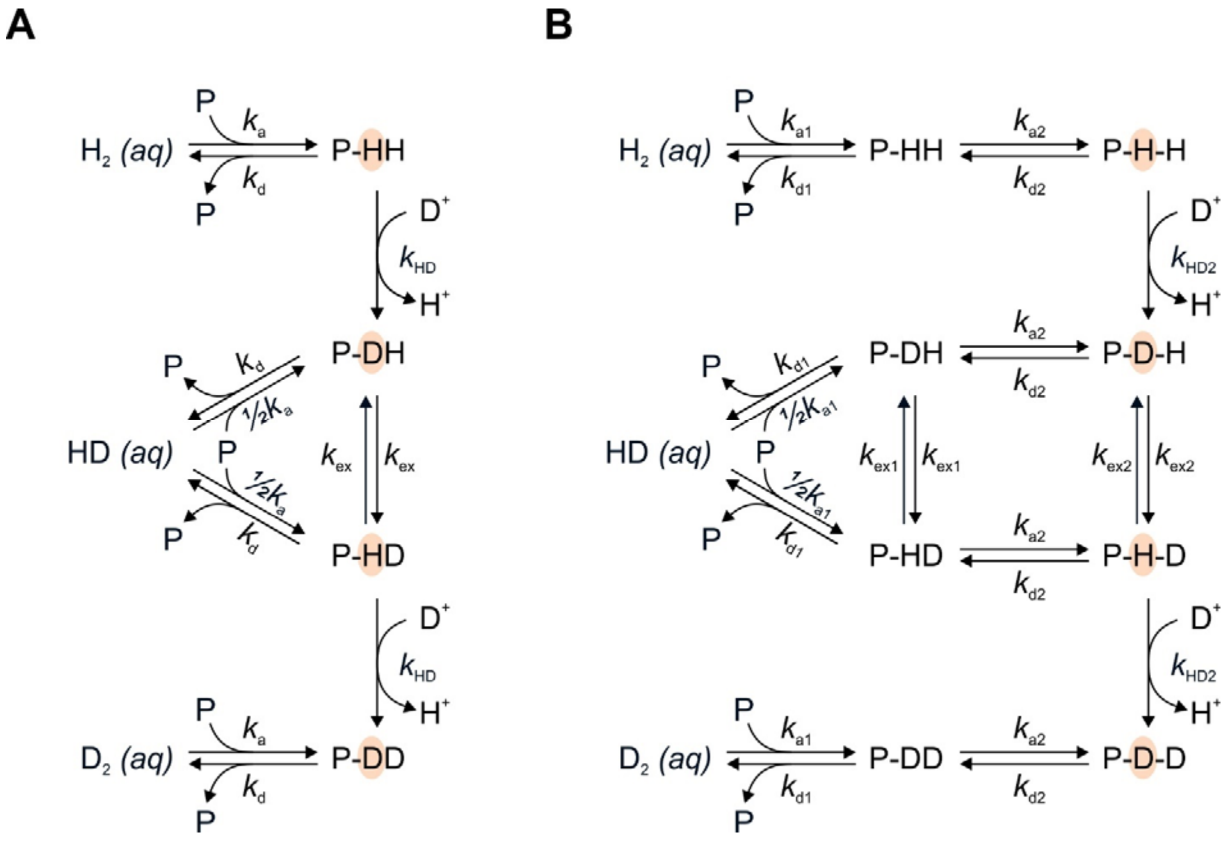


**Fig. S13.**

Kinetic models used to obtain expressions for $K_m$ and $v_{max}$ from steady state analysis. **A)** Model with one bound state geometry, **B)** Model with two bound state geometries.

**Table S4.**

Best fit values obtained from fitting the kinetic constants $k_a$, $k_d$ and $k_{HD}$ to the $H_2 \rightleftarrows HD \rightleftarrows D_2$ kinetics catalyzed by jHmd. Samples contained 1 µM *j*Hmd and 3 µM $^{13}CH_2$=$H_4$MPT in $D_2O$-buffer (1 mM EDTA, 120 mM potassium phosphate, pD 6.0). The menten constants $K_m$ and $v_{max}$ with respect to the substrate $H_2$ are reported.
Data was fit using multiple datasets acquired with varying hydrogen partial pressure $p(H_2)_0$ during bubbling. Data was acquired according to Fig. S1, with predefined mixtures of $N_2$ and *n*-$H_2$ being used during the second bubbling period. All experiments were performed at the same pressure ($p(H_2)_0 + p(N_2)_0$ = 7 bars), to retain the same bubbling characteristics for all $p(H_2)_0$. Data fitting was performed for multiple test values of $k_{ex}$, which were fixed during fitting. The results for the best fitting $k_{ex}$ are given in the top row, together with a lower bound estimate $k_{ex,min}$ for the rate of site exchange. The bottom row lists the best fit values for the limit $k_{ex} = \infty$. Menten constants obtained from the fitted values of $k_a$, $k_d$ and $k_{HD}$ are further listed.

| $k_a$ [$mM^{-1}$ $s^{-1}$] | $k_d$ [$s^{-1}$] | $k_{HD}$ [$s^{-1}$] | $k_{ex}$ [$s^{-1}$] | $k_{ex,min}$ [$s^{-1}$] | $K_m$ [mM] | $v_{max}$ [U/mg] |
|---|---|---|---|---|---|---|
| (40 ± 16)[a] | 35 ± 2 | 257 ± 26 | 300 | 90 ± 10 | 0.9 ± 0.4 | 92 ± 10 |
| (45 ± 16)[a] | 37 ± 2 | 178 ± 13 | ∞ | - | | |

a: Sample activities decayed during sample storage, handling and during longer experiments. The reduced concentration of active enzyme leads to a downscaling of the apparent $k_a$, whereas $k_d$ and $k_{HD}$ remain unaffected. Values reported for $k_a$ should represent the values of fully active samples. These were extrapolated from the fitted $k_a$ values, using the measured sample activities.
For all simulations, the apparent association constants $k_a$' was chosen such that the isotope exchange kinetics, measured on the same sample immediately before or after the PHIP experiments, were reproduced, rather than using the best fit values listed here.

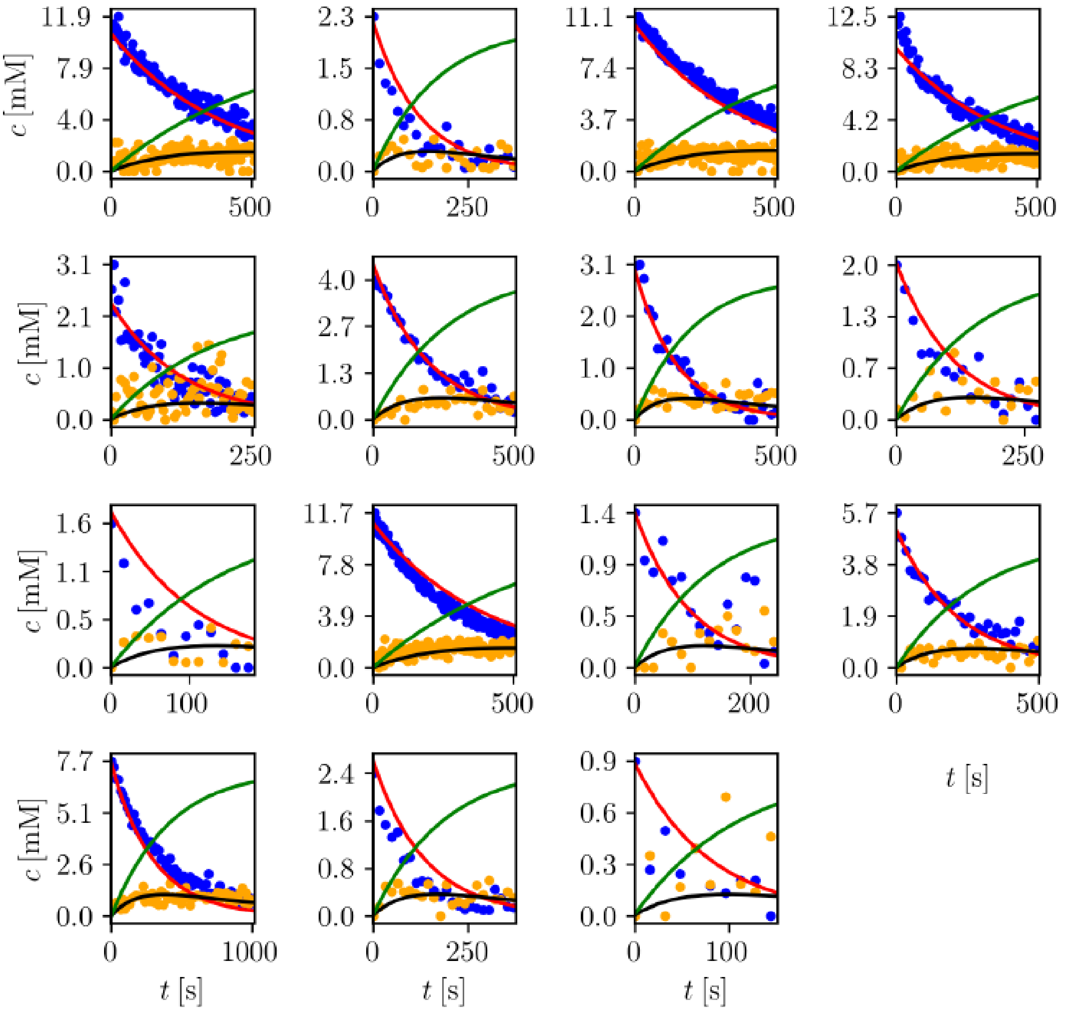


**Fig. S14.**

Simulated vs. experimental hydrogen isotope exchange kinetics measured for varying hydrogen partial pressure during bubbling $p(H_2)_0$. The figure shows experimental concentraions of $H_2$ (blue) and HD (orange) and simulated kinetic profiles for $H_2$ (red), HD (black) and $D_2$ (green) using the best-fit results obtained, assuming $k_{ex} = 300\ s^{-1}$.

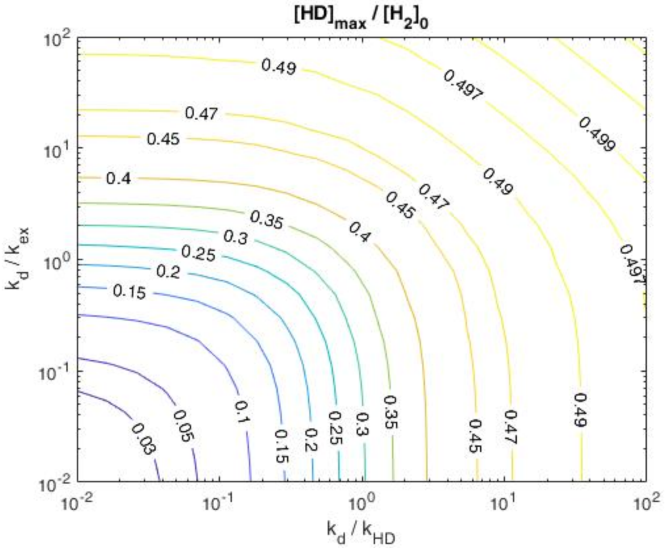


**Fig. S15.**

Contours for different exemplary ratios of $[HD]_{max}/[H_2]_0$, obtained from equation ( 23 ). The ratio $[HD]_{max}/[H_2]_0$ is experimentally easily available and allows a straightforward estimate of upper bounds of the ratios $k_d/k_{HD}$ and $k_d/k_{ex}$. Experimentally we observe $[HD]_{max}/[H_2]_0 \approx 0.1$.

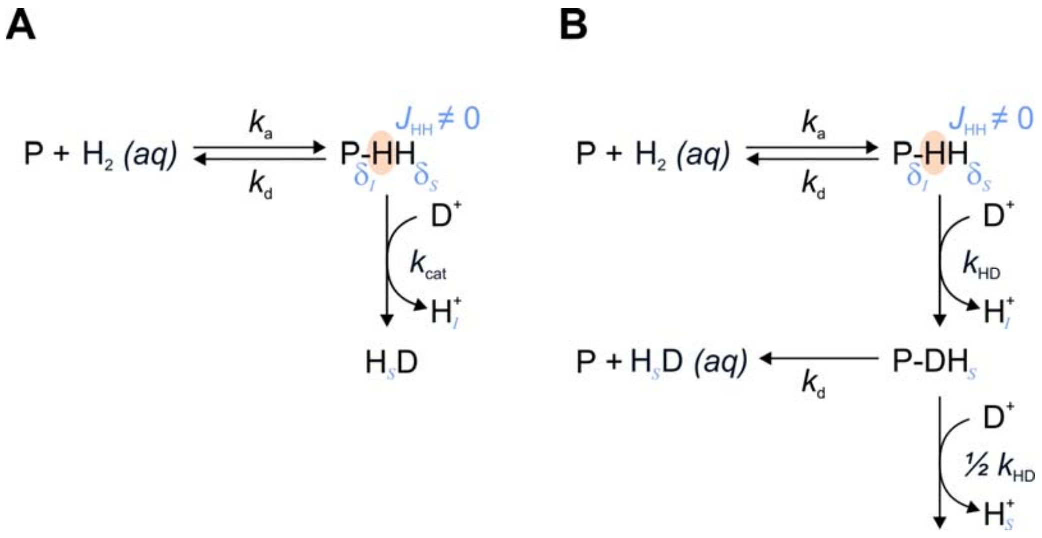


**Fig. S16.**
Models used for the analytical description of HD-PHIP effect in section 2.4. The hydrogen positions undergoing the isotope exchange reaction are highlighted in orange. Hydrogens are labeled as *I* and *S*, according to the chemical shift positions they occupy in intermediate P-HH.

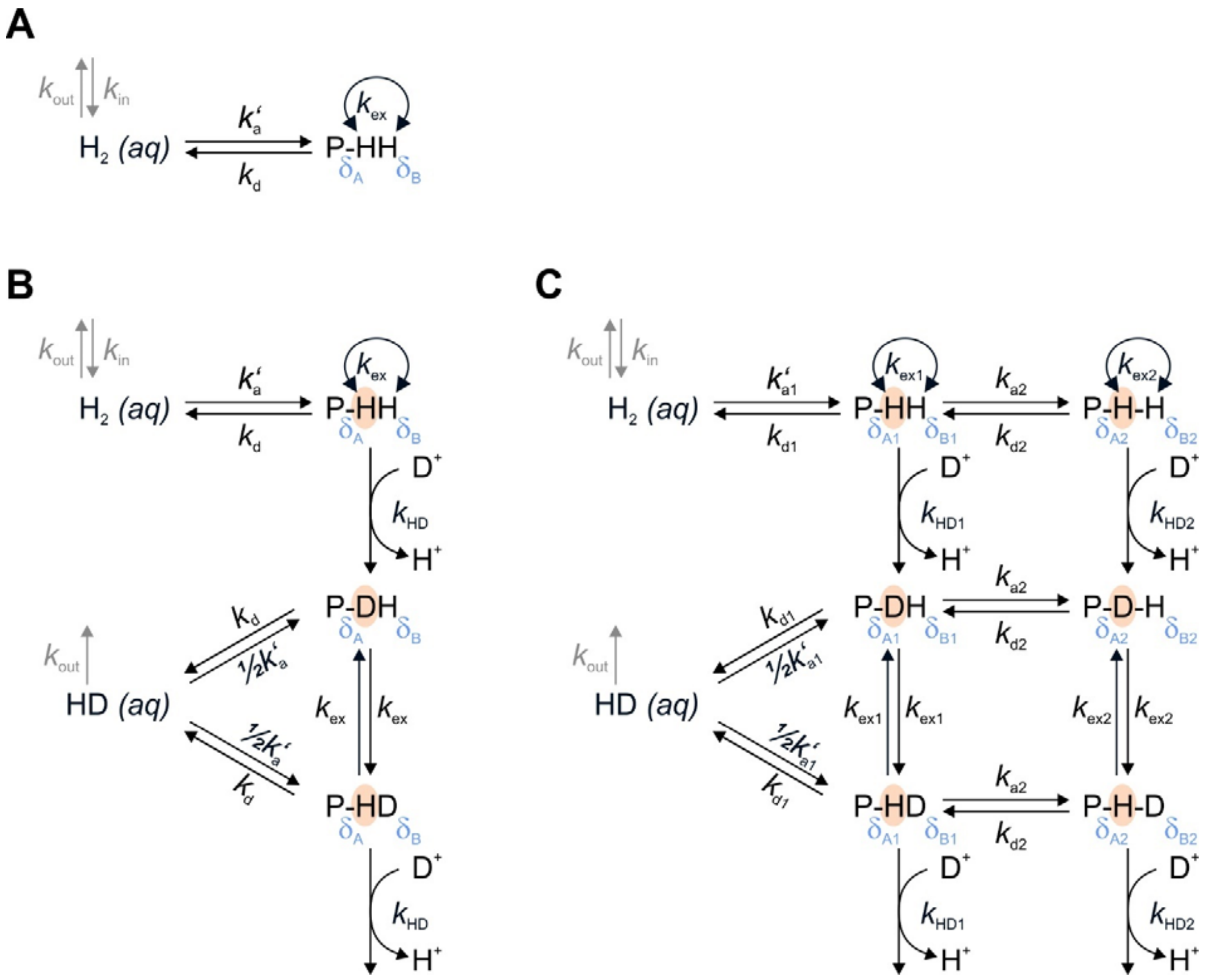


**Fig. S17.**

Models used for numerically simulating combined nuclear spin dynamics and chemical kinetics. **(A)** Model treating only reversible association to one bound state geometry, used for fitting the PNL only. **(B)** Model assuming one bound state geometry. **(C)** Model assuming two bound state geometries.

Note that association was treated as pseudo-first-order reaction for numeric spin dynamics simulations ($k'_{a1} = k_{a1}[P]$). The hydrogen positions undergoing the isotope exchange reaction are highlighted in orange. These models are used in sections 2.6 – 2.10.

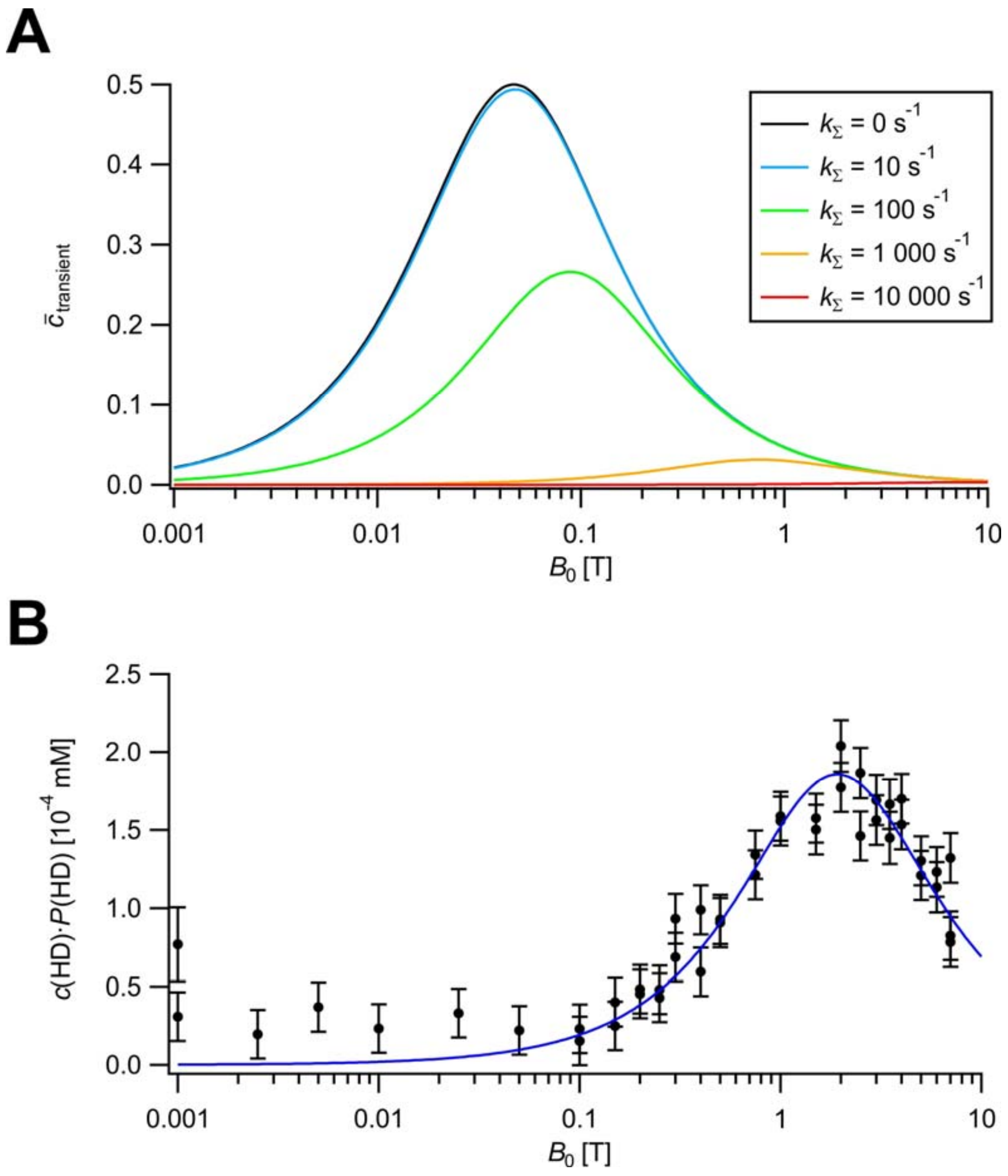


**Fig. S18.**

(**A**) Exemplary plots of $\bar{c}_{transient}$ from equation ( 66 ) for different inverse lifetimes ($k_\Sigma = \tau^{-1}$) of the bound state. Plots are for an exemplary spin system with $J_{HH} = \pm 5\ Hz$ and $\Delta\delta = \mp 5\ ppm$. (**B**) HD-PHIP intensity observed in manual field cycling experiments according to Fig. S4 as a function of the $B_0$ field at which bubbling was performed. Signal integrals are plotted in the field-independent $c{\cdot}P$ scale (*concentration·polarization*) described in section 1.12. The best fit solution assuming a model with coherent spin evolution in a transiently formed intermeditate described in section 2.4 (equations ( 83 ) and ( 84 )) is shown in blue. Fitting parameters and results are described in section 2.5.

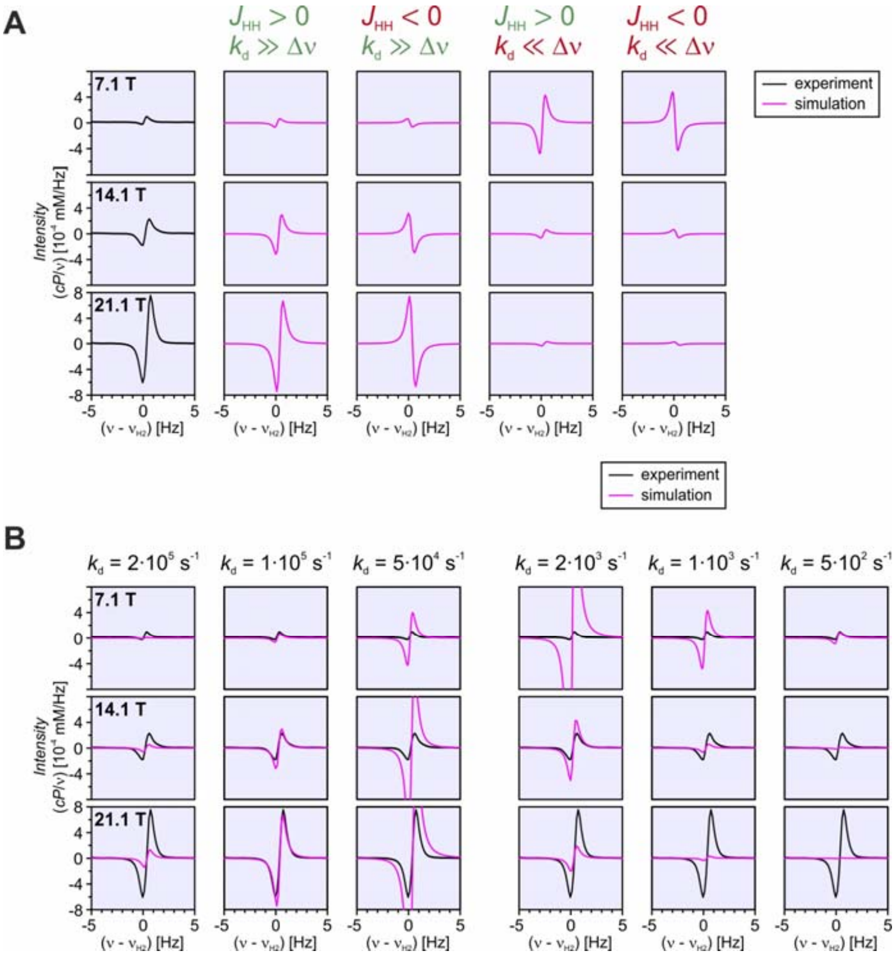


**Fig. S19.**

Exemplary simulated PNL lineshapes (magenta) compared with the experimental spectra (black). (**A**) Sign and field dependence for different signs of $J_{HH}$, in the limit of slow or fast dissociation. (**B**) Signal intensity trends for varying $k_d$ in the limit of fast (left) or slow (right) dissociation. Simulations were performed using the model in Fig. S17A, using $\delta_A = -1\ ppm, \delta_B = +9\ ppm, J_{HH} = +220 Hz, k_a' = k_d/6000, k_{ex} = k_d$. In panel (**A**) $k_d = 10^5$ for plots with $k_d \gg \Delta\nu$ and $k_d = 10^3$ for plots with $k_d \ll \Delta\nu$.

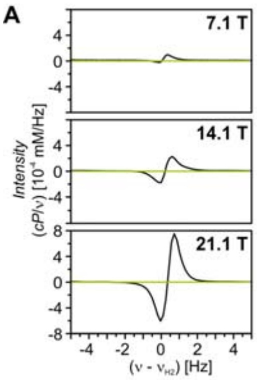


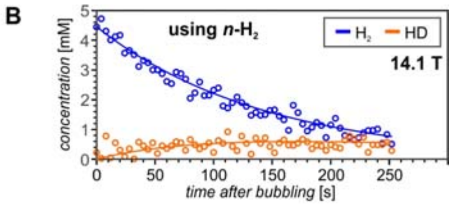


$k_d = 35\ s^{-1}$
$k'_a = 8 * 10^{-3} s^{-1}$ ($k_a \approx 48\ mM^{-1}\ s^{-1}$)
$k_{HD} = 257\ s^{-1}$
$k_{ex} = 300\ s^{-1}$

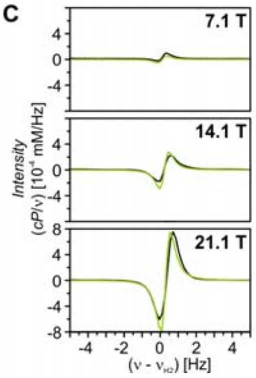


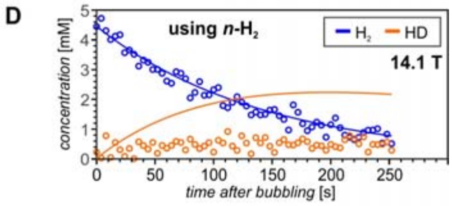


$k_d = 3.5 * 10^5\ s^{-1}$
$k'_a = 80\ s^{-1}$ ($k_a \approx 48 * 10^5\ mM^{-1}\ s^{-1}$)
$k_{HD} = 30.8\ s^{-1}$
$k_{ex} = 300\ s^{-1}$

**Fig. S20.**

Illustration of the incompatibility of the PNL effect with the kinetic model with one bound state geometry (Fig. S17A). (**A**) & (**B**) Simulation results using the rate constants extracted from net isotope exchange kinetics (see Table S4; $k'_a$ adjusted to the activity of the sample). (**C**) & (**D**) Simulation with $k_d$ and $k'_a$ increased to simultaneously reproduce the measured PNL effect and the $H_2$ consumption kinetics.

The $H_2$ region of simulated (green) and measured (black) $^1$H-PHIP spectra is shown in panels (**A**) & (**C**). The measured (oben cycles) and simulated (solid lines) hydrogen isotope exchange kinetics are shown in panels (**B**) & (**D**).

Simulations shown assume the model with one bound state geometry (Fig. S17A), with $J_{HH} = 220\ Hz$, $\delta_{av} = 4.3\ ppm$ and $|\Delta\delta| = 10.0\ ppm$. Other simulation parameters are as listed in Table S9.

**A**

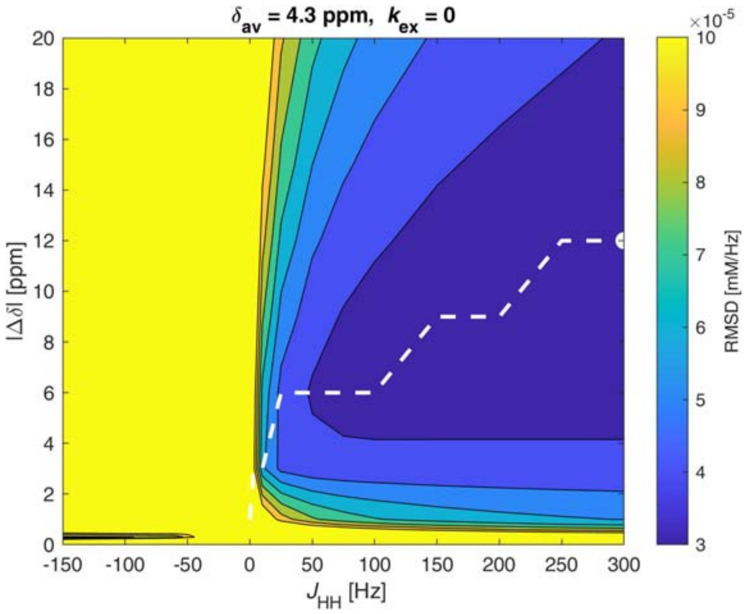


**B**

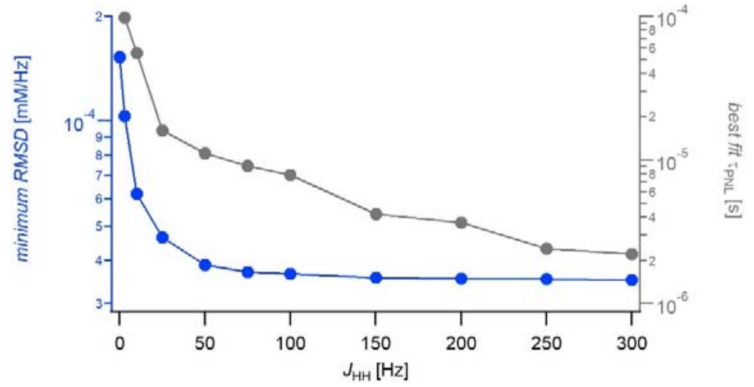


**Fig. S21.**

**A)** Exemplary parameter scan over possible Δδ and *J* values for the intermediate causing the observed PNL effect. Plotted is the RMSD between the measured and the simulated data over three spectra, collected at 7.05 T, 14.10 T and 21.15 T. The parameter scan providing the best fit found is shown ($\delta_{av} = 4.3\ ppm, k_{ex} = 0$). The positions of lowest RMSD found is indicated by a white circle.

**B)** Blue: Minimum RMSD found for a given value of *J*, for the parameter scan shown in A). The dotted line in A) indicates the position of the minimum RMSD path in the Δδ - *J* plane. Grey: Bound state lifetimes obtained for the minimum RMSD found for a given value of *J*.

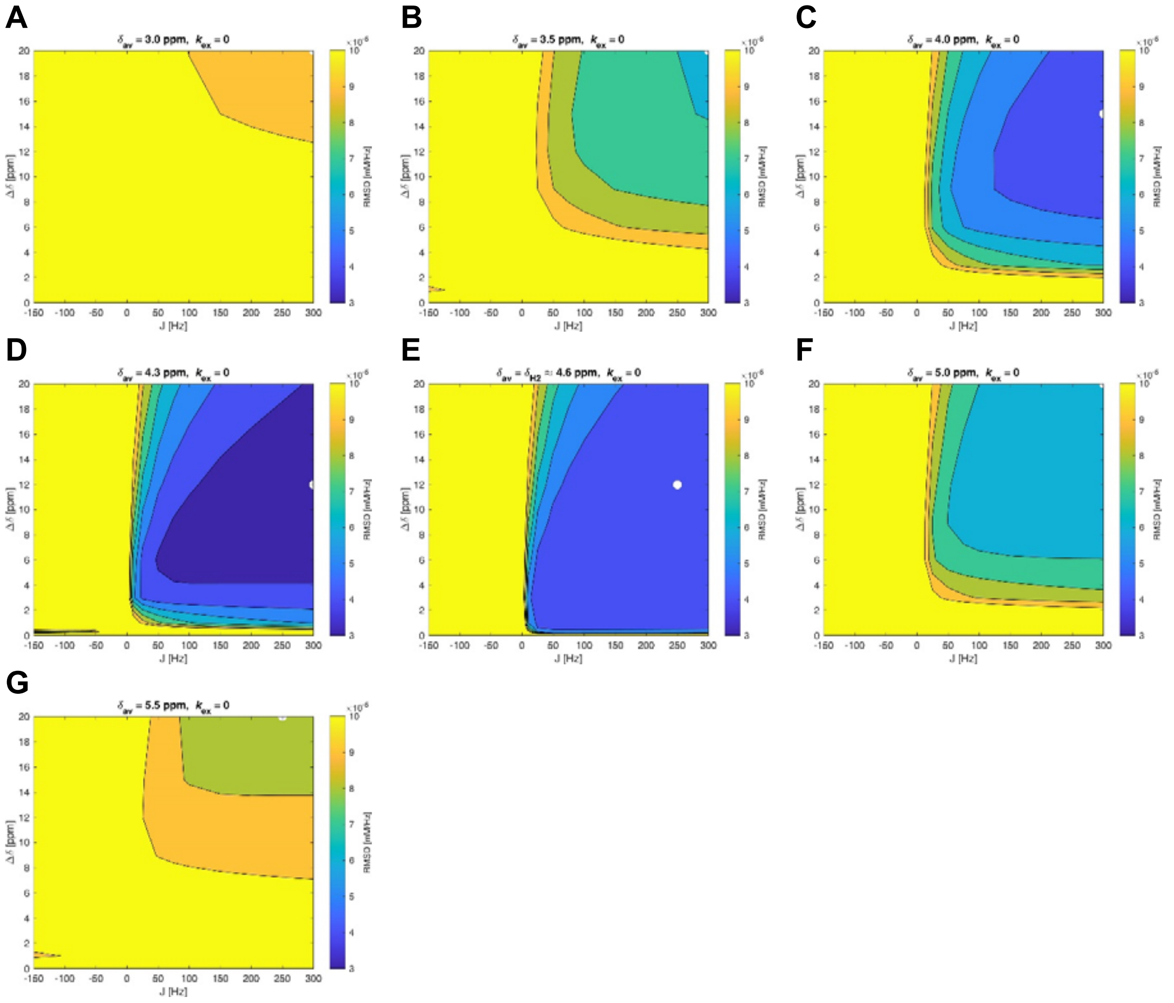


**Fig. S22.**

Parameter scans over possible Δδ and *J* values for the intermediate causing the observed PNL effect for different choices of the averaged chemical shift $\delta_{av}$. Plotted is the RMSD between the measured and the simulated data over three spectra, collected at 7.05 T, 14.10 T and 21.15 T. The positions of lowest RMSD found is indicated by a white circles.

**A**

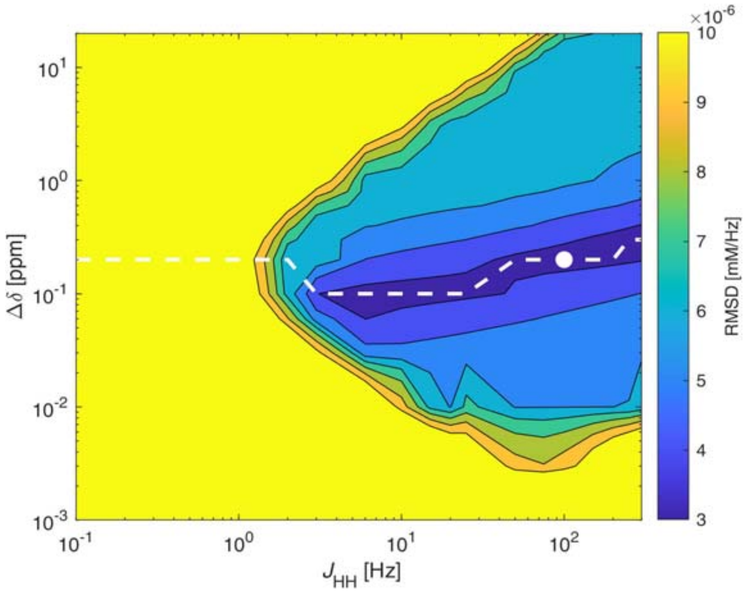


**B**

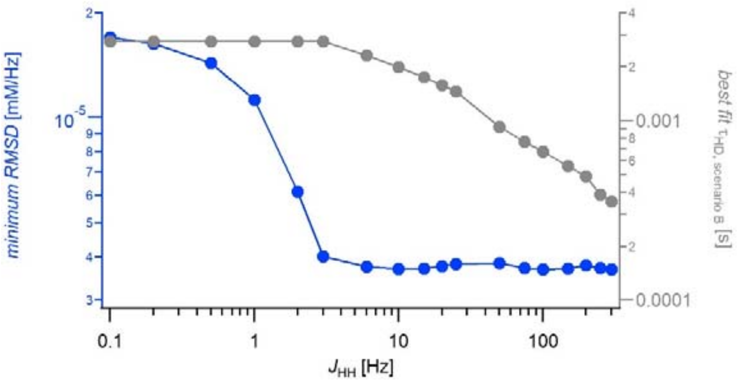


**Fig. S23.**

Systematic parameter scans for HD-PHIP creation according to scenario B (creation of the HD-PHIP via a non-zero *J*-coupling and $H^+ \rightarrow D^+$ isotope exchange both in ensemble 2, see section S2.9).

**A)** Parameter scan over possible Δδ and *J* values. Plotted is the RMSD between the measured and the simulated data over three spectra, collected at 7.05 T, 14.10 T and 21.15 T. The position of lowest RMSD found is indicated by a white circle and the dotted white line indicates the path of minimum RMDS for a given value of $J_{HH}$, for which the RMSD is plotted in panel **B**.

**B)** Blue: Best fit RMSD found for a given value of *J*, for the parameter scan shown in panel **A**. Grey: Bound state lifetime for the best fit solution at the given value of *J*. The upper limit $\tau_{\text{HD, scenario B}} \leq 2.8\ ms$ results from $k_d = 35\ s^{-1}$, $k_{HD} = 257\ s^{-1}$ and $k_{ex} \geq 70\ s^{-1}$ (see Table S4 and section 2.9).

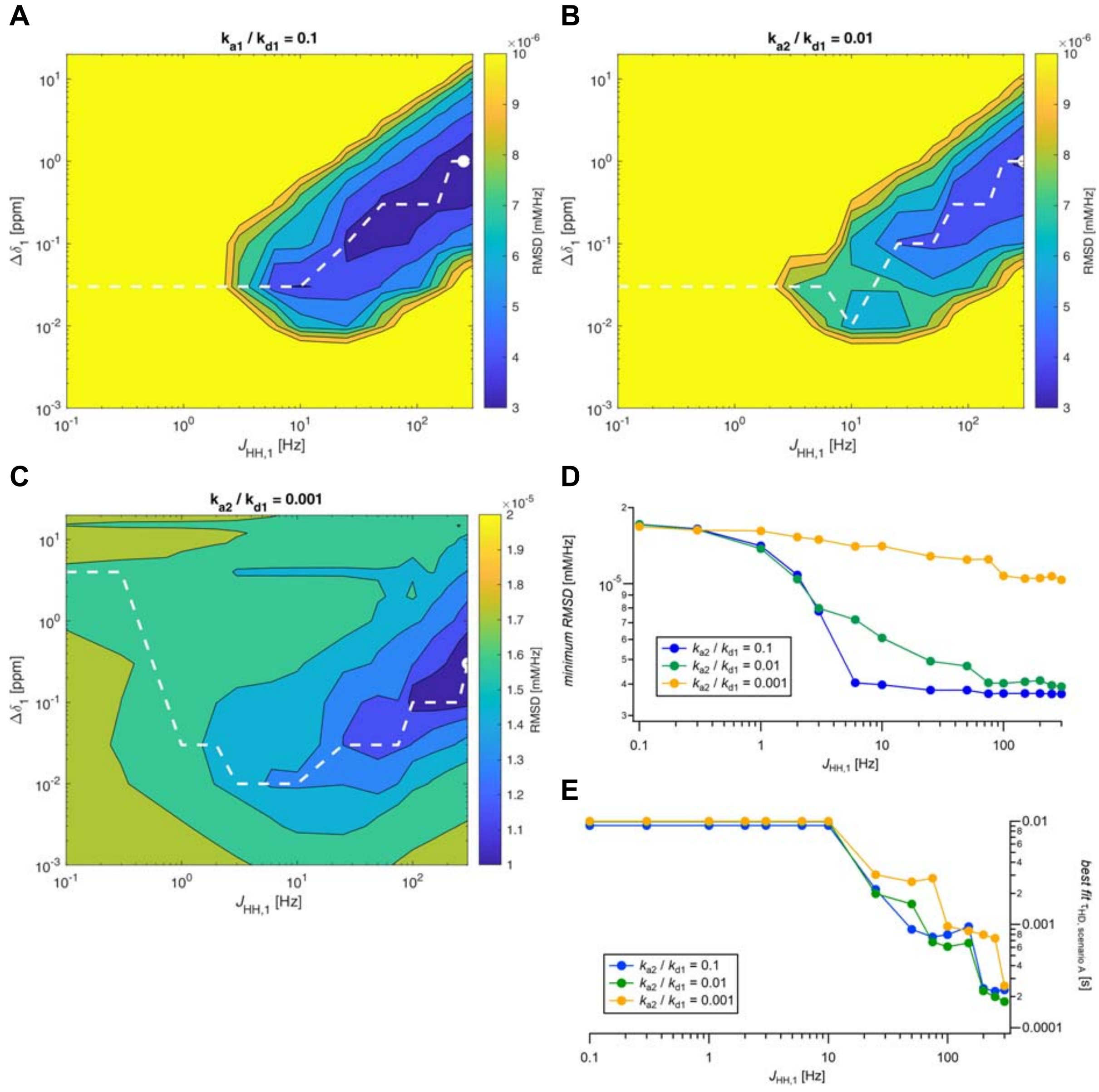


**Fig. S24.**

Systematic parameter scans for HD-PHIP creation according to scenario A (creation of the HD-PHIP via a non-zero *J*-coupling in ensemble 1; $H^+ \rightarrow D^+$ isotope exchange in ensemble 2, see section S2.9). **A - C)** Parameter scans over possible Δδ and *J* values for different choices of the ratio $k_{a2}/k_{d1}$. the intermediate causing HD-PHIP effect. Plotted is the RMSD between the measured and the simulated data over three spectra, collected at 7.05 T, 14.10 T and 21.15 T. The positions of lowest RMSD found is indicated by a white circle and the dotted white lines indicate the path of minimum RMDS for a given value of $J_{HH}$, for which the RMSDs are plotted in panel **D**. For panel **C**, mind the different colorscale used. **D)** Best fit RMSD found for a given value of *J*, for the parameter scan shown in panels **A** - **C**. **E)** Lifetimes of ensemble 1 (P-HH) for the best fit results. A lower limit constraint of $k_{d1} \geq 100s^{-1}$ was used during the scan.

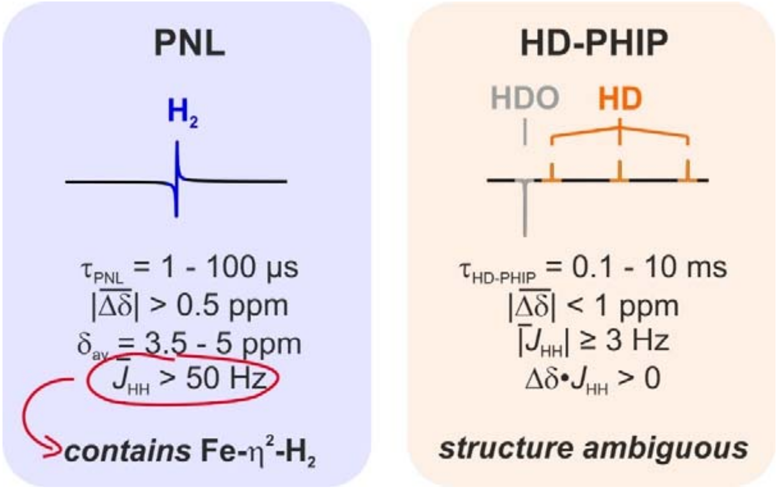


**Fig. S25.**
Summary of the parameter ranges compatible with the observed PNL and HD-PHIP signals at 309 K shown in Fig. 2C of the main text. Description of the method of estimating these parameter ranges is provided in sections S2.8 and S2.9.

**Table S5.**

Fixed simulation parameters assumed for numeric spectrum simulations.

| Paramter name | *Fixed value* | Parameter description |
|---|---|---|
| $k_{in}$ | $10\ mMs^{-1}$ | Rate of *p*-$H_2$ supplied to the solution during bubbling |
| $k_{out}$ | $2\ s^{-1}$ | Rate of hydrogen escaping the solution to the gas phase during bubbling |
| $\delta(H_2)$ | $4.57728\ ppm$ | Chemical shift of free $H_2$ dissolved in the buffer |
| $\delta(HD)$ | $4.54277\ ppm$ | Chemical shift of free HD dissolved in the buffer |
| $J_{HH}(H_2)$ | $278.2\ Hz$ | *J*-coupling within free $H_2$ dissolved in the buffer |
| $J_{HD}(HD)$ | $42.83\ Hz$ | *J*-coupling within free HD dissolved in the buffer |
| $T_1(H_2)$ | $2\ s$ | $T_1(\,^1H)$ of free $H_2$ dissolved in the buffer |
| $T_S(H_2)$ | $500\ s$ | Singlet lifetime in free $H_2$ dissolved in the buffer |
| $T_1(HD)$ | $8\ s$ | $T_1(\,^1H)$ of free HD dissolved in the buffer |
| $T_1(\text{bound})$ | $1\ s$ | $T_1(\,^1H)$ of bound-state hydrogen species (P-HH, P-HD, P-DH, P-H-H, P-H-D, P-D-H) |

**Table S6.**

Simulation parameters of the $^1$H-spectra shown in Fig. 2 of the main article (scenario A), as well as other models reproducing the $^1$H-spectra. Simulations were performed in the two bound-state model shown in Fig. S17C. Additional parameters fixed in all simulations can be found in Table S5.

| | *Scenario A* | Scenario B1 | *Scenario B2* |
|---|---|---|---|
| Paramter name | Ensemble 1: Fe-η$^2$-$H_2$ Ensemble 2: fully dissociated | Ensemble 1: Fe-η$^2$-$H_2$ Ensemble 2: small positive $J_{HH2}$ | Ensemble 1: Fe-η$^2$-$H_2$ Ensemble 2: small negative $J_{HH2}$ |
| $k'_{a1}$ | $1.0\ s^{-1}$ | $10\ s^{-1}$ | $10\ s^{-1}$ |
| $k_{d1}$ | $1 * 10^4\ s^{-1}$ | $1.7 * 10^5\ s^{-1}$ | $1.7 * 10^5\ s^{-1}$ |
| $k_{ex1}$ | $1 * 10^3\ s^{-1}$ | $1 * 10^3\ s^{-1}$ | $1 * 10^3\ s^{-1}$ |
| $k_{HD1}$ | $0\ s^{-1}$ | $0\ s^{-1}$ | $0\ s^{-1}$ |
| $\tau_1$ | $90\ \mu s$ | $5.8\ \mu s$ | $5.8\ \mu s$ |
| $\delta_{mean1}$ | $\delta(H_2) = 4.57728\ ppm$ | $4.0\ ppm$ | $\delta 4.0\ ppm$ |
| $\Delta\delta_1$ | $+0.90 ppm$ | $+10.0 ppm$ | $+10.0 ppm$ |
| $\delta_{A1}$ | $4.12728\ ppm$ | $-1\ ppm$ | $-1\ ppm$ |
| $\delta_{B1}$ | $5.02728\ ppm$ | $+9\ ppm$ | $+9\ ppm$ |
| $J_{HH1}$ | $+250\ Hz$ | $+220\ Hz$ | $+220\ Hz$ |
| $k_{a2}$ | $100\ s^{-1}$ | $170\ s^{-1}$ | $170\ s^{-1}$ |
| $k_{d2}$ | $30\ s^{-1}$ | $30\ s^{-1}$ | $30\ s^{-1}$ |
| $k_{ex2}$ | $1 * 10^2\ s^{-1}$ | $1 * 10^2\ s^{-1}$ | $1 * 10^2\ s^{-1}$ |
| $k_{HD2}$ | $3 * 10^2\ s^{-1}$ | $3 * 10^2\ s^{-1}$ | $3 * 10^2\ s^{-1}$ |
| $\tau_2$ | $2.3\ ms$ | $2.3\ ms$ | $2.3\ ms$ |
| $\delta_{av2}$ | $7.25\ ppm$ | $3.975\ ppm$ | $3.975\ ppm$ |
| $\Delta\delta_2$ | $-6.5 ppm$ | $+0.05 ppm$ | $-0.05 ppm$ |
| $\delta_{A2}$ | $10.5\ ppm$ | $3.95\ ppm$ | $4.00\ ppm$ |
| $\delta_{B2}$ | $3.0\ ppm$ | $4.00\ ppm$ | $3.95\ ppm$ |
| $J_{HH2}$ | $0\ Hz$ | $+5\ Hz$ | $-15\ Hz$ |

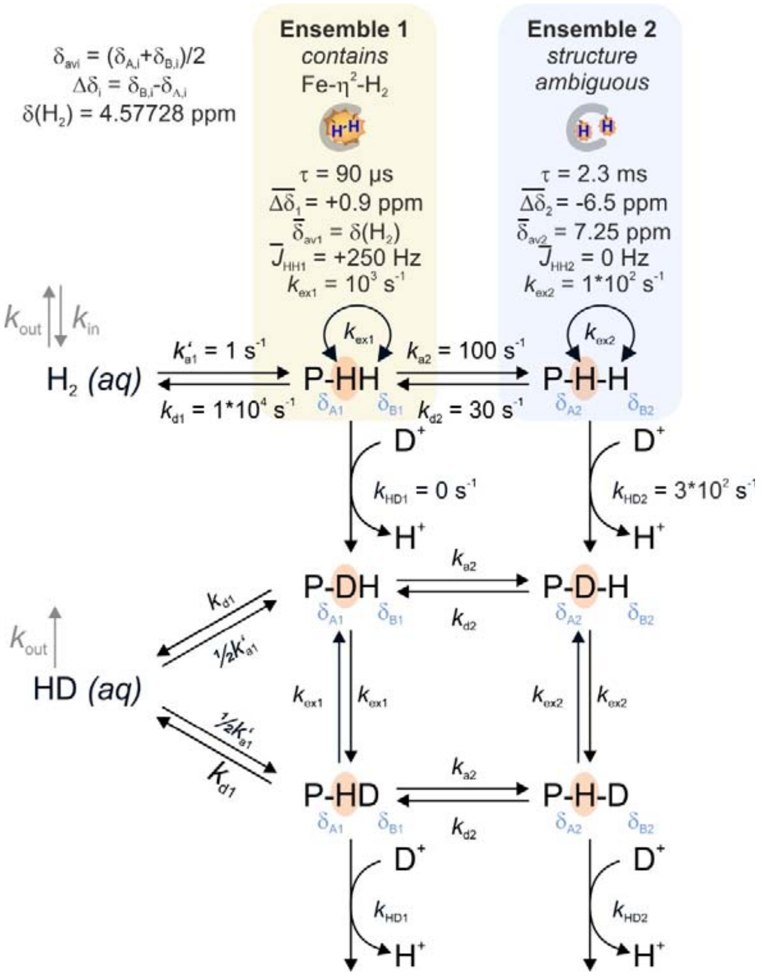


**Fig. S26.**
Kinetic scheme including modeling parameters for the simulated spectra shown in Fig. 2C of the main article.

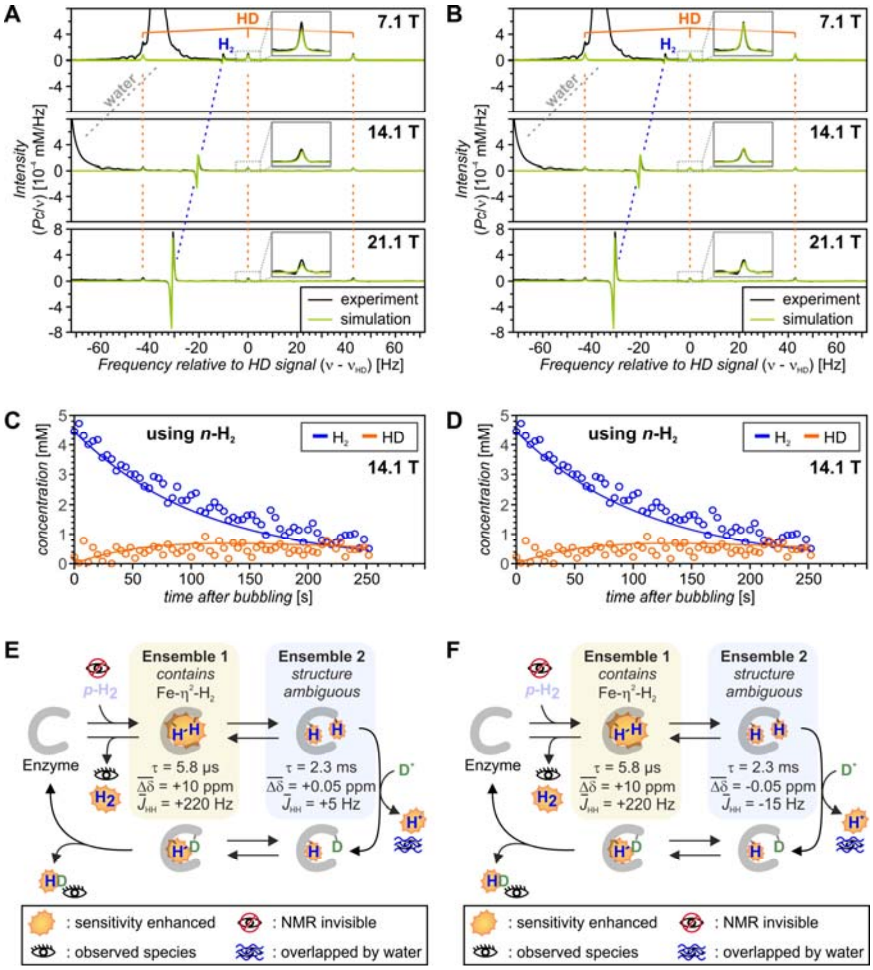


**Fig. S27.**

Simulation results using modeling scenarios B1 (panels **A**, **C** & **E**) and B2 (panels **B**, **D** & **F**). For description of the modeling scenarios, see chapter 2.9 and Table S6.

Top panels show the simulated spectra (green) overlayed with the experimentally observed spectra (black). The respective figure for modeling scenario A is shown in Fig. 2C of the main article. Mittdle panels show overlays of the simulated isotope exchange kinetics with the kinetics measured at 14.1 T. Bottom panels show the short summaries of the modeling comparable to Fig. 2D of the main article.

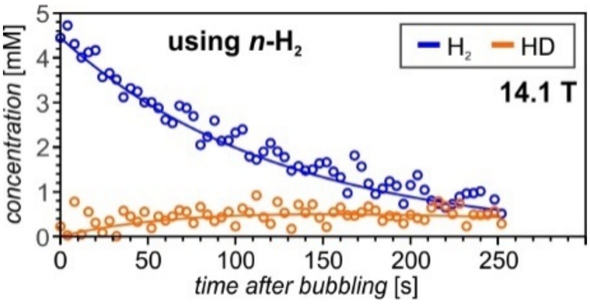


**Fig. S28.**

Kinetics of $H_2$-consumption by $H_2 \rightleftarrows HD \rightleftarrows D_2$ exchange measured immediately after collecting the [1]H-PHIP spectra shown in Fig. 2A&B of the main article. The spectrum shown in Fig. 2B of the main article is the first point ($t = 1s$) of this series of spectra. Data was collected at 14.1 T and 309 K, after bubbling *n*-$H_2$ through a sample containing 1 µM *j*Hmd and 3 µM $^{13}CH_2$=$H_4$MPT in $D_2O$-buffer (pD 6.0, 1 mM EDTA, 120 mM potassium phosphate, degree of deuteration: 98.2%). The solid lines show the simulated kinetics, using the same kinetic parameters as assumed for spectrum simulations in Fig. 2C of the main article (kinetic model assumed shown in Fig. S17C; parameters as listed in Table S6 (scenario A): $k'_{a1} = 1\ s^{-1}, k_{d1} = 1 * 10^4\ s^{-1}, k_{ex1} = 10^3\ s^{-1}, k_{HD1} = 0\ s^{-1}, k_{a2} = 100\ s^{-1}, k_{d2} = 30\ s^{-1}, k_{ex2} = 100\ s^{-1}, k_{HD2} = 300\ s^{-1}$).

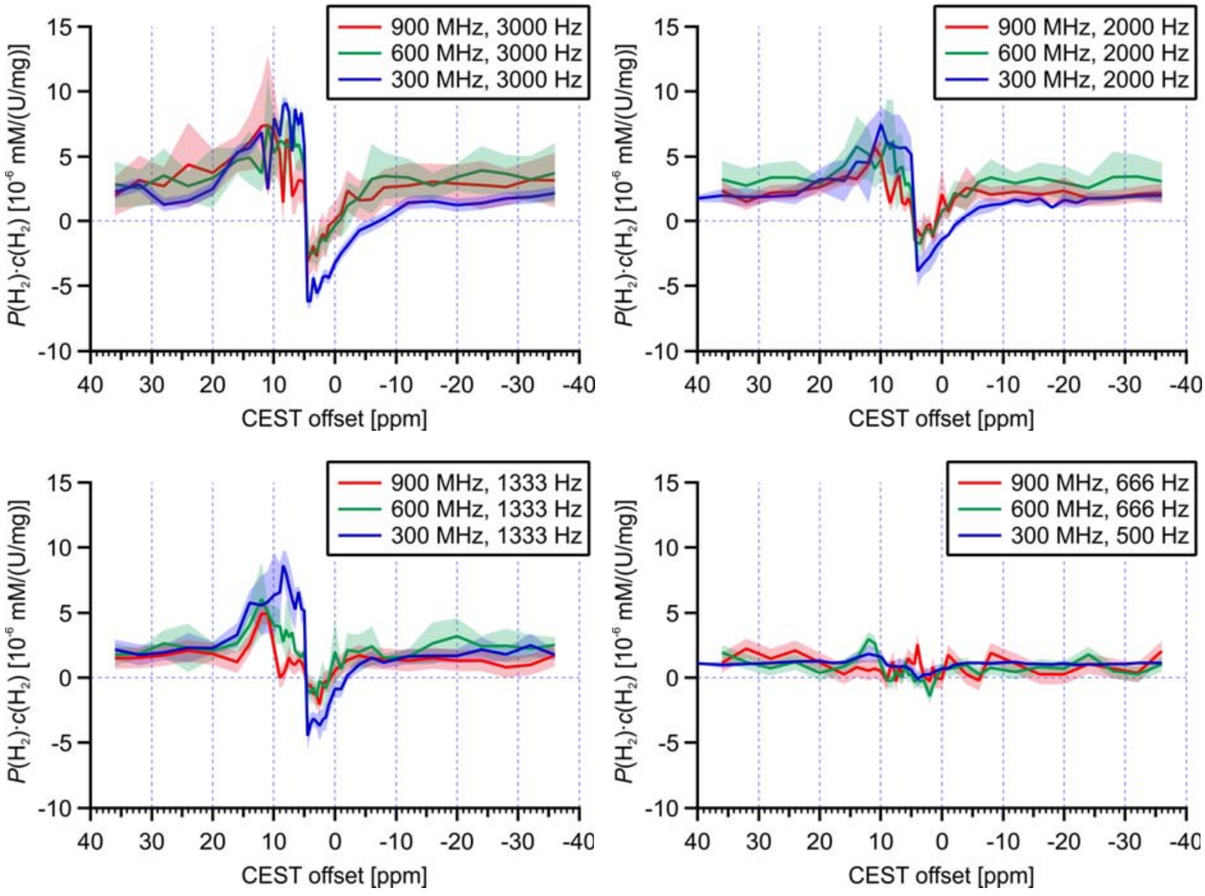


**Fig. S29.**

PHIP-CEST effect measured at the $H_2$ line position at three different fields (red: 21.1 T, 900 MHz, green: 14.1 T, 600 MHz, blue: 7.0 T, 300 MHz). Plotted is the signal integral of the $H_2$-line as a function of CEST offset. The vertical axes are presented on the same (field-independent) scale (polarization*concentration, normalized by sample activity). Solid lines represent the average integrals measured and transparent regions correspond to the estimated uncertainty.

All data was acquired with spin-locking for 2 s, using the experiment shown in Fig. S2. Experiments were performed with different spin locking field ($\gamma B_1$ = 3000 Hz, 2000 Hz, 1333 Hz, 666 Hz or 500 Hz), as stated in the legends. The *p*-H2 content ranged from 85% to 90% during the experiments.

Spectra were acquired at 309 K using 1 µM *j*Hmd and 3 µM $^{13}CH_2$=$H_4$MPT in $D_2O$-buffer (96.4 – 98.6% deuteration), *p*D 6.0, 1 µM EDTA, 120 mM potassium phosphate. Sample activities ranged from 82 U/mg to 28 U/mg at the start of the experiment and typically decayed to 80% of its starting value during the experiments. A linear correction was applied to account for activity loss.

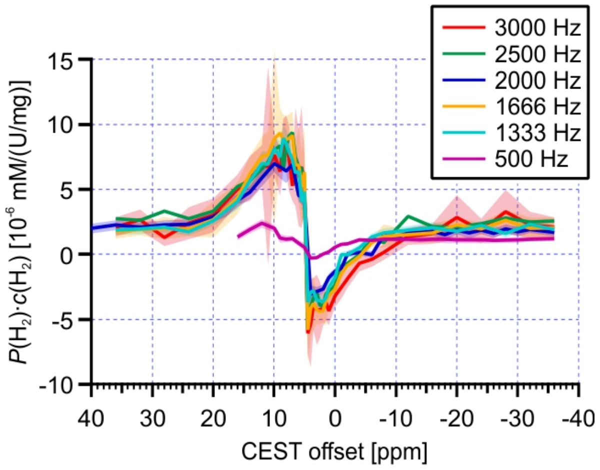


**Fig. S30.**

Spin-lock field dependence of the CEST profiles acquired at 7.0 T (300 MHz) using the pulse sequence shown in Fig. S2 with 2 s spin locking with variable spin-lock field strength ($\gamma_H B_1$). PHIP-CEST profiles were acquired at 309 K using 1 µM *j*Hmd and 3 µM $^{13}CH_2$=$H_4$MPT in $D_2O$-buffer, *p*D 6.0, 1 µM EDTA, 120 mM potassium phosphate.

Sample activities ranged from 72 U/mg to 28 U/mg at the start of the experiment and typically decayed to 80% of its starting value during the experiments. A linear correction was applied to account for activity loss.

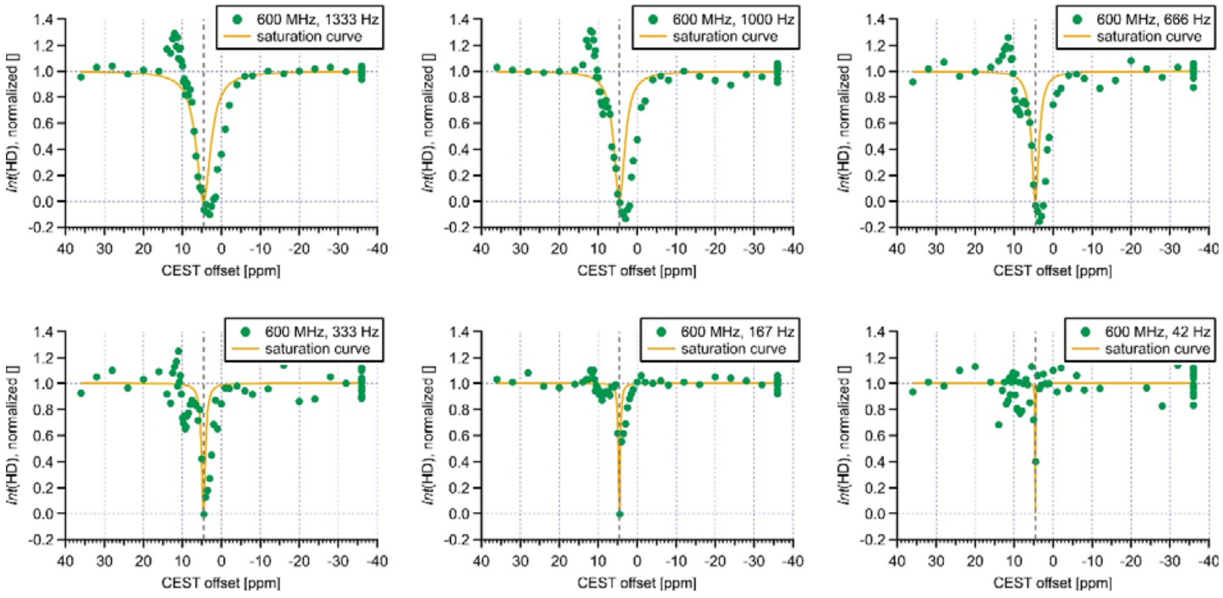


**Fig. S31.**

CEST-curves acquired with varying CEST field-lock strength, using the HMQC-filtered PHIP-CEST experiment (Fig. S3). All data was acquired at 309 K and 14.1 T (600 MHz). Samples contained 1 µM *j*Hmd, 3 µM $^{13}CH_2=H_4MPT$ and 1 mM EDTA in a 120 mM potassium phosphate buffer at *p*D 6.0.

CEST curves were acquired in a shuffled fashion and with interleaving multiple reference experiments with the offset placed at -36 ppm with the experiments at the other frequencies. The experiments at -36 ppm were used to monitor sample activity decay during the experiment. The sample activity profile was fitted to a decaying monoexponential function with offset and the measured integrals were normalized with this decay function.

For easier visualization of the central dip asymmetry, the expected resonance curve for irradiation with negligible radiation damping ($M_z = M_0\left\{1 - \frac{\omega_1^2}{\omega_1^2+(1/T_1)^2+(\omega-\omega_0)^2}\right\}$) (*69*) is overlaid as solid orange line and the on-resonance frequency of HD is highlighted with a dotted grey line.

**Table S7.**

Simulation parameters of the PHIP-CEST curves shown in Fig. 4 of the main article. Simulations were performed in the one bound-state model shown in Fig. S17B. Additional parameters fixed in all simulations can be found in Table S9.

| Paramter name | *value* |
|---|---|
| $k_a'$ | $0.01\ s^{-1}$ |
| $k_d$ | $3 * 10^2\ s^{-1}$ |
| $k_{ex}$ | $1 * 10^3\ s^{-1}$ |
| $k_{HD}$ | $2 * 10^3\ s^{-1}$ |
| $\delta_{av}$ | $\delta(H_2) = 7.35\ ppm$ |
| $\Delta\delta$ | $-6.3\ ppm$ |
| $\delta_A$ | $10.5\ ppm$ |
| $\delta_B$ | $4.2\ ppm$ |
| $J_{HH}$ | $-10\ Hz$ |

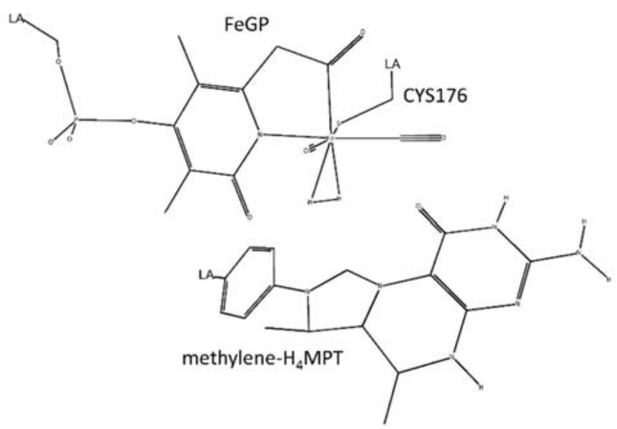


**Fig. S32.**
QM regions used for the QM/MM calculation. For computations using the crystal structure based models E & G, the sidechain of His14 was included in the QM region, whereas it was excluded for the MD derived models A & C from ref. (*16*). Hydrogen link atoms are denoted with LA. Hydrogen atoms bonded to carbon are not shown.

**Table S8.**

Labels of the "2H" optimized models of the [Fe]-hydrogenase active site. Successfully optimized models are marked with "yes". The notation "→X" means that the given model converged instead to structure X during optimization. See section S2.12 for details.

| | **Label** | **A** | **E** | **Corresponding structure in Fig. 3 of main article** |
|---|---|---|---|---|
| 1 | $FeH_2$ | yes | →E6 | **3** |
| 2 | OHtoH-FeH | yes | →E6 | **4** |
| 3 | OHaway-FeH | →A7 | →E7 | **4** |
| 4 | SHtoO-FeH | →A8 | | |
| 5 | SHaway-FeH | →A9 | | |
| 6 | OHtoH-$CH_2$ | yes | yes | **5** |
| 7 | OHaway-$CH_2$ | yes | yes | **5** |
| 8 | SHtoO-$CH_2$ | yes | yes | |
| 9 | SHaway-$CH_2$ | yes | yes | |
| | Source | Ref. (*16*), fig. 9 | Ref. (*10*), PDB: 6hav | |

**Table S9.**

[1]H chemical shifts (in ppm) and couplings (in Hz) for different structures with two hydrogens included in the active site. Results shown in the right column (**E6**-**E9**) are obtained from QM/MM optimized models derived from the closed-state crystal structure (PDB: 6HAV) published in ref. (*10*). Results shown in the left column (**A1**-**A9**) are obtained from the QM/MM optimized MD-snapshot presented in Fig. 9 of ref. (*16*). For an overview of the models with three hydrogens in the active site, see Table S8. Chemical shifts were computed at the TPSS/pS2 and *J*-couplings – at the PBE0/pJ2 level, both with electrostatic embedding. The relative energy with respect to **A/E6**(OHtoH-$CH_2$) for models **A/E**, respectively, at the TPSS-D3BJ/def2-TZVP/MM level is given in kcal·mol$^{-1}$. In the figures, shifts are shown in orange and couplings in teal.

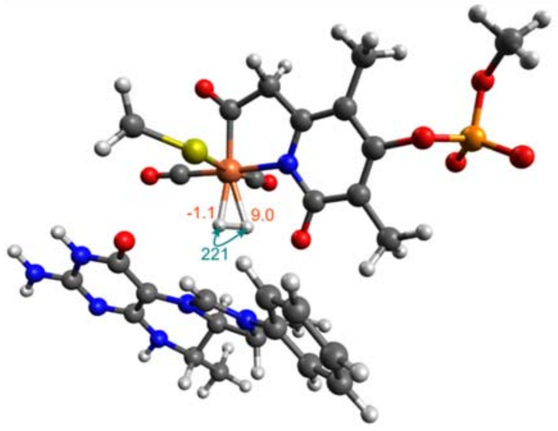


**A1(FeH$_2$)**

| | | |
|---|---|---|
| **Δ*E*** | | 11.3 |
| ***δ*** | Fe-$H_2$ (δ+) | 9.03 |
| | Fe-$H_2$ (δ-) | -1.13 |
| ***J*** | Fe-$H_2$/Fe-$H_2$ | 221.5 |
| | CH/Fe-$H_2$ (δ+) | 1.72 |
| | CH/Fe-$H_2$ (δ-) | 0.39 |

**Table S9 (continued)**

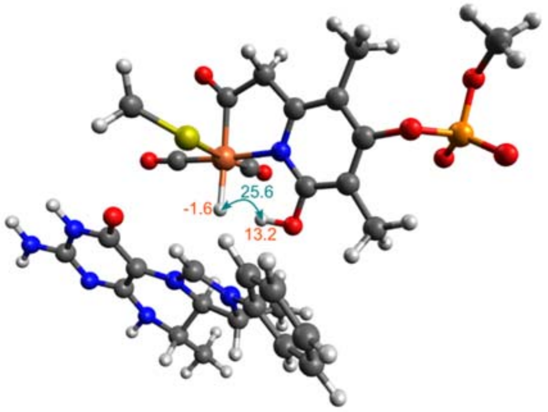


| | | A2(OHtoH-FeH) |
|---|---|---|
| **Δ*E*** | | 10.1 |
| ***δ*** | Fe-H | -1.59 |
| | OH | 13.21 |
| ***J*** | OH/Fe-H | 25.59 |
| | CH/Fe-H | -3.24 |
| | OH/CH | 0.71 |

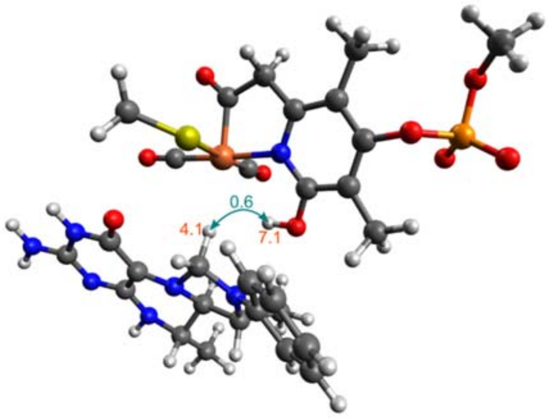


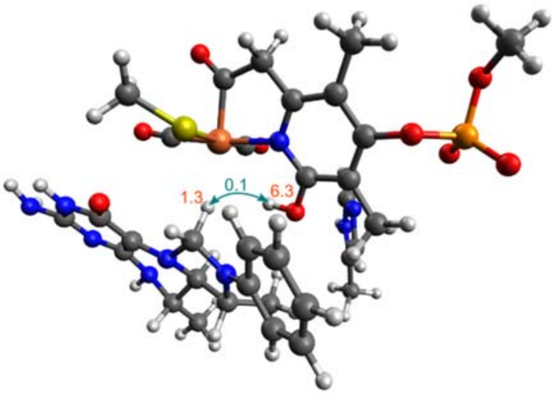


| | | A6(OHtoH-$CH_2$) | | E6(OHtoH-$CH_2$) |
|---|---|---|---|---|
| **Δ*E*** | | 0.0 | | 0.0 |
| ***δ*** | OH | 7.05 | OH | 6.29 |
| | $CH_2$ (pro-R) | 4.06 | $CH_2$ (pro-R) | 1.32 |
| | $CH_2$ (pro-S) | 5.94 | $CH_2$ (pro-S) | 5.90 |
| ***J*** | OH/$CH_2$ (pro-R) | 0.58 | OH/$CH_2$ (pro-R) | 0.14 |
| | OH/$CH_2$ (pro-S) | 0.18 | OH/$CH_2$ (pro-S) | -0.07 |
| | $CH_2$/$CH_2$ | -10.42 | $CH_2$/$CH_2$ | -10.00 |

**Table S9 (continued)**

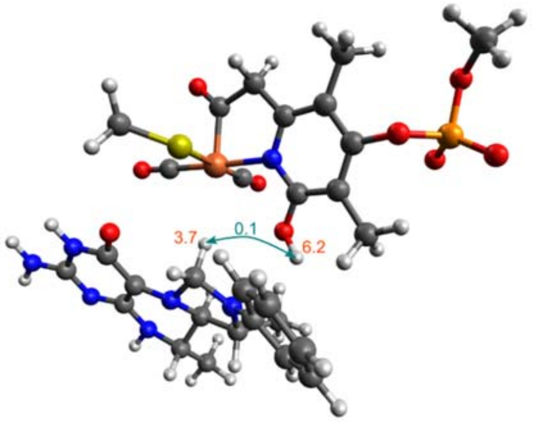


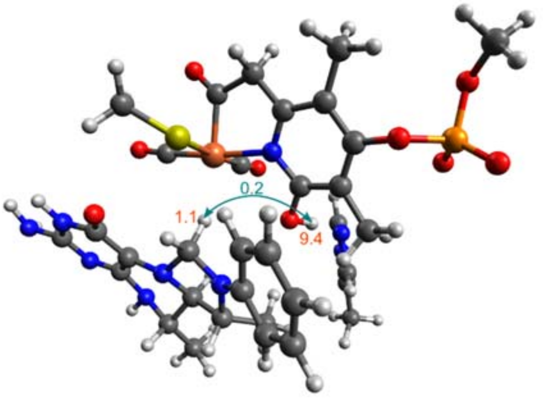


| | A7(OHaway-$CH_2$) | | E7(OHaway-$CH_2$) | |
|---|---|---|---|---|
| **Δ*E*** | | 0.0 | | -3.0 |
| ***δ*** | OH | 6.15 | OH | 9.44 |
| | $CH_2$ (pro-R) | 3.74 | $CH_2$ (pro-R) | 1.08 |
| | $CH_2$ (pro-S) | 5.57 | $CH_2$ (pro-S) | 5.73 |
| ***J*** | OH/$CH_2$ (pro-R) | 0.12 | OH/$CH_2$ (pro-R) | 0.23 |
| | OH/$CH_2$ (pro-S) | 0.08 | OH/$CH_2$ (pro-S) | 0.05 |
| | $CH_2$/$CH_2$ | -8.03 | $CH_2$/$CH_2$ | -7.61 |

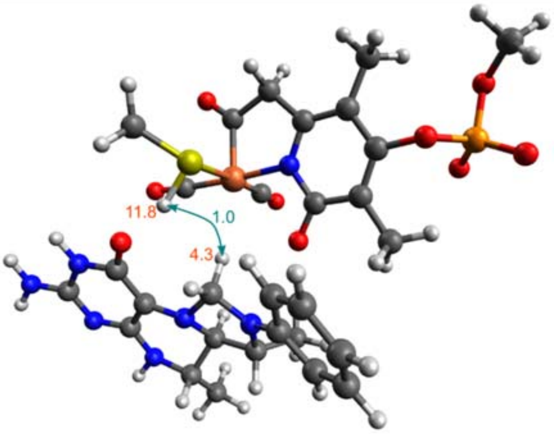


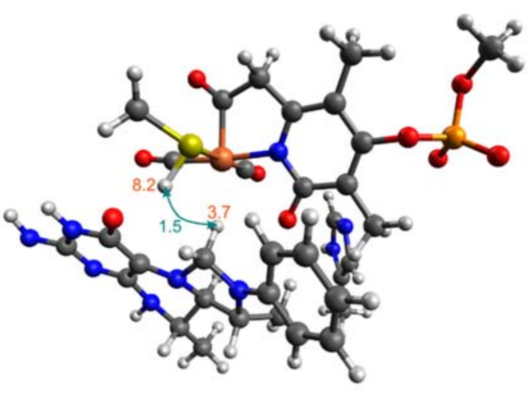


| | A8(SHtoO-$CH_2$) | | E8(SHtoO-$CH_2$) | |
|---|---|---|---|---|
| **Δ*E*** | | 20.6 | | 23.0 |
| ***δ*** | SH | 11.81 | SH | 8.18 |
| | $CH_2$ (pro-R) | 4.27 | $CH_2$ (pro-R) | 3.67 |
| | $CH_2$ (pro-S) | 4.83 | $CH_2$ (pro-S) | 5.00 |
| ***J*** | SH/$CH_2$ (pro-R) | 0.96 | SH/$CH_2$ (pro-R) | 1.52 |
| | SH/$CH_2$ (pro-S) | 0.43 | SH/$CH_2$ (pro-S) | 0.25 |
| | $CH_2$/$CH_2$ | -5.22 | $CH_2$/$CH_2$ | -3.62 |

**Table S9 (continued)**

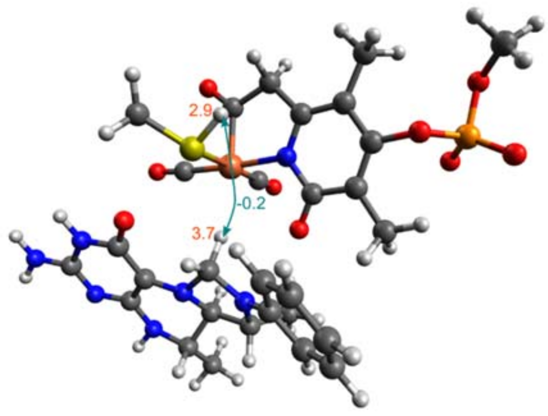


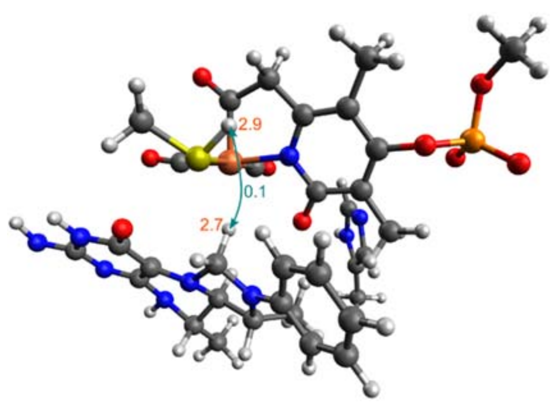


| | | A9(SHaway-$CH_2$) | | E9(SHaway-$CH_2$) |
|---|---|---|---|---|
| **Δ*E*** | | 20.5 | | 21.7 |
| ***δ*** | SH | 2.86 | SH | 2.94 |
| | $CH_2$ (pro-R) | 3.70 | $CH_2$ (pro-R) | 2.74 |
| | $CH_2$ (pro-S) | 5.81 | $CH_2$ (pro-S) | 5.65 |
| ***J*** | SH/$CH_2$ (pro-R) | -0.15 | SH/$CH_2$ (pro-R) | 0.12 |
| | SH/$CH_2$ (pro-S) | -0.23 | SH/$CH_2$ (pro-S) | -0.29 |
| | $CH_2$/$CH_2$ | -7.40 | $CH_2$/$CH_2$ | -6.14 |

**Table S10.**

Labels of the "3H" optimized models of the [Fe]-hydrogenase active site. Successfully optimized models are marked with "yes". The notation "→X" means that the given model converged instead to structure X during optimization. See section S2.10 for details.

| | Label | C | G | Corresponding structure in Fig. 3 of main article |
|---|---|---|---|---|
| 1 | OHtoH-$FeH_2$ | yes | →G2 | **2** |
| 2 | OHaway-$FeH_2$ | yes | yes | **2** |
| 3 | OHtoH-FeH-SHtoO | yes | yes | |
| 4 | OHtoH-FeH-SHaway | yes | yes | |
| 5 | OHaway-FeH-SHtoO | yes | yes | |
| 6 | OHaway-FeH-SHaway | yes | yes | |
| 7 | OHtoH-$CH_2$-SHtoO | yes | yes | |
| 8 | OHtoH-$CH_2$-SHaway | yes | yes | |
| 9 | OHaway-$CH_2$-SHtoO | yes | | |
| 10 | OHaway-$CH_2$-SHaway | yes | | |
| | Source | Ref. (*16*), fig. 9 | Ref. (*10*), PDB: 6hav | |

**Table S11.**

[1]H chemical shifts (in ppm) and couplings (in Hz) computed for different structures with three hydrogens included in the active site. Results shown in the right column (**G2-G8**) are obtained from QM/MM optimized models derived from the closed-state crystal structure (PDB: 6HAV) published in ref. (*10*). Results shown in the left column (**C1-C10**) are obtained from the QM/MM optimized MD-snapshot presented in Fig. 9 of ref. (*16*). For an overview of the models with two hydrogens in the active site, see Table S10. Chemical shifts were computed at the TPSS/pS2 and *J*-couplings – at the PBE0/pJ2 level, both with electrostatic embedding. The relative energy with respect to **C/G2**(OHaway-FeH$_2$) for models **C/G**, respectively, at the TPSS-D3BJ/def2-TZVP/MM level is given in kcal·mol$^{-1}$. In the figures, shifts are shown in orange and couplings in teal.

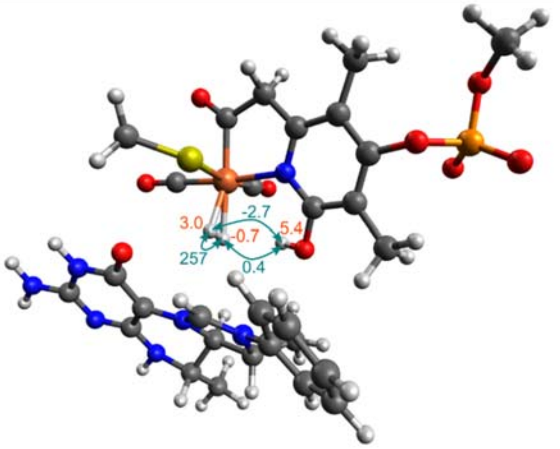


**C1(OHtoH-FeH$_2$)**

| | | |
|---|---|---|
| **Δ*E*** | | 3.3 |
| ***δ*** | OH | 5.42 |
| | Fe-H$_2$ (δ+) | 2.97 |
| | Fe-H$_2$ (δ-) | -0.75 |
| | CH (pro-S) | 9.14 |
| ***J*** | Fe-H$_2$/Fe-H$_2$ | 256.7 |
| | OH/Fe-H$_2$ (δ+) | -2.71 |
| | OH/Fe-H$_2$ (δ-) | 0.37 |
| | CH/Fe-H$_2$ (δ+) | 0.37 |
| | CH/Fe-H$_2$ (δ-) | 1.65 |
| | OH/CH | 0.09 |

**Table S11 (continued)**

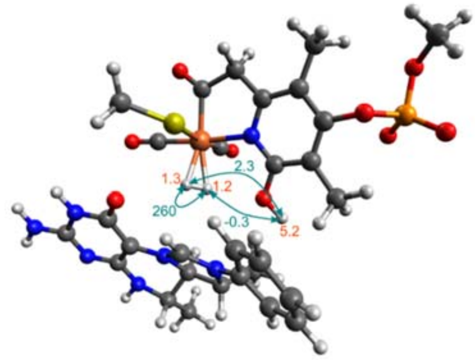


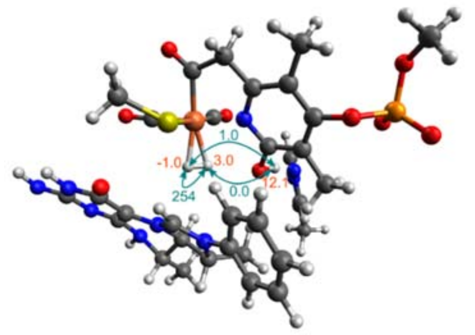


| | C2(OHaway-FeH$_2$) | | G2(OHaway-FeH$_2$) | |
|---|---|---|---|---|
| **Δ*E*** | | 0.0 | | 0.0 |
| ***δ*** | OH | 5.15 | OH | 12.11 |
| | Fe-H$_2$ (δ+) | 0.94 | Fe-H$_2$ (δ-) | -0.98 |
| | Fe-H$_2$ (δ-) | 0.96 | Fe-H$_2$ (δ+) | 3.04 |
| | CH (pro-S) | 9.32 | CH (pro-S) | 10.14 |
| ***J*** | Fe-H$_2$/Fe-H$_2$ | 260.5 | Fe-H$_2$/Fe-H$_2$ | 253.6 |
| | OH/Fe-H$_2$ (δ+) | 2.39 | OH/Fe-H$_2$ (δ-) | 0.99 |
| | OH/Fe-H$_2$ (δ-) | -0.31 | OH/Fe-H$_2$ (δ+) | 0.03 |
| | CH/Fe-H$_2$ (δ+) | 0.28 | CH/Fe-H$_2$ (δ-) | 0.05 |
| | CH/Fe-H$_2$ (δ-) | 1.88 | CH/Fe-H$_2$ (δ+) | 2.43 |
| | OH/CH | 0.06 | OH/CH | 0.00 |

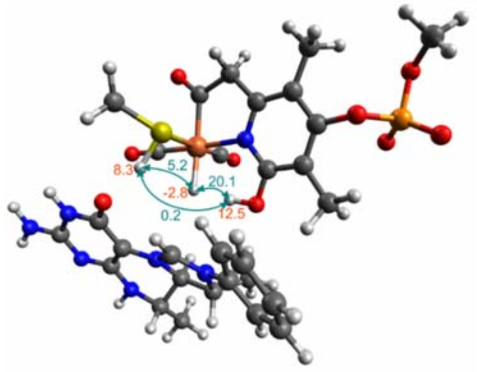


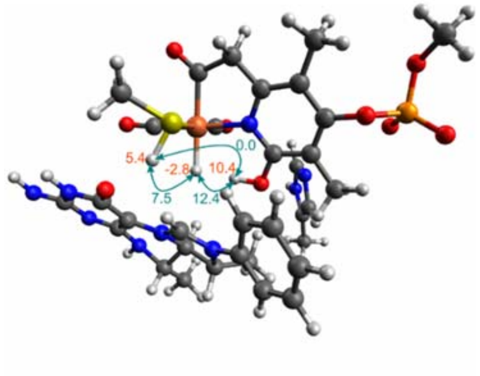


| | C3(OHtoH-FeH-SHtoO) | | G3(OHtoH-FeH-SHtoO) | |
|---|---|---|---|---|
| **Δ*E*** | | 4.9 | | 8.5 |
| ***δ*** | Fe-H | -2.76 | Fe-H | -2.76 |
| | OH | 12.48 | OH | 10.43 |
| | SH | 8.26 | SH | 5.43 |
| | CH (pro-S) | 8.80 | CH (pro-S) | 8.64 |
| ***J*** | OH/Fe-H | 20.09 | OH/Fe-H | 12.40 |
| | SH/Fe-H | 5.17 | SH/Fe-H | 7.45 |
| | CH/Fe-H | 0.39 | CH/Fe-H | 0.03 |
| | OH/SH | 0.19 | OH/SH | 0.03 |
| | SH/CH | 0.48 | SH/CH | 0.72 |
| | OH/CH | 0.32 | OH/CH | 0.28 |

**Table S11 (continued)**

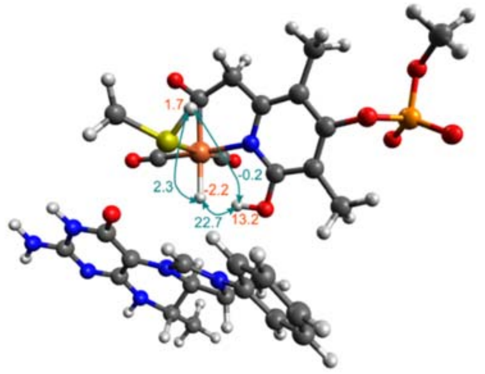


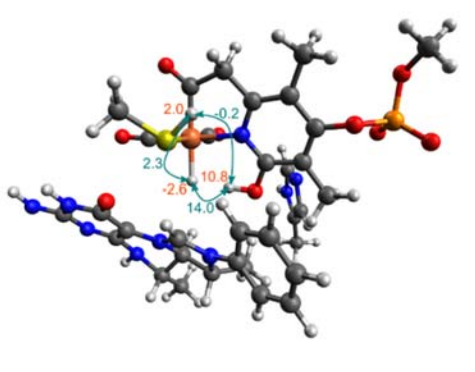


| | | C4(OHtoH-FeH-SHaway) | | G4(OHtoH-FeH-SHaway) |
|---|---|---|---|---|
| **Δ*E*** | | 5.9 | | 7.3 |
| ***δ*** | Fe-H | -2.36 | Fe-H | -2.56 |
| | OH | 13.15 | OH | 10.79 |
| | SH | 1.83 | SH | 2.01 |
| | CH (pro-S) | 9.51 | CH (pro-S) | 9.39 |
| ***J*** | OH/Fe-H | 22.92 | OH/Fe-H | 14.00 |
| | SH/Fe-H | 1.80 | SH/Fe-H | 2.31 |
| | CH/Fe-H | -0.07 | CH/Fe-H | -0.48 |
| | OH/SH | -0.18 | OH/SH | -0.24 |
| | SH/CH | -0.23 | SH/CH | -0.35 |
| | OH/CH | 0.39 | OH/CH | 0.33 |

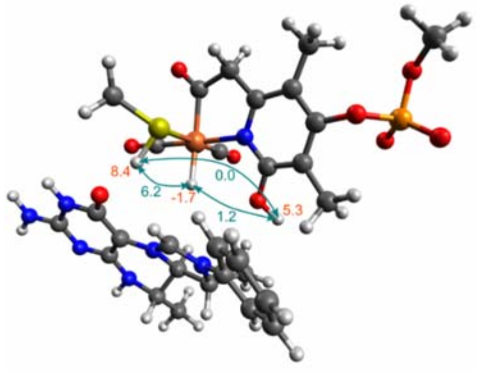


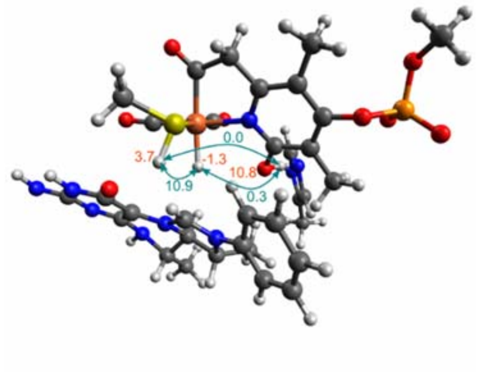


| | | C5(OHaway-FeH-SHtoO) | | G5(OHaway-FeH-SHtoO) |
|---|---|---|---|---|
| **Δ*E*** | | 14.9 | | 9.4 |
| ***δ*** | Fe-H | -1.68 | Fe-H | -1.26 |
| | OH | 5.28 | OH | 10.83 |
| | SH | 8.37 | SH | 3.73 |
| | CH (pro-S) | 8.89 | CH (pro-S) | 8.49 |
| ***J*** | OH/Fe-H | 1.25 | OH/Fe-H | 0.32 |
| | SH/Fe-H | 6.19 | SH/Fe-H | 10.93 |
| | CH/Fe-H | -1.97 | CH/Fe-H | -3.62 |
| | OH/SH | 0.00 | OH/SH | 0.00 |
| | SH/CH | 0.61 | SH/CH | 1.22 |
| | OH/CH | 0.06 | OH/CH | 0.00 |

**Table S11 (continued)**

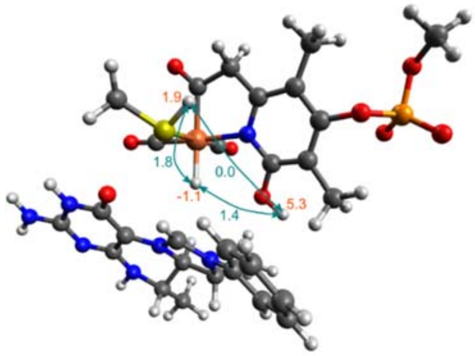


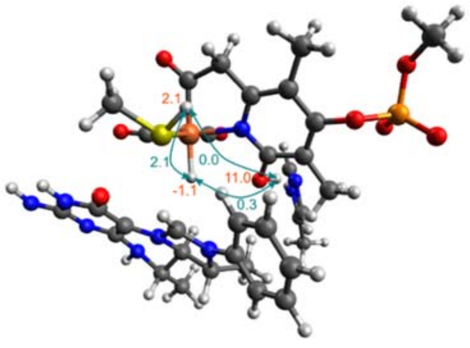


| | | C6(OHaway-FeH-SHaway) | | G6(OHaway-FeH-SHaway) |
|---|---|---|---|---|
| **Δ*E*** | | 16.6 | | 8.7 |
| ***δ*** | Fe-H | -1.13 | Fe-H | -1.12 |
| | OH | 5.33 | OH | 11.04 |
| | SH | 1.86 | SH | 2.06 |
| | CH (pro-S) | 9.81 | CH (pro-S) | 9.19 |
| ***J*** | OH/Fe-H | 1.38 | OH/Fe-H | 0.32 |
| | SH/Fe-H | 1.78 | SH/Fe-H | 2.07 |
| | CH/Fe-H | -4.37 | CH/Fe-H | -4.90 |
| | OH/SH | 0.00 | OH/SH | 0.00 |
| | SH/CH | -0.25 | SH/CH | -0.41 |
| | OH/CH | 0.07 | OH/CH | 0.00 |

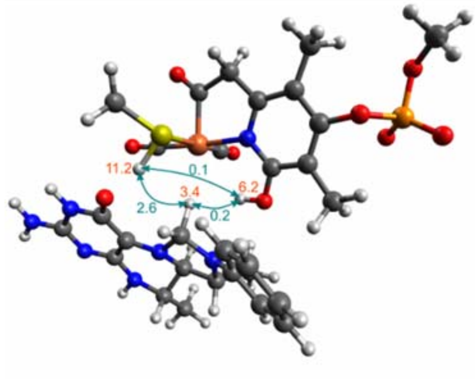


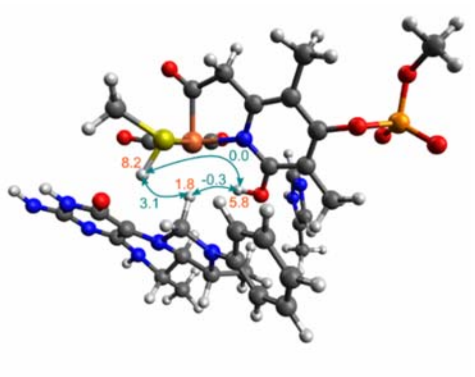


| | | C7(OHtoH-$CH_2$-SHtoO) | | G7(OHtoH-$CH_2$-SHtoO) |
|---|---|---|---|---|
| **Δ*E*** | | 10.7 | | 7.8 |
| ***δ*** | OH | 6.20 | OH | 5.84 |
| | SH | 11.20 | SH | 8.17 |
| | $CH_2$ (pro-R) | 3.39 | $CH_2$ (pro-R) | 1.84 |
| | $CH_2$ (pro-S) | 6.03 | $CH_2$ (pro-S) | 5.89 |
| ***J*** | OH/$CH_2$ (pro-R) | 0.21 | OH/$CH_2$ (pro-R) | -0.27 |
| | OH/$CH_2$ (pro-S) | -0.24 | OH/$CH_2$ (pro-S) | -0.44 |
| | SH/$CH_2$ (pro-R) | 2.65 | SH/$CH_2$ (pro-R) | 3.13 |
| | SH/$CH_2$ (pro-S) | 1.01 | SH/$CH_2$ (pro-S) | 1.06 |
| | OH/SH | 0.08 | OH/SH | -0.03 |
| | $CH_2$/$CH_2$ | -12.10 | $CH_2$/$CH_2$ | -10.82 |

**Table S11 (continued)**

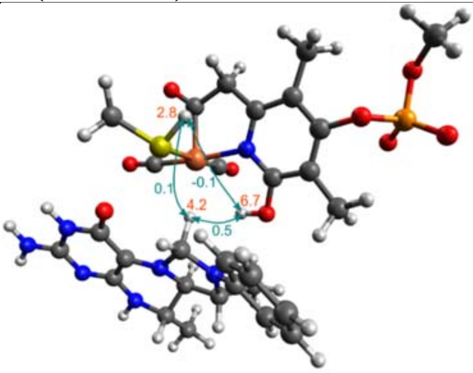


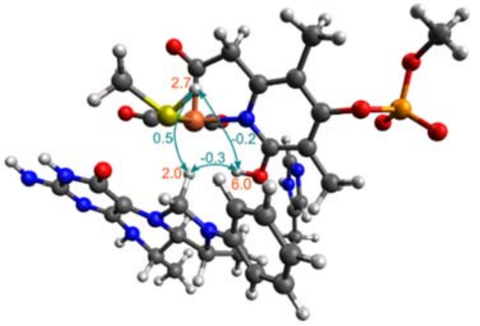


| | | C8(OHtoH-$CH_2$-SHaway) | | G8(OHtoH-$CH_2$-SHaway) |
|---|---|---|---|---|
| **Δ*E*** | | 8.9 | | 7.6 |
| ***δ*** | OH | 6.69 | OH | 6.02 |
| | SH | 2.81 | SH | 2.73 |
| | $CH_2$ (pro-R) | 4.18 | $CH_2$ (pro-R) | 2.04 |
| | $CH_2$ (pro-S) | 7.36 | $CH_2$ (pro-S) | 6.81 |
| ***J*** | OH/$CH_2$ (pro-R) | 0.54 | OH/$CH_2$ (pro-R) | -0.26 |
| | OH/$CH_2$ (pro-S) | -0.15 | OH/$CH_2$ (pro-S) | -0.41 |
| | SH/$CH_2$ (pro-R) | 0.11 | SH/$CH_2$ (pro-R) | 0.52 |
| | SH/$CH_2$ (pro-S) | -0.15 | SH/$CH_2$ (pro-S) | -0.26 |
| | OH/SH | -0.13 | OH/SH | -0.18 |
| | $CH_2$/$CH_2$ | -15.20 | $CH_2$/$CH_2$ | -14.09 |

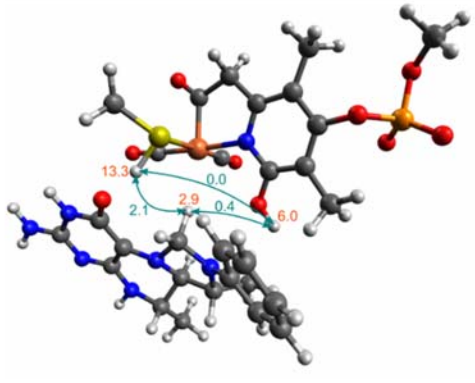


| | | C9(OHaway-$CH_2$-SHtoO) |
|---|---|---|
| **Δ*E*** | | 6.5 |
| ***δ*** | OH | 6.03 |
| | SH | 13.33 |
| | $CH_2$ (pro-R) | 2.90 |
| | $CH_2$ (pro-S) | 5.33 |
| ***J*** | OH/$CH_2$ (pro-R) | 0.40 |
| | OH/$CH_2$ (pro-S) | 0.22 |
| | SH/$CH_2$ (pro-R) | 2.06 |
| | SH/$CH_2$ (pro-S) | 0.76 |
| | OH/SH | 0.00 |
| | $CH_2$/$CH_2$ | -7.64 |

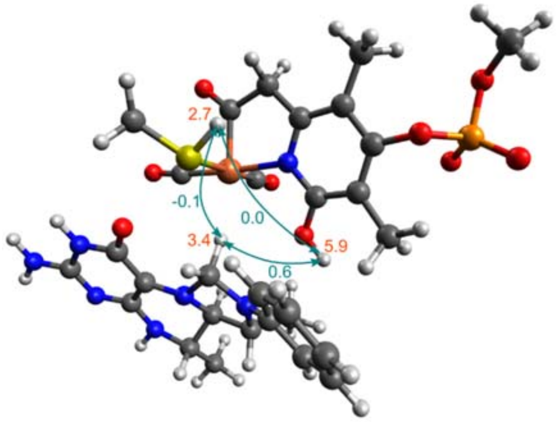


| | C10(OHaway-$CH_2$-SHaway) | |
|---|---|---|
| ***ΔE*** | | 7.3 |
| ***δ*** | OH | 5.91 |
| | SH | 2.71 |
| | $CH_2$ (pro-R) | 3.37 |
| | $CH_2$ (pro-S) | 6.76 |
| ***J*** | OH/$CH_2$ (pro-R) | 0.59 |
| | OH/$CH_2$ (pro-S) | 0.26 |
| | SH/$CH_2$ (pro-R) | -0.10 |
| | SH/$CH_2$ (pro-S) | -0.22 |
| | OH/SH | 0.00 |
| | $CH_2$/$CH_2$ | -10.25 |

**Table S12.**

$^1H$ chemical shifts computed for model **C2**(OHaway-FeH2) at the TPSS/pS2 level with MM embedding and deviations when using different basis sets or functionals. All values are in ppm.

| Nucleus | $\delta$ (TPSS/pS2) | $\Delta\delta$ (pS2-pS1)[a] | $\Delta\delta$ (pS3-pS2)[a] | $\Delta\delta$ (r$^2$SCAN-TPSS)[b] |
|---|---|---|---|---|
| Fe-$H_2$ | 1.07 | -0.00 | -0.02 | -0.07 |
| | 0.91 | 0.02 | 0.12 | 0.04 |
| OH | 5.21 | 0.26 | 0.25 | 0.27 |
| CH (pro-S) | 9.31 | 0.32 | 0.17 | -0.13 |

a Using the TPSS functional.
b Using the pS3 basis set.

**Table S13.**

$^1$H chemical shifts computed for model **C1**(OHtoH-$FeH_2$) at the TPSS/pS2 level with MM embedding and deviations when using a CPCM embedding with ε=4.0 instead. All values are in ppm.

| Nucleus | *δ* (QM/MM) | Δ*δ* (CPCM-QM/MM) |
|---|---:|---:|
| Fe-$H_2$ (δ+) | 2.97 | -0.20 |
| Fe-$H_2$ (δ-) | -0.75 | 0.00 |
| OH | 5.42 | -0.36 |
| CH (pro-S) | 9.14 | -0.08 |

**Table S14.**

[1]H coupling constants computed for model **C2**(OHaway-FeH$_2$) at the PBE0/pJ2 level with MM embedding and deviations when using different basis sets or functionals. All values are in Hz.

| **Coupling** | ***J* (PBE0/pJ2)** | **Δ*J* (pJ2-pJ1)[a]** | **Δ*J* (pJ3-pJ2)[a]** | **Δ*J* (PBE0-PBE)[b]** | **Δ*J* (big-small)[c]** |
|---|---|---|---|---|---|
| Fe-H$_2$/Fe-H$_2$ | 260.21 | 5.32 | 1.93 | 10.77 | 0.20 |
| OH/Fe-H$_2$ | -0.38 | -0.18 | -0.08 | -0.06 | 0.13 |
| | 2.47 | -0.04 | -0.03 | -0.10 | 0.13 |
| CH/Fe-H$_2$ | 1.86 | -0.31 | -0.11 | 0.08 | 0.26 |
| | 0.33 | -0.45 | -0.17 | -0.05 | 0.26 |
| OH/CH | 0.06 | -0.03 | -0.02 | 0.00 | 0.15 |

a Using the PBE0 functional.

b Using the pJ3 basis set.

c Using the PBE functional, pJ2 on the QM atoms and pcseg-1 on the “extended” QM region.

**Table S15.**

[1]H coupling constants computed for model **C1**(OHtoH-$FeH_2$) and at the PBE0/pJ2 level with MM embedding and deviations when using a CPCM embedding with ε=4.0 instead. All values are in Hz.

| **Coupling** | ***J* (QM/MM)** | **Δ*J* (CPCM-QM/MM)** |
|---|---|---|
| Fe-$H_2$/Fe-$H_2$ | 256.72 | -0.20 |
| OH/Fe-$H_2$ (δ+) | -2.71 | 0.10 |
| OH/Fe-$H_2$ (δ-) | 0.37 | 0.09 |
| CH/Fe-$H_2$ (δ+) | 0.37 | -0.02 |
| CH/Fe-$H_2$ (δ-) | 1.65 | -0.11 |
| OH/CH | 0.09 | 0.00 |
